# The genome of stress tolerant crop wild relative *Paspalum vaginatum* leads to increased biomass productivity in the crop *Zea mays*

**DOI:** 10.1101/2021.08.18.456832

**Authors:** Guangchao Sun, Nishikant Wase, Shengqiang Shu, Jerry Jenkins, Bangjun Zhou, Cindy Chen, Laura Sandor, Chris Plott, Yuko Yoshinga, Christopher Daum, Peng Qi, Kerrie Barry, Anna Lipzen, Luke Berry, Thomas Gottilla, Ashley Foltz, Huihui Yu, Ronan O’Malley, Chi Zhang, Katrien M. Devos, Brandi Sigmon, Bin Yu, Toshihiro Obata, Jeremy Schmutz, James C. Schnable

**Affiliations:** Quantitative Life Sciences Initiative, University of Nebraska-Lincoln, Lincoln, NE, 68588 USA; Center for Plant Science Innovation, University of Nebraska-Lincoln, Lincoln, NE, 68588 USA; Department of Agronomy and Horticulture, University of Nebraska-Lincoln, Lincoln, NE, 68588 USA; Department of Biochemistry, University of Nebraska-Lincoln, Lincoln, NE, 68588 USA; Department of Energy Joint Genome Institute, Lawrence Berkeley National Laboratory, CA 94598, USA.; HudsonAlpha Institute for Biotechnology, Huntsville, AL 35806, USA; School of Biological Sciences, University of Nebraska-Lincoln, Lincoln, NE, 68588 USA; Institute of Plant Breeding, Genetics and Genomics, Department of Crop and Soil Sciences, University of Georgia, Athens, GA 30602, USA; Department of Crop and Soil Sciences, University of Georgia, Athens, GA 30602, USA; Department of Plant Biology, University of Georgia, Athens, GA 30602, USA; Department of Plant Pathology, University of Nebraska-Lincoln, Lincoln, NE, 68588 USA

## Abstract

A number of crop wild relatives can tolerate extreme stressed to a degree outside the range observed in their domesticated relatives. However, it is unclear whether or how the molecular mechanisms employed by these species can be translated to domesticated crops. Paspalum *Paspalum vaginatum* is a self-incompatible and multiply stress-tolerant wild relative of maize and sorghum. Here we describe the sequencing and pseudomolecule level assembly of a vegetatively propagated accession of *P. vaginatum*. Phylogenetic analysis based on 6,151 single-copy syntenic orthologous conserved in 6 related grass species placed paspalum as an outgroup of the maize-sorghum clade demonstrating paspalum as their closest sequenced wild relative. In parallel metabolic experiments, paspalum, but neither maize nor sorghum, exhibited significant increases in trehalose when grown under nutrient-deficit conditions. Inducing trehalose accumulation in maize, imitating the metabolic phenotype of paspalum, resulting in autophagy dependent increases in biomass accumulation.

## Introduction

Domesticated crops from the grass family provide, directly or indirectly, the majority of total calories consumed by humans around the globe. Among the domesticated grasses the yields of three crops dramatically increased as part of the green revolution: rice, wheat and maize. These yield increases resulted from both breeding and greater availability and application of fertilizer. From 1960 to 2014, the amount of nitrogen (N) and phosphorus (P) fertilizer applied worldwide increased nine-fold and five-fold respectively^1–3^. Today these three crops account for approximately one half of total harvested staple crop area and total global calorie production as well as greater than one half of total global fertilizer consumption. Manufacturing N fertilizer is an energy intensive process^4^ and the production of P from mineral sources may peak as early as 2030^5^. Fertilizer costs are often the second largest variable input after seed in rain fed agricultural systems. In the United States Corn Belt alone, 5.6 million tons of N and 2.0 million tons of P have been applied annually to maize (*Zea mays*) fields since 2010^6^. In the 2015 growing season, these fertilizers accounted for an estimated $5 billion in input costs^7, 8^. Fertilizer runoff resulting from inefficient uptake or over application can result in damage to both aquatic ecosystems and drinking water quality^9–12^.

Improving the productivity of crop plants per unit of fertilizer applied would increase the profitability of agriculture while decreasing its environmental impact^13–15^. A significant portion of the overall increase in maize yields appears to be explained by selection for increased stress tolerance and yield stability in the decades since the 1930s^16, 17^. The observations from maize suggest it may be possible to increase the stress tolerance and resource-use efficiency of crops in a manner that is either neutral or beneficial to overall yield potential. Some crop wild relatives exhibit degrees of stress tolerance well outside the range observed in their domesticated relatives, and therefore may employ mechanisms not present in the primary germplasm of crops^14^.

*Paspalum vaginatum* (seashore paspalum – or simply paspalum) is a relative of maize (*Zea mays*) and sorghum (*Sorghum bicolor*). It is currently found on saltwater beaches and in other regions of high salinity around the globe^18, 19^. Reports suggest that paspalum is tolerant of drought^20–23^, cold stress^24–26^, low light^27^, and crude oil contamination^28^. Paspalum grows primarily in the wild, but breeding efforts have led to the development of turfgrass cultivars for use in areas with high soil salinity, limited access to freshwater, or where turf is irrigated with wastewater^27, 29^. Paspalum requires less N to maintain visible health than other grasses employed as turfgrasses in environments where paspalum thrives^29^. Historically few genetic resources have been available for this species, although a set of genetic maps were recently published^30^. The paucity of genetic and genomic investigations may in part result from the challenging reproductive biology of this species; paspalum is self-incompatible and is primarily propagated as heterozygous vegetative clones^29, 31^.

Here, we generate a pseudomolecule level genome assembly for a reference genotype of paspalum (PI 509022), enabling comparative transcriptomic and genomics analysis. Phylogenetic analyses employing syntenic gene copies confirmed paspalum’s placement as a close outgroup to maize and sorghum and the paspalum genome exhibits a high degree of conserved collinearity with that of sorghum. The genes involved in telomere maintenance and DNA repair have experience significant copy number expansion in the paspalum lineage as do several gene families which transcriptionally respond to nitrogen or phosphorous deficit stress. Changes in trehalose accumulation and the expression of genes involved in trehalose metabolism were observed in response to multiple nutrient-deficit stresses were observed in paspalum, but not in paired datasets collected from sorghum and maize under identical conditions. Replicating the pattern of trehalose metabolism observed in wild-type paspalum by inhibiting the enzyme responsible for degrading trehalose increased trehalose accumulation, biomass accumulation and shoot-to-root ratios in maize under nutrient-optimal and -deficient conditions. The induced accumulation of trehalose in maize was associated with lipidation of AUTOPHAGY-RELATED8 (ATG8) a marker for autophagy activity. Treatment with a chemical inhibitor for autophagy abolished the increased biomass accumulation observed in maize plants accumulating additional trehalose, suggesting that increased autophagy as a potential mechanism for the observed increased productivity observed in maize plants accumulating additional trehalose.

## Results

### Characteristics of the paspalum genome

We generated 5,021,142 PacBio reads with a median length of 9,523 bp from genomic DNA isolated from dark-treated tissue of the heterozygous paspalum clone PI 509022. The reads were assembled into 1,903 main genome scaffolds with an N50 of 44.5 Mbp and a total length of 651.0 Mbp (Table S1). This is modestly larger than the estimated haploid gene size of the paspalum genome of 593 Mbp (See Methods and Supplementary Note 1)^32^. Flow cytometry carried out within this study confirmed that the genome size of the paspalum clone employed in this study was approximately 590 Mbp (Figure S1A & B). This modest over-assembly may represent haplotype-specific sequences which is supported by the binomial distribution of read coverage mapped to the current genome assembly (Figure S1C). We used published sequence data from markers genetically mapped in an F1 population generated from a cross between two heterozygous paspalum individuals^30^ to integrate a set of 347 scaffolds into ten pseudomolecules spanning >82% of the estimated total haploid paspalum genome (Supplementary Note 2). Scaffolds that were not anchored in a chromosome were classified into bins depending on sequence content. Contamination was identified using BLASTN against the NCBI nucleotide collection (NR/NT) and BLASTX using a set of known microbial proteins. Additional scaffolds were classified as repetitive (>95% masked with 24 mers that occur more than 4 times in the genome) (197 scaffolds, 12.4 Mb), alternative haplotypes (unanchored sequence with >95% identity and >95% coverage within a chromosome) (3,276 scaffolds, 187.9 Mb), and low quality (>50% unpolished bases, post polishing) (9 scaffolds, 204.5 Kb) (Table S1). A set of 45,843 gene models were identified and annotated using a combination of approaches (See Methods). A total of 22,148 syntenic orthologous gene pairs were identified between the paspalum and sorghum genomes (Figure S2A & B). The large inversions observed on chromosome 4 and chromosome 7 were previously validated by a study a genetic map was constructed using GBS genotyping technology^30^. Small translocations were also observed between chromosome 8 and chromosome 4 (Figure S2 A& B). The predicted protein sequences of annotated paspalum genes tended to cover the full length of the most closely related protein in sorghum, and vice versa, indicating most annotated gene models in the paspalum genome assembly are likely full length (Figure S2C). On a macro level, the paspalum genome displays many features common to other grass genomes: a higher gene density in the distal chromosome regions than pericentromeric regions and, conversely, a higher frequency of transposable elements and other repetitive sequences in pericentromeric regions than distal chromosome regions, and syntenic evidence of the pre-grass (rho) whole genome duplication (Figure 1 A).

**Figure 1.**
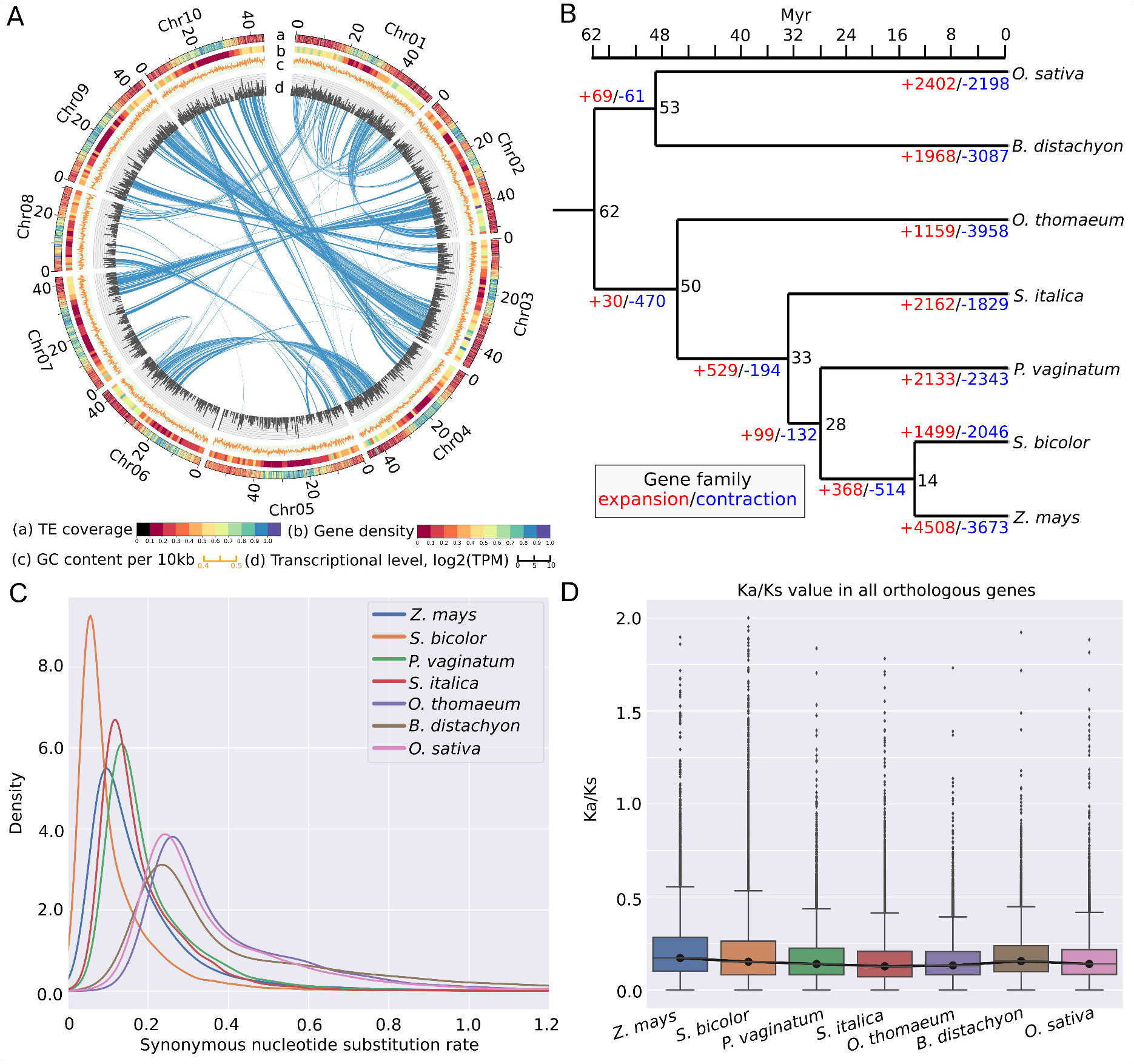
Paspalum (*Paspalum vaginatum*) genome and evolution. (A) Circos plot for the paspalum genome. a: TE coverage per 100 Kb, b:GC content per 10 kb region, c: gene density (generic region coverage) per 1 Mb, d: transcription represented by log_2_(TPM) per 100 kb and e: inter- and intra- chromosomal synteny. (B) Phylogeny and estimated divergence times among maize (*Zea mays*), sorghum, paspalum, foxtail millet (*Setaria italica*), *Oropetium* (*Oropetium thomaeum*), *Brachypodium* (*Brachypodium distachyon*), and rice (*Oryza sativa*). Numbers in black indicate the estimated divergence time (in millions of years before present) for each node. Numbers in blue and red indicate the number of gene families predicted to have experienced copy number expansion or contraction along each branch of the phylogeny, respectively. (C) Distribution of the estimated lineage-specific synonymous substitution rates for syntenically conserved genes in each of the seven species shown in panel A (see Methods) (D) Distribution of the estimated lineage-specific ratios of nonsynonymous substitution rates to synonymous substitution rates for syntenically conserved genes among each of the seven species shown in panel A.

### Comparative genomics analysis of paspalum and its relatives

Paspalum belongs to the grass tribe Paspaleae, a group which, together with the Andropogoneae (which includes maize and sorghum) – and the Arundinelleae, forms a clade sister to the Paniceae – which includes foxtail millet (*Setaria italica*). Paspaleae,Andropogoneae and Paniceae are all members of the grass subfamily Panicoideae, while *Oropetium* belongs to the grass clade Cynodonteae^33^. We constructed phylogenic trees using DNA alignments for 6,151 single-copy syntenic orthologous genes present in *Zea mays*, *Sorghum bicolor*, *Setaria italica*, *Oropetium thomaeum*, *Brachypodium distachyon*, *Oryza sativa*, and *Paspalum vaginatum*. A total of 5,859 trees representing 49 unique topologies and placing *B. distachyon* and *O. sativa* in a monophyletic outgroup survived quality filtering (see Methods for filtering criteria). The most common topology among these trees – represented by 4,265 individual gene trees (73%) – was consistent with the previous consensus placement of paspalum (Figure S3A & Figure 1B). The second most common topology, represented by 762 individual gene trees (13%), placed paspalum sister to foxtail millet (Figure S3A).

Dating placed the split of the Chloridoideae (represented by *Oropetium thomaeum*) from the Panicoideae at 50 million years before present and indicated that, within the Panicoideae, the Paniceae shared a common ancestor with the Andropogoneae/Paspaleae clade (represented by paspalum, sorghum, and maize) at 33 million years (Myr) before present, a date modestly earlier than previous estimates ( 26 Myr ago)^34, 35^ (Figure 1B). The divergence of the lineage leading to paspalum from that leading to maize and sorghum – (the split between the Andropogoneae and Paspaleae) – was estimated to have occurred approximately 28 million years before present. We calculated branch specific synonymous (Ks) and nonsynonymous (Ka) nucleotide substitution rates for syntenic orthologous gene groups based on known species relationships (Figure 1C; Supplementary note 3). Consistent with previous reports, maize exhibited greater modal synonymous substitution rates than sorghum, even though these are sister lineages in the phylogeny^36^ (Figure 1C). The modal synonymous substitution rates in paspalum were modestly higher than those observed in foxtail millet (Figure 1D).

Annotated protein sequences for sorghum, foxtail millet, *Oropetium*, *Brachypodium* (*Brachypodium distachyon*), and paspalum were grouped into 25,675 gene families. Of these families 16,038 were represented by at least one gene copy in each of the five species, with the remainder being present in 1-4 species (Figure S3B). A set of 721 gene families were unique to paspalum. This number was modestly less than the number of species-specific gene families identified in brachypodium and modestly more than the number of species-specific gene families observed in sorghum and foxtail millet (Figure S3B). Of the 21,091 gene families present in paspalum, 75% (15,769) were represented by only a single copy in the paspalum genome and 17 % (3,524) were represented by two copies. These values are similar to those observed in sorghum and foxtail millet which shared the same most recent common pre-grass (rho) whole genome duplication (Figure S3C). A set of 149 gene families were identified as undergoing copy number expansion with a significantly different evolution rate (lambda) in the paspalum lineage. These included families of genes annotated with gene ontology (GO) terms related to homeostatic processes such as telomere maintenance and DNA repair, protein modification, stress response, nutrient reservoir activity and oxidation-reduction process (Supplementary note 4).

### General and paspalum-specific physiological responses to nutrient-deficit stress

Paspalum requires little fertilizer in order to remain visibly healthy, however it was unclear whether paspalum is actually more efficient at producing biomass under nitrogen limited conditions. A comparison was conducted of growth of paspalum, sorghum, and maize plants under ideal, nitrogen limited and phosphorous limited conditions. Visible effects were apparent in maize and sorghum seedlings under N- or P-deficient conditions but not in clonally propagated paspalum plants (ramets) three weeks after planting (Figure 2A). Both maize and sorghum exhibited significant decreases in above ground fresh biomass accumulation when grown under N- or P-deficient conditions whereas paspalum did not (Figure 2B). This result should be interpreted with the caveat that paspalum accumulated the least biomass per plant of the three species under nutrient-optimal conditions. In both maize and sorghum, N-deficit was associated with significant increases in root length. However, a statistically significant increase in root length in response to P-deficit stress was only observed for sorghum (Figure 2C). Paspalum ramets grown under N-deficient conditions showed a modest but statistically significant increase in root length compared to optimal nutrient conditions, while P-deficit stress did not produce any statistically significant increase in root length in this species (Figure 2C).

**Figure 2.**
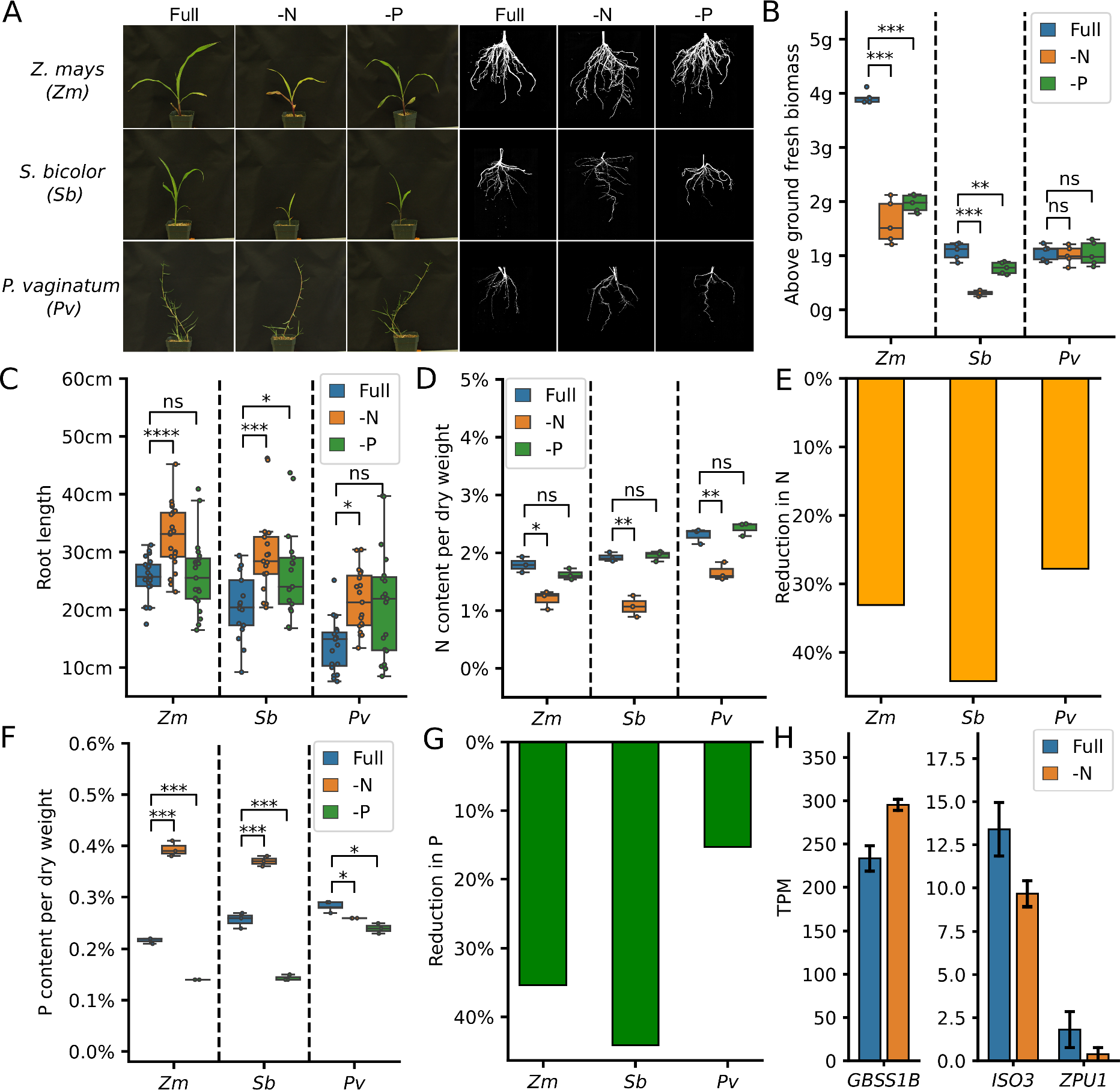
Responses of maize (*Zea mays*), sorghum (*Sorghum bicolor*), and paspalum (*Paspalum vaginatum*) to nutrient-deficit stress. (A) Representative images of above and below ground organs of maize, sorghum seedlings, and paspalum ramets at 21 days after panting (dap) under optimal (Full), nitrogen-deficit (-N), or phosphorus-deficit conditions (-P). (B) Change in fresh biomass accumulation under -N or -P conditions in maize, sorghum, and paspalum at 21 dap. (C) Changes in root length relative to Full at 21 daf under -N or -P conditions in maize, sorghum, and paspalum. (B-C) (* = p <**0.05**; *** = p <**0.0005**;**** = p <**0.00005**; t-test). (D-E) Abundance (D) and reduction (E) of N as a proportion of total dry biomass in the shoots of maize, sorghum seedlings, and paspalum ramets at 21 dap. (F-G) Abundance (F) and reduction (G)of P as a proportion of total dry biomass in the shoots of maize, sorghum seedlings, and paspalum ramets at 21 dap. (H) Change in the observed mRNA expression of the conserved and expressed paspalum orthologs of maize genes known to encode starch synthase (*GSSS1B*) and starch debranching enzymes (*ISO3* and *ZPU1*) in shoots under N-deficient or control conditions.

One potential explanation for the relative lack of plasticity observed in paspalum in response to N-deficit or P-deficit stress is that paspalum has limited potential to utilize N under optimal conditions and hence did not experience substantial internal declines in N availability in response to N-deficient conditions. We therefore measured the contents of N and P in plants of all three species included in this study. Substantial decreases in N as a proportion of total biomass were observed in all three species grown under N-deficient conditions relative to the full nutrient controls (Figure 2 D & E). P-deficient treatments produced significant declines in P as a proportion of dry biomass for all three species (Figure 2 F), although the decline in P abundance for paspalum was notably smaller in magnitude than the declines in maize and sorghum, with sorghum exhibiting the greatest reduction in P content (43%), followed by maize (36%) and paspalum (15%; Figure 2 G). N-deficient treatment produced significant increases in P as a proportion of dry biomass in maize and sorghum (Figure 2 F), which is consistent with previous reports of enhanced P uptake in plants grown under N-deficient conditions^37^. N deficit stress is also associated with increased starch accumulation in maize^38, 39^. In shoot tissues of paspalum seedlings grown under N-deficit conditions, the expression of the syntenically conserved gene *GBSS1B* (encoding granule-bound starch synthase 1)^40^ increased and the expression of *ISO3* and *ZPU1*(encoding starch debranching enzyme involved in starch degradation) decreased^41^ relative to nutrient optimal conditions (Figure 2 H). Taken as a whole, these results indicate that the external N-deficient treatment protocol employed here was sufficient to produce declines in internal N levels and N-deficit stress in paspalum.

### Comparisons of primary metabolic responses to nutrient-deficit stress

Numerous metabolic changes were observed between plants grown under nutrient-replete and nutrient-deficient conditions, with more metabolites exhibiting significant changes in abundance in response to N- or P-deficit stress in maize and sorghum than in paspalum (Figure 3A & B, Supplementary note 5). Twelve metabolites showed significant decreases in abundance in response to N-deficit stress in both sorghum and maize, and four metabolites showed significantly increased abundance in response to N-deficit stress (18 of the 32 metabolic responses were shared between the two species) (Figure 3A). A smaller number of statistically significant metabolic changes were observed in response to P-deficit stress, which is consistent with the less severe phenotype observed for P deficiency in the experimental design employed (Figure 3 B; Figure 2 A). A number of metabolic changes were again shared between maize and sorghum, with the levels of five tested metabolites decreasing in both species in response to P-deficit stress and one increasing (6 of the 16 metabolic responses were shared) (Figure 3B). All metabolic changes associated with N-deficit stress in paspalum were either species specific or shared with both maize and sorghum while all metabolic changes associated with P-deficit stress in paspalum were species specific (Figure 3 A & B). Metabolic changes associated with N-deficit stress shared by maize and sorghum but not paspalum included decreases in the abundance of many amino acids, including L-asparagin, L-glutamine, L-alanine and L-threonine (Figure 3A). This observation is consistent with the decreases in amino acid metabolism observed under N-limited conditions^42, 43^. A conserved increase in the abundance of caffeic acid was detected in both maize and sorghum in response to N-deficit conditions, which is consistent with previous reports from rice grown under similar N-limited conditions^44^ (Figure 3A).

**Figure 3.**
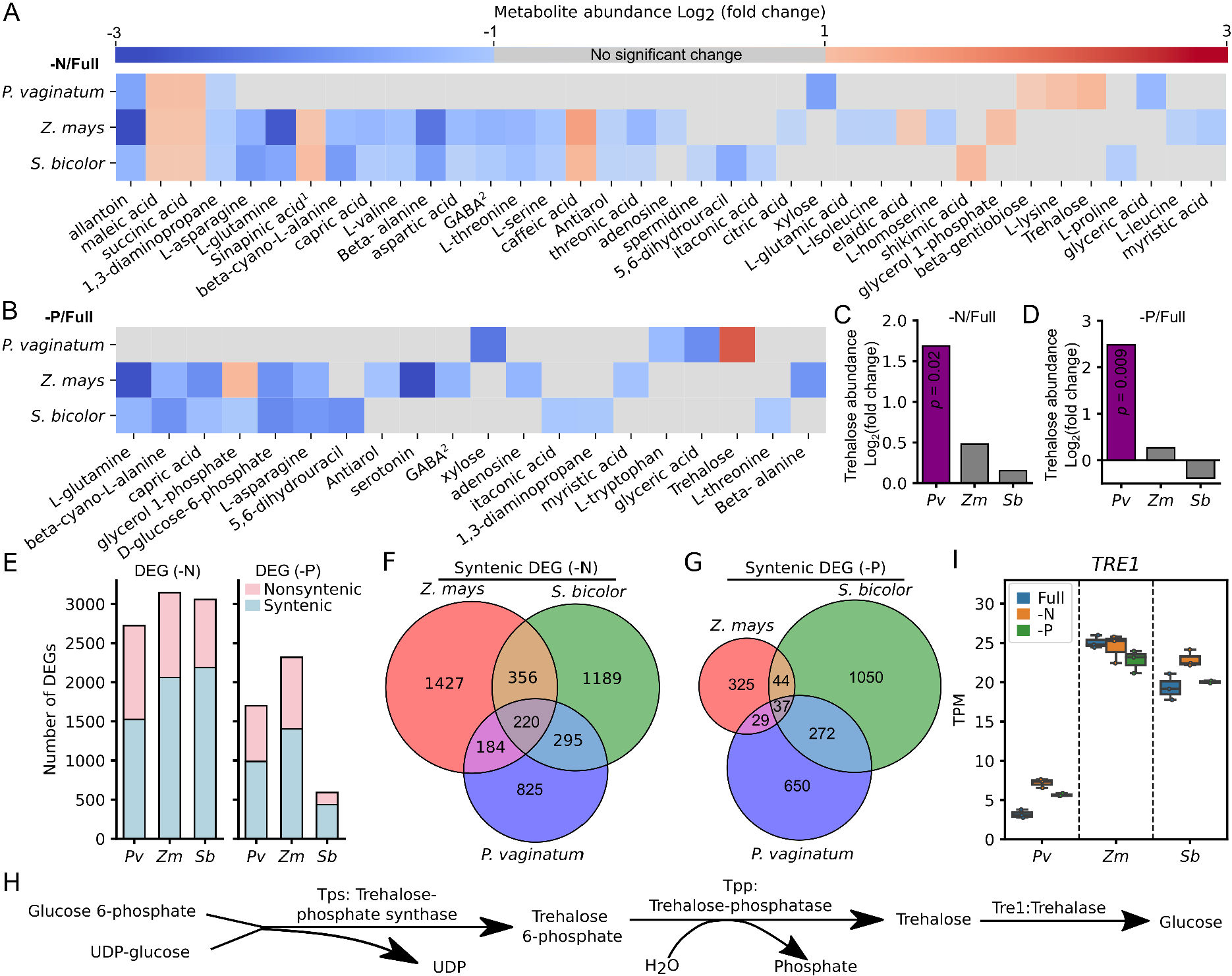
Primary metabolic and transcriptomic responses of maize (*Zea mays*), sorghum (*Sorghum bicolor*), and paspalum (*Paspalum vaginatum*) to nutrient-deficit stress. (A-B) Changes in the abundance of metabolites in the roots of maize, sorghum seedlings, and paspalum ramets grown under -N conditions (A) and -P condition (B) at 21 dap relative to plants grown under Full condition. Only the metabolites with a statistically significance change in abundance (p <0.05; t-test) and an absolute fold change >2 in at least one of the three species evaluated are shown. Cell marked in gray were not significantly different between conditions and/or exhibited an absolute fold change less than 2. ^1^3,5-dimethoxy-4-hydroxycinnamic acid; ^2^Gamma-aminobutyric acid. Raw data for fold change and t-test results are shown in Supplemental Document 2. (C-D) Change in trehalose abundance in the roots of 3-week-old maize, sorghum seedlings, and paspalum ramets under -N condition (C) and -P condition (D) relative to plants grown under Full condition. Statistically significant changes are indicated in purple (t-test), and non-statistically significant changes are indicated in gray. (E) Number of significantly differentially expressed genes in paspalum (*Pv*), maize (*Zm*) and sorghum (*Sb*) identified in comparisons between roots of 3-week old plants grown under either -N or -P conditions and Full condition. Shading indicates the proportion of differentially expressed genes (DEGs) in each species that are syntenically conserved across species, or present at a unique location in the genome of the individual species evaluated. (F-G) Number of syntenically conserved orthologous triplets exhibiting shared or species-specific differential expression in response to -N and -P conditions (G) in maize (*Z. mays*), sorghum (*S. bicolor*), and paspalum (*P. vaginatum*). (H) Simplified diagram for trehalose metabolic pathway. (I) Expression levels of trehalase-encoding genes in paspalum (*Pv*), maize (*Zm*) and sorghum (*Sb*) in roots of 3-week-old plants grown in nutrient optimal (Full), N-deficient (-N) and P-deficient (-P) conditions.

All three species exhibited decreases in 1,3-diaminopropane and allantoin levels under N-deficit conditions (Figure 3A). Allantoin acts as a pool of relocalizable N that can be catabolized into ammonia for N assimilation and amino acid biogenesis^45^. In addition, the abundance of both succinic acid and maleic acid (MaA) increased in all three species in response to N-deficit stress (Figure 3B). Maleic acid produced and secreted in response to another abiotic stress (drought) in holm oak (*Quercus ilex*)^46^. As we examined internal metabolite abundance but did not profile root exudates in the current study, it is not possible to determine whether the internal accumulation of maleic acid resulted in additional secretion in these three grass species. Succinate, the anion of succinic acid, forms part of the tricarboxylic acid (TCA) cycle. The increase in succinic acid levels, combined with the decreased abundance of gamma-aminobutyric acid (GABA), is consistent with these species employing the GABA shunt pathway, which was proposed to act as an additional energy source to support cellular metabolism under stress conditions^47–49^.

We observed changes in metabolite abundance in maize and sorghum grown under P-deficient conditions relative to the nutrient optimal conditions, including L-asparagine, GABA, L-glutamine, L-alanine, capric acid, D-glucose-6-phosphate and glycerol-1-phosphate. However, none of these metabolites exhibited significant changes in abundance in paspalum plants grown under P-deficient and vs. nutrient optimal conditions (Figure 3 B). The abundance of D-glucose-6-phosphate (D-G6P), the primary entry molecule for glycolysis was significantly lower in maize and sorghum plants grown under P-deficient conditions vs. those grown under nutrient optimal conditions (Figure 3A). The reduction in D-G6P levels might reflect the lack of free phosphate available to produce adenosine triphosphate (ATP) to drive the phosphorylation of glucose, as P is a major component of ATP. The abundance of D-G6P did not decrease in paspalum plants grown under P-deficient conditions (Figure 3 B). None of the metabolites that exhibited significant changes in abundance in paspalum between nutrient optimal and P-deficient conditions, including tryptophan, xylose, glyceric acid, and trehalose exhibited changes in abundance in maize or sorghum (Figure 3 A-D). The abundance of trehalose, a di-saccharide that predominantly functions as a signaling molecule in plants in response to abiotic stresses, was significantly higher in paspalum plants grown under N-deficient or P-deficient conditions vs. the nutrient optimal conditions, but this difference was not observed in maize or sorghum (Figure 3 C & D).

### Conserved and differential transcriptomic responses of paspalum to nutrient-deficit conditions

The sequencing, assembly, and annotation of the paspalum genome provided the opportunity to quantify differences and commonalities in how maize, sorghum, and paspalum transcriptionally respond to nutrient-deficit stress. We collected RNA from the root tissues of three biological replicates of each species and used it to generate an average of approximately 40 million high-quality reads per sample. Principal component analysis based on the transcriptomes of each sample showed a clear separation based on growth conditions in maize (Figure S4 A), sorghum (Figure S4 B) and paspalum (Figure S4 C). We identified 3,057, 3,144 and 2,723 genes with significantly differential expression levels between nutrient optimal and N-deficit stress conditions in maize, paspalum, and sorghum, respectively. In addition, 591, 2,318 and 1,698 genes showed significantly differential expression levels between nutrient optimal and P-deficit stress conditions in maize, paspalum, and sorghum, respectively (Figure 3 E).

Most differentially expressed genes (DEGs) identified for each treatment in each species were themselves syntenically conserved (Figure 3 E). Members of a number of paspalum specific expanded gene families showed significant transcriptional responses to N-deficit stress (Figure S5A) and/or P-deficit stress (Figure S5B). However, consistent with a previous study of transcriptional responses to abiotic stress^50^, the conservation of transcriptional responses was much less common than the conservation of the genes themselves; syntenic genes that showed significant fold changes varied across the three species under N-deficient and P-deficient conditions (Figure 3 F & G).

The set of 220 syntenically conserved orthologous gene groups that responded transcriptionally to N-deficit stress in a consistent fashion among maize, sorghum, and paspalum was disproportionately enriched in GO terms related to response to nutrient levels, nitrate assimilation, metal ion transporter activities and divalent inorganic cation transmembrane transporter activity (Figure 3F; Figure S6A). The set of 37 syntenically conserved orthologous gene groups that responded transcriptionally to P-deficit stress in a consistent fashion among the three grasses was disproportionately enriched in GO terms related to lipid metabolic process, phosphate ion transport, response to nutrient levels and cell communication (Figure 3G; Figure S6A). Syntenically conserved orthologous gene groups where a transcriptional response to N-deficit stress was unique to paspalum where enriched in genes involved in proton transport, glycoside biosynthetic process and serine family amino acid metabolic process (Figure 3 F; Figure S6 B)). By contrast, the syntenically conserved orthologous gene groups that were uniquely differentially expressed in paspalum in response to P-deficit stress were involved in processes related to antioxidation, gene regulation and primary metabolism (Figure 3 G; Figure S6 B).

The significant accumulation of trehalose in paspalum in response to nitrogen deficient conditions motivated us to examine the expression of genes involved in the trehalose metabolic pathway including the genes encoding enzymes that catalyze three steps in trehalose metabolism: trehalose-6-phosphate synthase, trehalose-6-phosphate phosphatase, and trehalase (Figure 3 H). Two maize genes encoding trehalose-6-phosphate synthase 1 and 12 are syntenic homeologs resulting from the maize whole-genome duplication and are co-orthologous to single gene copies in sorghum and paspalum. These genes formed a clade sister to the well characterized *Arabidopsis thaliana* gene *AtTPS1*^51^ which is consistent with the previous study that characterized *ZmTPS1*^52^ (Figure S7 A; Supplementary Note 6). Copies of this gene in both sorghum and paspalum showed a significant increase in mRNA abundance in response to N-deficient treatment, as did the maize gene encoding trehalose-6-phosphate synthase 1 (*ZmTPS1*), which possesses all catalytic domains of TPS (Figure S7 B). Plots of the detectable transcriptional responses of other trehalose-6-phosphate synthase encoding homologs are shown in Figure S7 C - I. Genes annotated as encoding trehalose-6-phosphate phosphatase 6 (*ZmTPP6*) and trehalose-6-phosphate phosphatase 11 (*ZmTPP11*) were phylogenetically clustered with Arabidopsis *AtTPPA* (Figure S8 A-C; Supplementary Note 7), and both tended to be differentially expressed between control and stress conditions in all three species. Similar transcriptional responses of homologs encoding other trehalose-6-phosphate phosphatases were observed across all three species (Figure S8 D-I). Trehalase, an enzyme that breaks trehalose down into two molecules of glucose, is encoded by a single gene copy in maize, with conserved syntenic orthologs in sorghum and paspalum. Lower levels of mRNA abundance were associated with the trehalase encoding gene in paspalum than its syntenic orthologs in sorghum or maize (Figure 3 I).

### Inhibiting trehalase activity in maize and sorghum recapitulates the paspalum phenotype

As shown above, paspalum exhibited a significant accumulation of trehalose in response to the two types of nutrient-deficit stresses while maize and sorghum did not (Figure 3 C & D). However, as equivalent P-deficient treatments introduced notably smaller changes in P abundance in paspalum relative to maize and sorghum, we elected to focus exclusively on N-deficit stress in all subsequent experiments.

In an attempt to phenocopy the reduced plasticity in response to N-deficient treatment originally observed in paspalum, we treated maize and sorghum plants with validamycin A (β-d-glucopyranosilvalidoxylamine, ValA) – a specific inhibitor of trehalase activity^53–55^. Visibly better growth under both optimal and nitrogen deficient conditions was observed in maize (Figure4A) and a slightly better growth under nitrogen deficient condition was observe in sorghum but no obvious changes in growth under both conditions was observed in paspalum (Figure S9A & E). Metabolic profiling of treated and untreated plants confirmed that a treatment with 30 µM ValA significantly increased the accumulation of trehalose under both optimal and N-deficient nutrient conditions in maize and sorghum (Figure 4B; Figure S9B) but failed to increase trehalose accumulation in paspalum (Figure S9F). ValA treatment produced significant increases in dry biomass accumulation in maize under both control and N-deficient treatment (Figure 4 C) and in sorghum only under N-deficient treatment (Figure S9 C). ValA treatment did not significantly alter biomass accumulation in paspalum under either treatment condition (Figure S9 G). Nutrient-deficit stress is known to alter shoot-to-root biomass ratios, increasing root biomass as a percentage of the total biomass^56, 57^. Root biomass made up a smaller proportion of the total biomass for both maize and sorghum seedlings treated with ValA than untreated seedlings under both full-nutrient and N-deficient conditions (Figure 4 D; Figure S9 D). However, no significant changes in shoot-to-root ratio were observed in paspalum upon ValA treatment irrespective of nutrient conditions(Figure S9 H).

**Figure 4.**
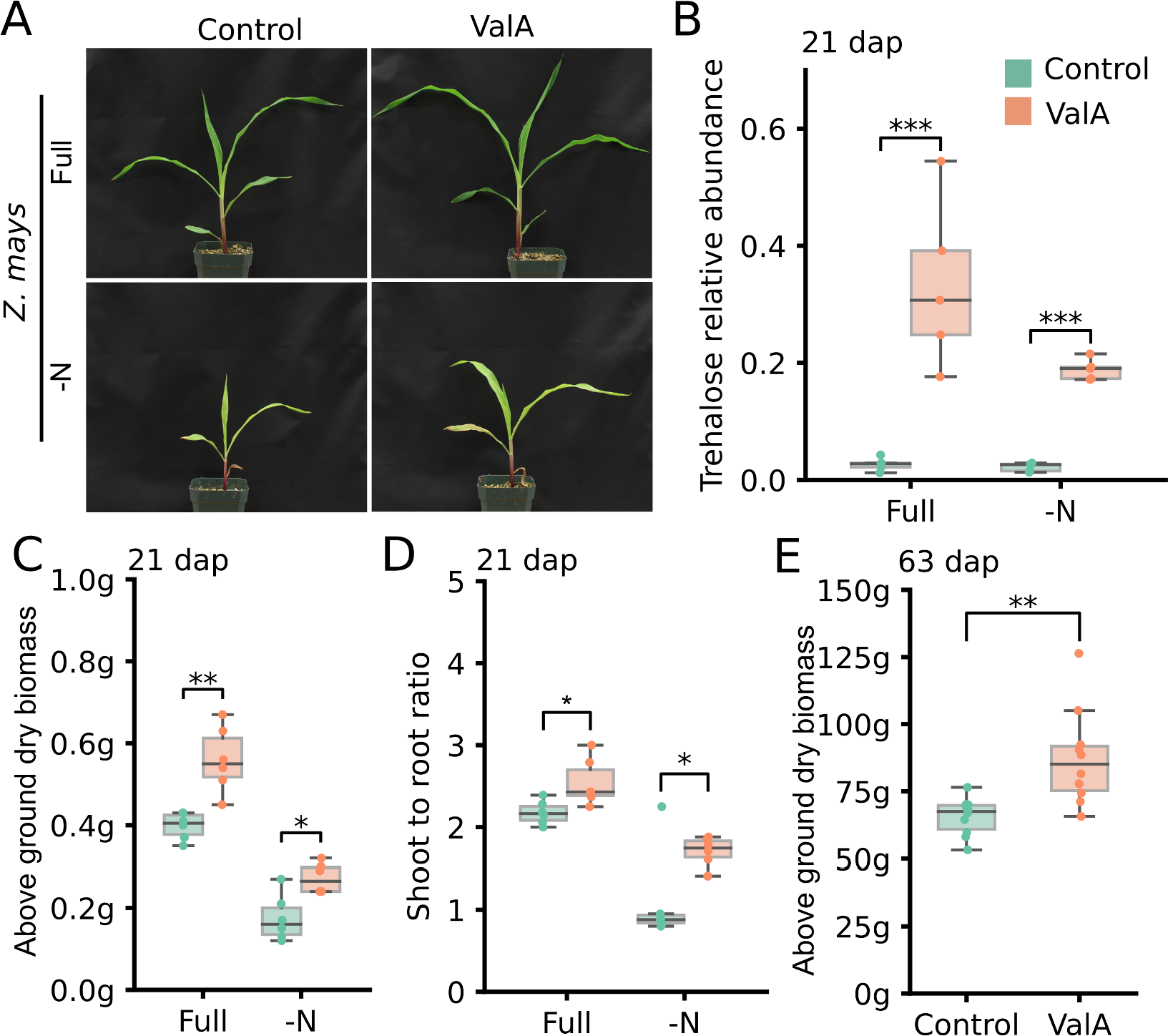
Validamycin A treatment is associated with increased trehalose accumulation and greater biomass production in maize. (A) Representative images showing maize seedlings at 21 dap grown under Full and -N conditions with (ValA) or without (Control) validamycin A treatment. (B) Changes in observed trehalose abundance – normalized relative to an internal reference (ribitol) – in response to validamycin A (ValA) under Full and -N in maize root tissue from seedlings at 21 dap. (C) Changes in the above ground dry weight of maize seedlings at 21 dap in response to validamycin A (ValA) treatment under Full or -N conditions. (D) Ratio of shoot-to-root dry biomass in 3-week-old maize seedlings grown under Full and -N conditions with (ValA) or without (Control) validamycin A treatment. (E) Statistically significant increases in biomass accumulation observed in late vegetative stage (63 dap) validamycin A (ValA) treated maize relative to control (untreated) plants under Full condition. (** = p <**0.005**; *** = p <**0.0005**; t-test)

To extend our observations beyond the late seedling stage, we grew a cohort of maize plants for for 63 days (until the late vegetative stage) under either control or ValA treated conditions. ValA treated plants accumulated significantly more biomass than control plants grown as part of the same experiment (control mean = 65.6 grams/plant, ValA mean = 87.3 grams/plant; p = 0.002; t-test) (Figure 4 E). In a preliminary experiment, a smaller number of maize plants were grown under either control or ValA treated conditions to reproductive stage (Figure S10 A & B). ValA treated plants flowered earlier (Figure S10 C) and produced larger tassels (Figure S10 B & D) and leaves than their untreated siblings (Figure S10 E). In previous studies, genetically modifying trehalose metabolic pathway altered photosynthesis and nutrient partitioning in maize reproductive tissues, thereby affecting yields, via its effect on SnRK1 activity^58, 59^. However, in the current smaller experiment, ValA induced differences in above ground biomass accumulation, including both tassels and ear shoots as well as vegetative tissues at the reproductive stage, were not statistically significant (Figure S10 F). Perhaps this was due to an earlier transition to reproductive development in ValA treated plants, or perhaps because of low statistical power, we failed to detect differences between such small numbers of plants. A set of 27 genes associated with trehalose metabolism exhibited significantly more rapid rates of protein sequence evolution in paspalum than did the orthologs of these same genes in foxtail millet (*S. italica*, p = 0.002), sorghum (*S. bicolor*, p = 0.014) and *Oropetium* (*O. thomaeum*, p = 0.025) (Figure 4 E). These data are consistent with, but not conclusive evidence for, a role for trehalose metabolism in the reduced phenotypic plasticity paspalum exhibits in response to a range of abiotic stresses such as salinity^60^.

### ValA treatment is associated with increased autophagy in maize

A number of potential mechanisms could explain the association between the ValA associated increases in trehalose accumulation and increased growth under nutrient deficient conditions. Relative to other disaccharides, trehalose accumulates to only low levels and is thought to act as a signal rather than a carbon source^66, 67^. The precursor to trehalose, trehalose-6-phosphate, has been shown to regulate cell growth by inhibiting SNRK1 activity^68–71^. Hence, one potential model to explain the observed result is that treatment with ValA, which inhibits trehalase activity and increases trehalose accumulation^53^, might also increase the abundance of trehalose-6-phosphate, one step earlier in the pathway^69, 72^. Consistent with this model, the maize ortholog of the Arabidopsis gene encoding trehalose-6-phosphate phosphatase A was significantly downregulated in ValA treated plants relative to control samples under both full-nutrient and N-deficient treatment conditions (Figure 5 A; Figure S8 A; Supplementary Note 6). The trehalose-6-phosphate synthase encoding maize gene *ZmTPS1*^52^ also exhibited significant declines in expression in response to ValA treatment under both full-nutrient and N-deficient treatment conditions (Figure 5 B; Supplementary Note 7). Expression change of the genes encoding other redundant TPSs and TPPs did not show specific patterns (Figure S11 A). Furthermore, the maize gene encoding the SNRK1 alpha subunit A (Zm00001d038745)^64^ was upregulated in response to ValA treatment under both Full and -N conditions (Figure 5 C). In addition, the known SNRK1 induced gene in maize *ZmAKIN11* (Zm00001d028733) (Figure S11 B) was significantly up-regulated, the known SNRK1 repressed genes in maize *ZmMDH3* (Zm00001d044042) (Figure S11 C), *ZmMDH6* (Zm00001d031899) (Figure S11 D) and (ZmBZIP11) (Figure S11 E)^73, 74^ were significantly down-regulated. While abundance of trehalose-6-phosphate was not directly assayed, these transcriptional changes observed in current study were consistent with an increased SNRK1 activity in ValA treated plants and, hence, inconsistent with increases in the abundance of trehalose-6-phosphate which inhibits SNRK1 activity^68, 69^. The expression of a number of ammonium and nitrate transporters in root tissues from maize seedlings treated with ValA were upregulated relative to untreated seedlings whether grown under Full and -N treatment conditions (Figure 5D).

**Figure 5.**
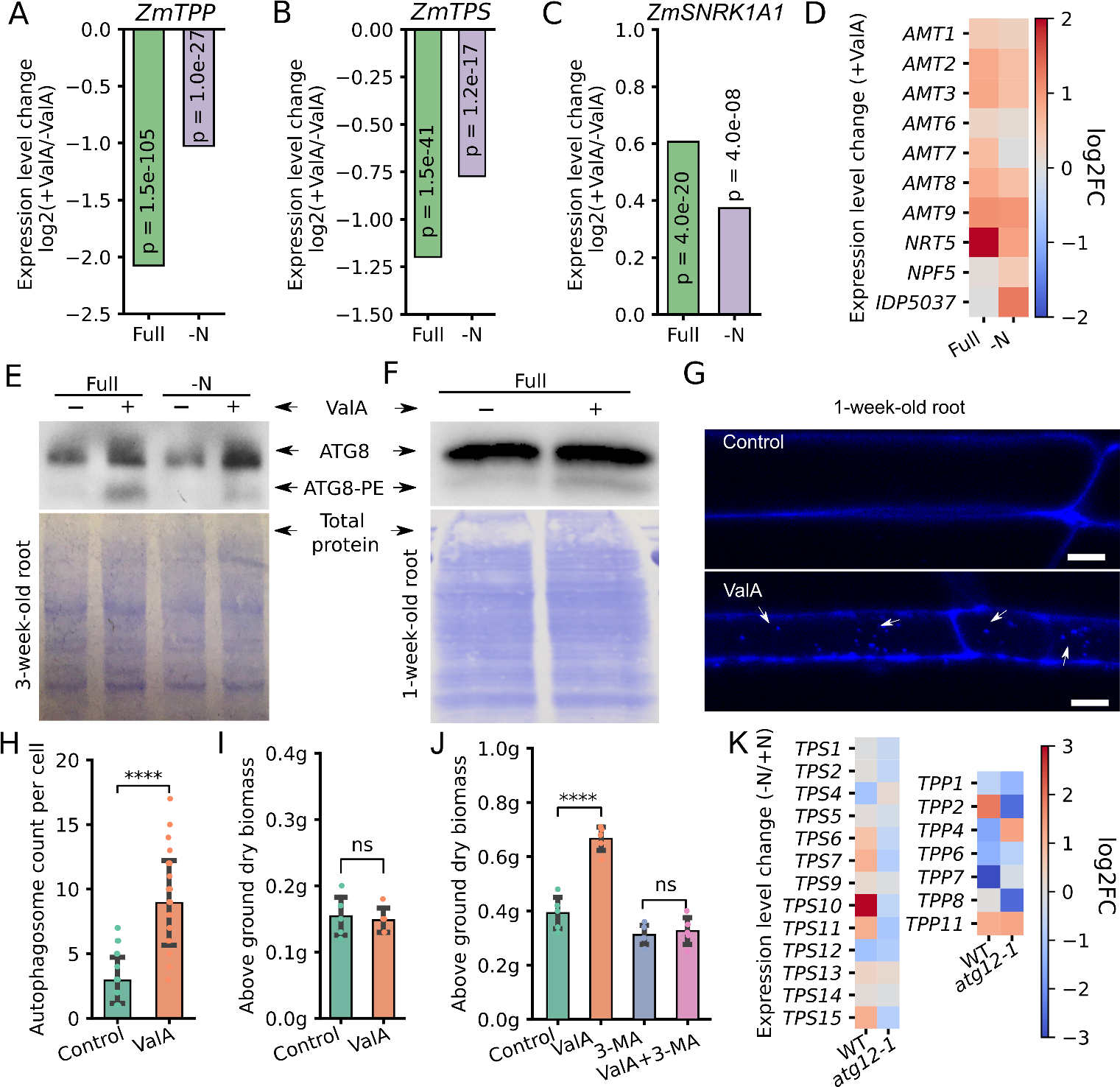
Evidence for increased autophagy in maize seedlings treated with validamycin A. (A-B) Decrease in the expression of the trehalose-6-phosphate synthase encoding gene *ZmTPS1* (A)^52^ and the trehalose-6-phosphate phosphatase enconding maize gene *ZmTRPP6*^58, 61, 62^ (B) in root tissues from ValA treated maize seedlings relative to control seedlings under both Full and -N conditions at 21 dap. (C) Increase in the expression of the SNRK1 alpha subunit enconding maize gene *SNRK1A1*^63, 64^ in roots from three week old ValA treated maize seedlings relative to control seedlings under both full nutrient and N-deficient treatment conditions. (D) Uniform upregulation of genes encoding ammonium and nitrate transporters. (E-F) Immunoblot measuring the abundance of both free ATG8 (upper band) and the ATG8-PE conjugate (lower band) in root samples collected from 3-week-old maize seedlings grown under Full and -N conditions with or without ValA treatment and 1-week-old maize seedlings grown under Full conditions with or without ValA treatment. Total protein loading control is shown in the lower panel. (G-H) Microscopy images (G) and counts (H) of autophagosomes stained by 40 µM MDC (monodansylcadaverine) in root tips of 1-week-old maize seedlings grown under Full condition with or without ValA treatment (**** = p <**0.00005**, t-test). (I) Above around dry biomass accumulated in 1-week-old maize seedling grown under Full condition with or without ValA treatment (ns = p >**0.05**, t-test) (J) Accumulation of above ground dry biomass for 3-week-old control seedlings, seedlings treated with 3 mM 3-MA (3-methyladenine), 30 µM ValA or both 3 mM 3-MA and 30 µM ValA. (* = p <**0.05**; **** = p <**0.00005**; ns = No significance; t-test) (K) Detectable changes in the expression levels of annotated maize genes encoding trehalose-6-phosphate synthase (TPS) or trehalose-6-phosphate phosphatase (TPP) in response to nitrogen deficient treatment in both the wild type and *atg12-1* mutant backgrounds^65^

SNRK1 is an upstream promoter of autophagy^61, 63, 75–77^ and more rapid turnover of damaged or unneeded cellular components and proteins allows for more growth with a fixed quantity of N supply (Figure (S11F)). During autophagy, the protein ATG8 becomes conjugated to phosphatidylethanolamine (PE). Increases in the abundance of both free ATG8 and ATG8-PE are associated with autophagy activation^78–80^. In maize, N-deficit stress did not produce any obvious change in the accumulation of either free ATG8 or ATG8-PE; however, under both control and N-deficit stress conditions, two independently replicated plants treated with ValA accumulated more free ATG8 and more ATG8-PE (Figure 5E & Figure S11G). Similar results were observed in two independently replicated one week-old-seedlings (Figure 5F & Figure S11H). Consistent to an increased autophagy activity, the abundance of autophagosomes stained by Monodansylcadaverine (MDC) in root tip cells from ValA treated plants was approximately twice as high as the untreated plants (Figure 5G & H). At this stage of growth, no differences in growth could be observed (Figure 5I) as the seedlings were still utilizing the nutrients stored in the cotyledon and therefore the increase in autophagy under nutrient optimal conditions did not result from possible nutrient deficiency stress caused by higher nutrient consumption in faster growing ValA-treated plants, instead, was due to ValA treatment.

We treated seedlings grown under optimal nutrient conditions with 3-methyladenine (3-MA, a phosphatidylinositol 3-kinase (PI3K) inhibitor) that prevents autophagosome formation^80, 81^, ValA, or both. Treatment with 3-MA alone slightly reduced seedling growth. As previously observed, treatment with ValA alone produced significant increases in biomass accumulation. However, in the presence of 3-MA, no significant change in biomass accumulation was observed in response to ValA treatment (Figure 5 J). In an *atg12-1* mutant background, maize genes encoding TPS were predominately down-regulated in response to N-deficient treatment, while in wild-type controls, several of the same genes exhibited increased expression in response to N-deficient treatment (Figure 5 K)^65^. In data taken from the same experiment, maize genes encoding TPP showed varying responses to both N-deficit stress and genetic background (Figure 5 K)^65^.

## Discussion

In paspalum, a crop wild relative that is resilient to numerous abiotic stresses, nutrient-deficit stress was associated with substantial accumulation of trehalose. The sequencing of a reference genome for this species allowed us to perform comparative evolutionary analyses, which identified accelerated protein sequence evolution of genes involved in trehalose metabolism in paspalum (Figure 4 J). Treating maize and sorghum exposed to N-deficit stress with a specific inhibitor of trehalase resulted in higher internal trehalose accumulation and recapitulated a number of paspalum phenotypes including reduced decreases in biomass accumulation in response to N-deficit stress, and increased allocation of biomass to shoots under N-deficit stress (Figure 4; Figure S9).

Imposing equivalent stress treatment protocols across species presents numerous challenges. One potential concern with the initial finding that paspalum is less phenotypically plastic in response to nutrient-deficient treatment than maize is that the slower baseline accumulation of biomass in paspalum may deplete the modest reserves of nitrate and phosphate in soil more slowly than they would be used by maize. Here comparison of paspalum to sorghum may be more informative than comparison of paspalum to maize. Under nutrient replete conditions, individual paspalum ramets accumulated approximately equivalent amounts of biomass to sorghum seedlings, while sorghum exhibited greater phenotypic plasticity in response to nutrient deficit stress. In addition, several lines of evidence indicate that the three species experienced nutrient deficit stress in response to the nutrient deficient treatment protocols employed in this study: the depletion of the nitrate storage compound allantoin under the N-deficit conditions as well as other metabolic changes (Figure 3 A & B); transcriptional evidence of increased starch biosynthesis in paspalum shoots (Figure 2 H); and significant declines in the abundance of N and P in the above-ground tissue all three species when grown under N- and P-deficient conditions (Figure 2 E & G).

In many flowering plant species, the abundance of trehalose is quite low^82^. In *Arabidopsis thaliana*, trehalose accumulation was observed under high and low temperature stress^83, 84^, high light intensity^85^, high cadmium levels^86^ and dehydration^87^, but no increase was observed under N-deficient conditions^82, 88^. A non-targeted metabolic profiling of six legume plants in the *Lotus* genus under drought conditions revealed a significant increase in trehalose abundance across all species tested^89^. In maize, external trehalose treatment enhanced antioxidant activities under high salinity and P-deficient conditions, thus achieving better seedling growth^90^. Genetically modified rice plants that over-expressed a fusion gene encoding both *Escherichia coli* Trehalose-6-phosphate synthase and Trehalose-6-phosphate phosphatase, which are responsible for trehalose biosynthesis, exhibited 200-fold greater accumulation of trehalose and significantly higher tolerance to drought, salinity, and cold stress^91^. Associations between trehalose and nutrient-deficit stress appear to have been largely uninvestigated, although one study found that the exogenous application of trehalose to *Nicotianan benthamiana* leaves partially rescued the N-deficiency phenotypes under N-limited conditions^92^. The transgenic expression of trehalose-6-phosphate phosphatase in developing maize ears was associated with increased yield under control and drought stressed conditions^59, 71^. By contrast, the increased accumulation of trehalose in response to nutrient-deficit stresses in wild-type plants is, to our knowledge, specific to paspalum. Given the environment that paspalum has adapted to during evolution, paspalum has an extraordinary ability to tolerate high-salinity stress^19, 29, 93^. Trehalose accumulation in paspalum might also act to ameliorate osmotic stress caused by a larger amount of salt uptake from the soil driven by a higher transpiration when nutrient deficient plants seek to increase nutrient uptake.

Trehalose has been recognized as an autophagy activator in animals^94–96^ and plants^76^. Autophagy plays pivotal roles in proteome remodeling, lipid turnover^65, 76^, nitrogen remobilization^97–100^, nitrogen use efficiency^101^, and abiotic stress responses in a variety of plant species (as reviewed previously^102^). In the resurrection plant *Tripogon loliiformis*, trehalose abundance correlated with an increase in ATG8 lipidation and the number of autophagosomes^79^. Here, we pharmaceutically inhibited trehalase activity with ValA to increase treahalose abundance in maize and observed increases in both ATG8 protein abundance and lipidation in both 3-week-old and 1-week-old maize seedlings (Figure 5 E & F). The effect of ValA treatment on biomass accumulation in maize was autophagy dependent as inhibiting autophagy via treatment with 3-MA restored the wild-type phenotype of ValA treated plants (Figure 5 H). In addition, we observed upregulation of both ammonium and nitrate transporter expression in the seedlings treated with validamycin A under both Full and -N conditions (Figure 5 D) suggesting that validamycin A treated seedlings may performed better not only as a result of nitrogen recycling and remobilization, but could also exhibit increased nitrogen uptake due to the upregulated transporter activities in roots. However, the reversion of validamycin A treated seedlings to wild type levels of biomass accumulation with treated with an autophagy inhibitor suggests that the role of increased nitrogen uptake, if any, is likely also autophagy dependent. The triggering of trehalose accumulation in response to nutrient deficit is specific to paspalum (Figure 3 A-D). However, caution should be taken in interpreting these results as multiple genes encoding enzymes in the trehalose biosynthetic pathway were also reported to be associated with changes in autophagy activity in different systems^68, 103^. Trehalase activity was initially observed in tissue cultures generated from a range of plant species more than three decades ago^104^. Inhibiting trehalase activity with ValA can control wheat Fusarium head blight (FHB) and inhibit Deoxynivalenol (DON) contamination^105^. Over-expression of *OsTRE1* in rice was associated with improved salt tolerance^106^ and over-expression of *AtTRE1* in Arabidopsis was associated with improved drought tolerance^107^. However, these known phenotypic consequences of alterations in trehalase activities would not necessarily predict an association between inhibition of trehalase activity and decreased plasticity in response to nutrient deficiency stress, as was observed here.

The maize experiments described in this paper would not have been conducted in the absence of the observation that paspalum accumulates trehalose in response to nutrient deficient treatments. However, at the same time it must be noted that the work linking trehalose accumulation to increased biomass accumulation via an autophagy dependent mechanism was conducted entirely in maize. Hence, while the maize data certainly suggests that an increase in autophagy induced by the increased accumulation of trehalose in paspalum observed under nutrient deficient conditions is also responsible for the low degree of phenotypic plasticity paspalum exhibits in response to nutrient deficit stress, strong tests of this model would require experimentation which is not yet practical, such as the transgenic overexpression of trehalase in paspalum. In any case, these results suggest that the manipulation of trehalose accumulation in maize and sorghum, as well as potentially on other domesticated grasses, whether chemically or via the modification of the expression of the endogenous trehalase enzyme, may increase agricultural productivity per unit of nitrate and phosphate fertilizer applied. Finally, the observation of autophagy dependent increases in biomass accumulation in even maize plants grown under nutrient-replete conditions suggests that current maize lines may exhibit a suboptimal level of autophagy in roots. However, again, caution should be taken in interpreting these results as, while increases in biomass accumulation were observed not only in seedlings but in late stage vegetative plants (Figure 4E), all data presented in this study were generated in controlled environmental conditions and changes in regulation or metabolism that are beneficial in the greenhouse may or may not generalize to the field.

## Materials and methods

### Determination of DNA content via flow cytometry and genome size estimation

One leaf per plant of paspalum (PI 509022) and sorghum (BTx623) were harvested and kept on ice until processing. A CyStain Propidium Iodide Absolute P kit (Sysmex, Milton Keynes, United Kingdom) was used to extract and stain the nuclei from a 1 cm^2^ piece of leaf tissue following the manufacturer’s instructions. To reduce the amount of cellular debris in the extracts, samples were passed through a 30 µm filter (CellTrics®-Sysmex Partec, Goerlitz, Germany) and centrifuged at 600×g before final staining. Sorghum was used as an internal standard to reduce the staining variability between samples. The stained samples were then analyzed on a CytoFLEX flow cytometer (Beckman Coulter, Brea, CA, USA) following a two-hour incubation at 4 °*C*. The propidium iodide was excited with a yellow-green 561 nm laser and detected with a 585/42 emission filter. The genome size was calculated for a total of six samples. The 2C genome size for BTx623 is 1.67 pg DNA^108^; therefore the formula to calculate the DNA content of paspalum was (median fluorescence_sample nuclei_ /median fluorescence_standard nuclei_) X 1.67 pg. The mean and standard error of these six samples were calculated, and the mean was converted to a 1C genome size using the conversion factor 1 pg = 980 Mbp.

### Paspalum genome assembly and annotation

The paspalum genome assembly was generated using an error-corrected 74.3x coverage of PacBio reads with an average read length of 9,523 bp. The reads were assembled using MECAT^109^ and polished using QUIVER^110^. Comparisons with the genome of *Panicum hallii* var. HAL2 (v2) and two paspalum F_2_ genetic maps (see Supplemental Methods for map generation) were used to identify and split 15 misjoins in the initial assembly. The resulting scaffolds were ordered and orientated using the two paspalum genetic maps. A total of 357 scaffolds were assembled into 10 pseudomolecules representing 75% of the overall assembled genome. A set of six F_1_ maps (total of 8,861 markers)^30^ were used to refine the order/orientation of the contigs. The final numbering and orientation were verified using *S. bicolor* cDNAs obtained from Phytozome (https://phytozome-next.jgi.doe.gov/). Heterozygous SNP/InDel phasing errors were corrected using both 74.3X raw PacBio data and 78X Illumina data (San Diego, CA, USA) (2x150 bp reads, 400 bp target insert size). A detailed genome assembly methods and assembly integrity assessment is provided in Supplementary Note 1.

The v3.0 paspalum genome assembly was annotated using a combination of an alignment of assembled transcripts from paspalum and protein sequences from other plant species. Prior to its annotation, the genome assembly was first repeat-masked using both known repeats from RepBase and *de novo* identified repetitive sequences from RepeatModeler^111, 112^. Transcript assemblies were generated via a two-stage assembly process utilizing PERTRAN followed by PASA^113^. A total of 112,258 RNAseq transcript assemblies were generated from approximately 1.6 billion 2x150 bp strand-specific Illumina sequencing reads. Protein sequences from Arabidopsis, soybean, sorghum, Kitaake rice (*Oryza sativa Kitaake*), green foxtail (*Setaria viridis*), grape (*Vitis vinifera*), and the Swiss-Prot proteomes were aligned to the repeat-masked genome using EXONERATE^114^. Independent sets of gene models were predicted using FGENESH+, FGENESH_EST, EXONERATE and AUGUST as implemented in BRAKER1, and the in-house PASA assembly open reading frames (ORFs; in-house homology constrained ORF finder) tool from JGI^114–116^. For each locus, the prediction with the best score based on the expressed sequence tag (EST) and protein support and a lack of overlap with repeats was selected. The best prediction for each locus was further improved using PASA to add untranslated regions, correct splice sites, and add alternative transcripts. Improved transcripts were assessed based on both the C-score (ratio of the BLASTP alignment score to the mutual best hit BLASTP alignment score) and protein coverage. Transcripts were retained if any one of three criteria were met: 1) Transcripts where the C-score and protein coverage score were each ≥ 0.5 and less than 20% of the transcript overlapped with sequence annotated as repetitive. 2) Transcripts supported by EST coverage and less than 20% of the transcript overlapped with sequence annotated as repetitive. 3) Transcripts with a Cscore ≥ 0.9 and a protein coverage score ≥ 0.7, regardless of the proportion of overlap with annotated repeat sequences. Sequences that satisfied one or more of the above three criteria and where more than 30% of predicted protein sequence was covered by Pfam domains annotated as belonging to transposable elements were also removed. Short single exon (predicted coding sequence <300 bp) genes without protein domain support and expression data, incomplete gene models and those with low homology support (sum of Cscore and coverage <1.5 for complete, <1.8 for incomplete) and without full transcriptome support (CDS and intron coverage supported by any transcript assemblies) were removed. Gene models that passed all the criteria described above were included in the gene model annotations for paspalum. The GO terms assignment was based on the InterProScan results^117^.

### Plant materials and growth conditions

The maize (*Zea mays* ssp. *mays*), sorghum (*Sorghum bicolor*), and seashore paspalum (*Paspalum vaginatum*) genotypes used to create the reference genomes for each species were: accessions B73, BTx623, and PI 509022, respectively^118, 119^. Maize and sorghum seeds were surface sterilized in 2% bleach for 40 minutes, rinsed, and imbibed overnight in deionized distilled water (ddiH_2_O). The seeds were sown in a mixture of 20% MitroMix200, 30% sterilized sand and 50% fine vermiculite(v/v) and grown under greenhouse conditions (temperature: 22-29°*C* with a 14-h light: 10-h dark photoperiod). The heterozygous reference clone PI 509022 was obtained from the USDA National Plant Germplasm Service and propagated via rhizome cuttings using the same growth medium and conditions used for sorghum and maize. All plants were watered with sterilized ddiH_2_O until three days after emergence (usually 4-5 days after planting). For each trial, three days after emergence, the seedlings were divided evenly into three trial groups. The first group received Hoagland nutrient solution (Supplementary Note 8) and ddiH_2_0 on alternating days. The second group received Hoagland nutrient solution in which the potassium nitrate and calcium nitrate were substituted with potassium sulfate and calcium chloride, respectively, to remove nitrate. The third group received Hoagland nutrient solution in which the monopotassium phosphate was substituted with potassium sulfate to remove phosphate. The nutrient treatments continued every other day until harvest. For the ValA treatment assay, plants grown under nutrient-optimal or N-deficit conditions were treated with 30 µM ValA dissolved in nutrient solutions beginning at 7 days after planting; the plants were treated at 6 PM every other day.

### Plant phenotyping and root sampling

On the date of harvest and phenotyping, the plants were taken to a dark room illuminated solely by green light, separated from the potting media and cleaned in a two stage process. The roots were washed in a 0.05% bleach solution and then were rinsed with warm running water and dried with paper towels. The root samples used for RNA extraction and metabolite analyses were flash frozen in liquid nitrogen. The roots were scanned using an EPSON scanner (Perfection V550, setting at 120 dpi; Epson, Suwa, Japan) with a green film covering the scanning surface to avoid exposing the roots to non-green light. Fresh biomass measurements were taken for the whole seedlings, after which they were divided into shoot and root fractions and weighted separately. Dry weight measurements of shoots and roots were taken after 48 h of freeze-drying. For paspalum, the weight of the original rhizome cutting was subtracted from the final whole-plant fresh biomass to estimate biomass accumulation.

### Species phylogeny construction

A set of 7,728 single-copy syntenic orthologs from the *Zea mays*, *Sorghum bicolor*, *Setaria italica*,*Oropetium thomaeum*, *Brachypodium distachyon* and, *Oryza sativa* genomes was extracted from the syntenic gene sets identified among the seven species. Of the 7,728 orthologs with primary transcript CDSs longer than 500 bp, 6,151 were aligned using the codon-based aligner ParaAT^120^. Subsequently, the 6,151 multiple sequence alignments, each consisting of one gene each from each of the seven species were trimmed to remove poorly aligned or highly divergent regions using Gblocks(v0.91b)^121^ with the following parameters: minimum number of sequences for a conserved position set at 5; minimum number of sequences for a flank position set at 6; maximum number of contiguous nonconserved positions set at 8; and minimum length of a block set at 10. The resulting nucleotide alignments were used to construct phylogenies using RAxML(v8.2) with the parameters ’-f a -N 1000 -m GTRGAMMA -x 1234 -p 1234’ and a clade containing rice and *Brachypodium* as an outgroup^122^. In 292 cases, it was not possible to form a monophyletic clade containing rice and *Brachypodium*. The remaining 5,859 trees were analyzed using Densitree to generate a consensus tree^123^. IQ-TREE was used to construct maximum likelihood phylogeny estimate branch lengths using a super gene concatenated from the trimmed nucleotide sequence alignments of the 5,859 single copy syntenic genes used for consensus tree analysis^124^. Divergence time estimates were then performed using these branch lengths, a previously estimated divergence date for *B. distachyon* and *O. sativa* of 54 Myr ago^125^ and an estimated divergence date for *Z. mays* and *S. bicolor* of 12 Myr ago^126^ as a reference with r8s software^127^.

### Syntenic and substitution rate analysis

Syntenic orthologous gene pairs were identified between the sorghum and paspalum genomes using sequence similarity data from LAST^128^ and a Python implementation of MCScan, JCVI^129, 130^. This analysis was run using the command ’python -m jcvi.compara.catalog ortholog paspalum sorghum – no_strip_names’. The LAST results were filtered using a Cscore setting of ≥ -0.7. Raw synteny gene pairs were polished using a previously described approach^50^. Sorghum-paspalum orthologous gene pairs were merged into a published sorghum referenced synteny list^50^ for maize (B73_RefGen_V4)^118^, sorghum v3.1^119^, foxtail millet v2.2^34^, *Oropetium* v2.0, rice v7^131^ and *Brachypodium* v3.1^132^. The final synteny list and the scripts used to generate it are hosted at https://github.com/gsun2unl/PaspalumNutrientStress.

Codon-level multiple sequence alignments of syntenic orthologous gene groups were generated with ParaAT2.0^120^. Synonymous nucleotide substitution rates (Ks), and non-synonymous nucleotide substitution rates (Ka) were estimated from these multiple sequence alignments using the ‘codeml’ package implemented in PAML^133^. The estimation was conducted using the maximum-likelihood method and the parameters runmode=0, Codon-Freq=2, model=1. The known phylogenetic relationships of the six included species were used as a known input tree. Syntenic orthologous groups containing any genes with a Ks greater than 2, a Ka greater than 0.5, and a Ka/Ks ratio greater than 2 were removed.

### Gas chromatography–mass spectrometry (GC-MS) metabolite profiling

Root samples from maize, sorghum, and paspalum seedlings grown as described above were collected in a dark room illuminated solely by a green bulb and ground into a fine powder in liquid nitrogen. Approximately 50 ± 0.5 mg of the ground powder was used for metabolite extraction and derivatization as described previously^134, 135^. A 1 µL sample of the derivatized material was analyzed in splitless mode using a 7200 GC-QTOF system (Agilent Technologies, Santa Clara, CA, USA). A solution of fatty acid methyl esters (C8 to C30) was added to each sample during derivatization to determine the retention index. The raw data were acquired using MassHunter Workstation v.08 (Agilent Technologies), while peak detection, deconvolution and identification were performed using MassHunter Unknown Analysis software (Agilent Technologies) using the Fiehn GC/MS Metabolomics RTL Library (Agilent Technologies) as a reference. Peak areas of the identified metabolites were computed using MassHunter Quantitative Analysis software (Agilent Technologies). Peak area was normalized by the precise sample fresh weight and the peak area of the ribitol added to each samples as an internal standard to calculate the relative levels of metabolites.

### Genetic map construction for genome assembly validation

Nine genetic maps generated from two populations were employed to order, and orient the scaffolds into pseudomolecules, and to validate the assembly. The first population employed was an F_1_ population of 184 individuals derived from a cross between paspalum accessions PI 509022 and HI33, previously described in Qi *et al.*^30^. The second population was generated by crossing two F1 sibs from the PI 509022 x HI33 population. Only 52 progeny of this cross were validated and ultimately used for map construction. Genotyping-by-sequencing (GBS), single nucleotide polymorphism (SNP) calling, and mapping of the F1 population were previously described^30^. Essentially the same protocols were used for marker development and genetic mapping in the F2 population, except that the restriction enzymes *PstI* and *MspI* were used for GBS library preparation. SNPs in the F1 population were called from GBS reads both independently of the genome assembly and by alignment to an early draft of the paspalum genome assembly. SNPs in the F2 population were called from GBS reads aligned to seashore paspalum assembly v2.0. Because the mapping software MAPMAKER^136^ does not have an algorithm to deal with outcrossing species, the three sets of SNPs were further split into HA sets (comprising markers heterozygous in the female parent and homozygous in the male parent), AH sets (homozygous in the female parent and heterozygous in the male parent) and HH sets (heterozygous in both parents), leading to a total of six F1 datasets^30^ and three F2 datasets (Supplementary Note 2). For the F2 population, information from the grandparents was used to rescore the progeny using the rules listed in Table S2 to ensure that all markers were in the same linkage phase.

To assist with scaffold ordering and assessment of the quality of the assembly, 500 bp on either side of mapped SNP markers were excised from the assembly used for GBS read alignment and mapped to consecutive improved versions of the assembly using BLASTN. The sequences and location of the mapped F1 and F2 markers on the seashore paspalum version 3.0 assembly reported here as determined by the top BLASTN hit are provided in Supplementary Note 2. Discrepancies between marker orders in any two of the nine maps and the order and orientation of scaffolds in the pseudomolecules triggered manual examination and in some cases error correction

### Gene family analysis in various crop species

Protein sequences of the primary transcripts for seven species were retrieved from Phytozome: *Zea mays*, *Sorghum bicolor*, *Setaria italica*, *Paspalum vaginatum*, *Oropetium thomeaum*, *Brachypodium distachyon*, and *Oryza sativa*^137^. These sequences were used as inputs for orthoFinder^138, 139^ to generate clusters of genes representing gene families. Family expansion and contraction were determined with CAFE5 using default settings^140^. Significantly expanded gene families in paspalum were defined as those with significantly different lambda value (p <0.05) that showed increases in gene copy numbers in the lineage leading to paspalum as estimated by CAFE5^140^.

### RNA isolation, sequencing, and quantification

Root samples were homogenized by grinding to a fine powder in liquid nitrogen. Approximately 50 mg of homogenized root tissue per sample was mixed with 1 mL of TRIzol reagents by robust vortexing and, incubated at room temperature (25°*C*) for 10 minutes. The samples were mixed with 200 µL chloroform and incubated for 15 minutes at room temperature until a clear separation of three layers was observed. The tubes containing the mixtures were centrifuged at 12,000 rpm for 15 minutes to achieve phase separation. The top layer was transferred to a new set of tubes containing 400 µL isopropanol and incubated on ice for at least 30 minutes. RNA precipitation was achieved by centrifugation at 12,000 rpm for 15 minutes at 4°*C*. Following the removal of the supernatants, the precipitates were washed with 75 % ethanol three times before being dissolved in 40 µL of 65°*C* DEPC treated water.

The quality of individual RNA samples was assessed using an Agilent 2100 Bioanalyzer. Samples with RNA Integrity Number (RIN) values >5 were used to isolate mRNA and construct RNA sequencing libraries using a TrueSeq v2 kit from Illumina^141^. Paired-end sequence data (2x75 bp) were generated using an Illumina NextSeq 500 platform. The overall quality of the RNAseq reads was assessed using FASTQC^142^ (Figure S4A). Demultiplexed reads were filtered and quality trimmed using Trimmomatic (v0.33) with the parameters “-phred33 LEADING:3 TRAILING:3 slidingwindow:4:15 MINLEN:36 ILLUMINACLIP:TruSeq3-PE.fa:2:30:10”.^143^. Trimmed reads were mapped to the reference genomes of their respective species using STAR/2.7^144^ with two rounds of mapping; the first round of mapping was run with the parameters “–alignIntronMin 20 –alignIntronMax 20000 –outSAMtype None –outSJfilterReads Unique –outSJfilterCountUniqueMin 10 3 3 3 –outSJfilterCountTotalMin 10 3 3 3” and the second round of mapping was run after a new genome index was built based on the known and novel splicing sites recognized by the first round of mapping with parameters “–alignIntronMin 20 –alignIntronMax 20000 –limitBAMsortRAM 5000000000 –outSAMstrandField intronMotif –alignSJoverhangMin 20 –outSAMtype BAM SortedByCoordinate”. Maize reads were mapped to B73_RefGen_V4^118^. Sorghum reads were mapped to v3.1 of the BTx623 reference genome downloaded from Phytozome^119^. Paspalum reads were mapped to the paspalum genome assembly described and released as a part of this paper. A Transcripts Per Million (TPM) table was generated using Kallisto^145^. Syntenic orthologous genes across paspalum, maize and sorghum with a mean TPM value higher than 50 were log transformed prior to principal component analysis (Figure S4B). For each individual sequencing library, the read counts were determined using the software package HTSeq (version 0.9) with the parameter settings “-r pos -s no -t exon -i gene_id”, the overlap mode used was the default (“union”)^146^. Statistically significant DEGs were identified from the read count matrix generated by HTSeq using DESeq2 (v1.22.2)^147^(Figure S4B). Genes were considered to be significantly differentially expressed when an absolute log_2_ fold change >1 and an adjusted p value lower than 0.05 were both observed. Total RNA of paspalum shoot was extracted using the same method and sequenced using the same library preparation protocol and sequencing platform as other samples described in this study, only genes with TPM higher than 5 and syntenically conserved were examined. Statistical significance of expression level changes was calculated by DESeq2^147^.

### MDC staining of samples and confocal microscopy

Microscopy visualization of autophagosomes by Monodansylcadaverine (MDC) staining was performed as described by Contento et al. (2005)^148^. Root tissues from maize seedlings one week after germination were gently rinsed with sterilized ddiH_2_O and submerged in 40 µM MDC solution for 30 minutes in the dark. Confocal microscopy imaging was performed with a Nikon A1 laser scanning confocal mounted on a Nikon 90i compound microscope (software version: NIS Elements 4.13). Excitation/emission for MDC detection was set to 488 nm/505–550 nm. Aperture and light source intensity were kept the same for all images taken. MDC stained autophagosomes from ten different cells in root tips were counted. Three biological samples were examined.

### Gene ontology enrichment analysis

Gene ontology (GO) enrichment analysis of the DEGs was performed using GOATOOLS^149^. To ensure consistency in cross species comparisons, the same population of syntenically conserved genes in maize, sorghum, and paspalum was used as the population set for enrichment analysis in each species. Similarly, to avoid bias introduced by the use of different GO term annotation pipelines, the same set of GO terms was assigned to each syntenic ortholog in each of the three species. These annotations were taken from the GO terms assigned to the maize copy of each conserved syntenic gene group by Maize-GAMER^150^. As the whole genome duplication in maize introduced bias into the background gene set (genes retained as duplicate homeologous gene pairs are enriched in the annotations transcription factor, “responds to X” and protein complex subunit) only a single copy from maize1 subgenome of each maize gene pair was retained for both the background population set and the DEG defined set.

### Immunoblot detection of free ATG8 and ATG8-PE conjugate

ATG8 and ATG8-PE conjugate were detected as previously described with slight modifications^151^. Maize seedlings were grown under full-nutrient or N-deficient conditions for three weeks with or without the 30 µM validamycin A treatment described above. Root tissues were collected in a dark room solely illuminated by green light and ground to a fine powder in liquid nitrogen. The ground root tissues were homogenized in lysis buffer (50 mM Tris-HCl, pH 8.0, 150 mM NaCl, 1 mM phenylmethylsulfonyl fluoride, 10 mM iodoacetamide, and 1 X complete protease inhibitor cocktail [Sigma Aldrich, St. Louis, MO, USA)]) and centrifuged at 2000 X*g*, 4°*C* for 5 min. The extracted protein samples were quantified using a Bradford assay, and 25 µg protein was loaded onto a 15% SDS-PAGE (polyacrylamide gel electrophoresis) gel containing 6 M urea. Immunoblotting was performed with affinity-purified anti-At ATG8 antibodies (1:1000 dilution)(Agrisera, Vännäs, Sweden; AS14 2769). The ATG8-PE (lipidation) band was confirmed by incubating protein samples at 37°*C* for 1 hour with *Streptomyces chromofuscus* phospholipase D (Thermo Fisher Scientific, Waltham, MA, USA; 525200-250U; 250 units mL^-1^ final concentration) as previously described^78^.

## Data availability statement

The genome sequence and annotation is accessible via Phytozome v13: https://phytozome-next.jgi.doe.gov/info/Pvagin RNAseq data for root tissues of paspalum, maize and sorghum under three nutrient conditions are available at NCBI under the BioProject: PRJNA746310. RNAseq data for root tissues of maize seedlings under three nutrient conditions with or without validamycin A treatment are available at NCBI under the BioProject: PRJNA746310. RNAseq data for paspalum shoots/rhizome is available at NCBI with SRA id SRR10230104; SRR10230108; SRR10230122; SRR10230130. RNAseq data of maize wild type and atg12 mutant is available at NCBI with Accession ID: PRJNA449498. Illumina sequence of papspalum genome is available at NCBI with Accession ID: PRJNA234783. All of the scripts and raw data used for figures can be accessed at github: https://github.com/gsun2unl/PaspalumGenome

## Acknowledgements

This manuscript is based upon work supported by award 2016-67013-24613 from the USDA National Institute of Food and Agriculture to JCS, by National Science Foundation Award OIA-1826781 to JS and JCS and OIA-1557417 to CZ, TO, BS, BY and JCS, by National Health Institute Award GM127414 to BY, and by USDA-SCRI award UFDSP00011194 to KMD. This work was completed utilizing the Holland Computing Center of the University of Nebraska, which receives support from the Nebraska Research Initiative. The work in this manuscript was enabled by the UNMC DNA Sequencing Core Facility which receives partial support from the Nebraska Research Network In Functional Genomics NE-INBRE P20GM103427-14, The Molecular Biology of Neurosensory Systems CoBRE P30GM110768, The Fred & Pamela Buffett Cancer Center - P30CA036727, and the Nebraska Research Initiative. The work conducted by the U.S. Department of Energy Joint Genome Institute is supported by the Office of Science of the U.S. Department of Energy under Contract no. DE-AC02-05CH11231. We thank the Plant Editors for their work and feedback on an early draft of the manuscript.

## Author contribution

J. C. S., J. S., and G. S., conceived this research. J. A. S., and G. S., designed and directed the study. T. O., N. W., and L. B., profiled primary metabolites. C. C., L. S., Y. Y., C. D., K. B., and R. O directed and performed paspalum genome sequencing. J. S., C.P., and J. J., directed and performed paspalum genome assembly. K. D., P. Q., and T. G., constructed genetic maps used for genome assembly. S. S., performed the annotation of the genome assembly. B. Y., and B. Z., designed and conducted the quantification of ATG8 and ATG8-PE. C. Z., A.L. and H. Y assisted with the RNAseq analysis. B. S., designed and conducted flow cytometry experiments. J. C. S., and G. S., drafted the manuscript. The final version of the manuscript was generated with input and contributions from N. W., S. S., J. J., B. Z., P. Q., H. Y., C. Z., K. D., B. S., B. Y., T. O., J. S., All authors approved the final version of the manuscript.

---

**Table S1.** Final summary assembly statistics for chromosome scale assembly

|  |  |
| --- | --- |
| Scaffold total | 1903 |
| Contig total | 2212 |
| Scaffold sequence total | 651.0 Mb |
| Contig sequence total | 648.0 Mb (0.5% gap) |
| Scaffold L/N50 | 7 / 44.5 Mb |
| Contig L/N50 | 111/ 1.5 Mb |
| Number of scaffolds >50 Kb | 1112 |
| % main genome in scaffolds >50 Kb | 95.5 % |

**Table S2.** Rescoring of progeny based on allele information from grandparents

| Grandparent 1 | Grandparent 2 | Parents | Rule (to be applied to the F2 progeny) | Marker suffix |
| --- | --- | --- | --- | --- |
| A | H | AH or HA | Keep original scores | - |
| H | A | AH or HA | Change 'A' to 'H', and 'H' to 'A' | R |
| H | H | AH or HA | Duplicate marker; keep original scores in one copy (marker suffix 'u'), change 'A' to 'H' and 'H' to 'A' in second copy (marker suffix 'ur') | u / ur |
| A | B | HH | Keep original scores | - |
| A | H | HH | Keep original scores | - |
| H | B | HH | Keep original scores | - |
| B | A | HH | Change 'A' to 'B', and 'B' to 'A' | r |
| H | A | HH | Change 'A' to 'B', and 'B' to 'A' | r |
| B | H | HH | Change 'A' to 'B', and 'B' to 'A' | r |
| H | H | HH | Duplicate marker; keep original scores in one copy (marker suffix 'u'), change 'H' to '-' and both 'A' and 'B' to 'H' in second copy (marker suffix 'uD') | u / uD |

**Figure S1.**
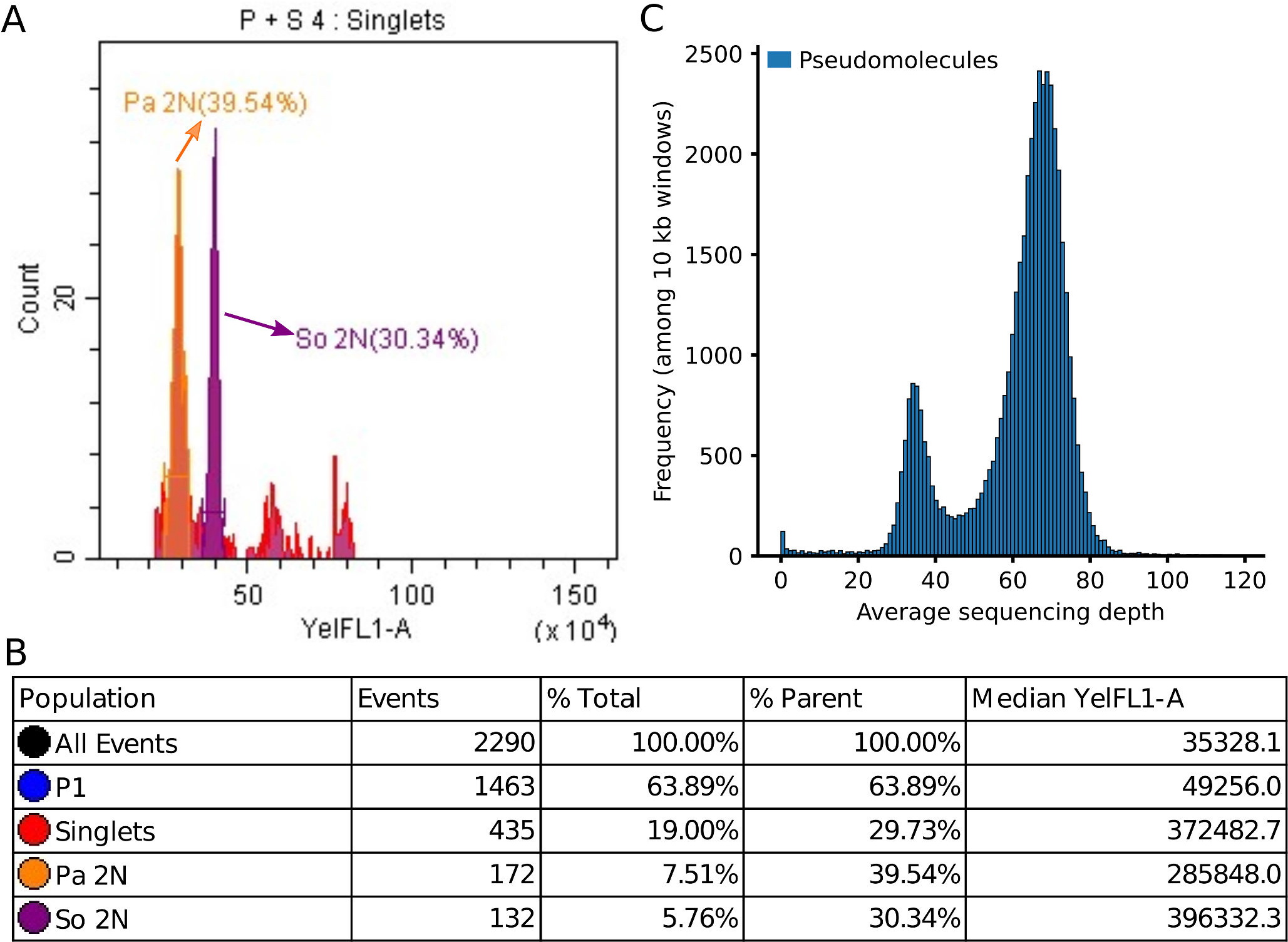
Estimating the genome size of the paspalum accession employed for genome sequencing. (A) Estimation of genome size using flow cytometry. The x axis indicates yellow fluorescence intensity, which is linearly correlated with the genome size. *Sorghum bicolor* BTX623 nuclei (So 2N) were used as an internal control. (B) Statistics of the flow cytometry results for one representative sample. Based on the median yellow fluorescence intensity, the ratio of genome size of *Paspalum vaginatum* (Pa) to *Sorghum bicolor*(So) is 285848:396332. The Sorghum bicolor genome size is 818 Mbp^108^; therefore, the estimated genome size of *Paspalum vaginatum* is 590 Mbp. (C) Genome wide read coverage of Illumina sequencing reads mapped to the current paspalum genome assembly.

**Figure S2.**
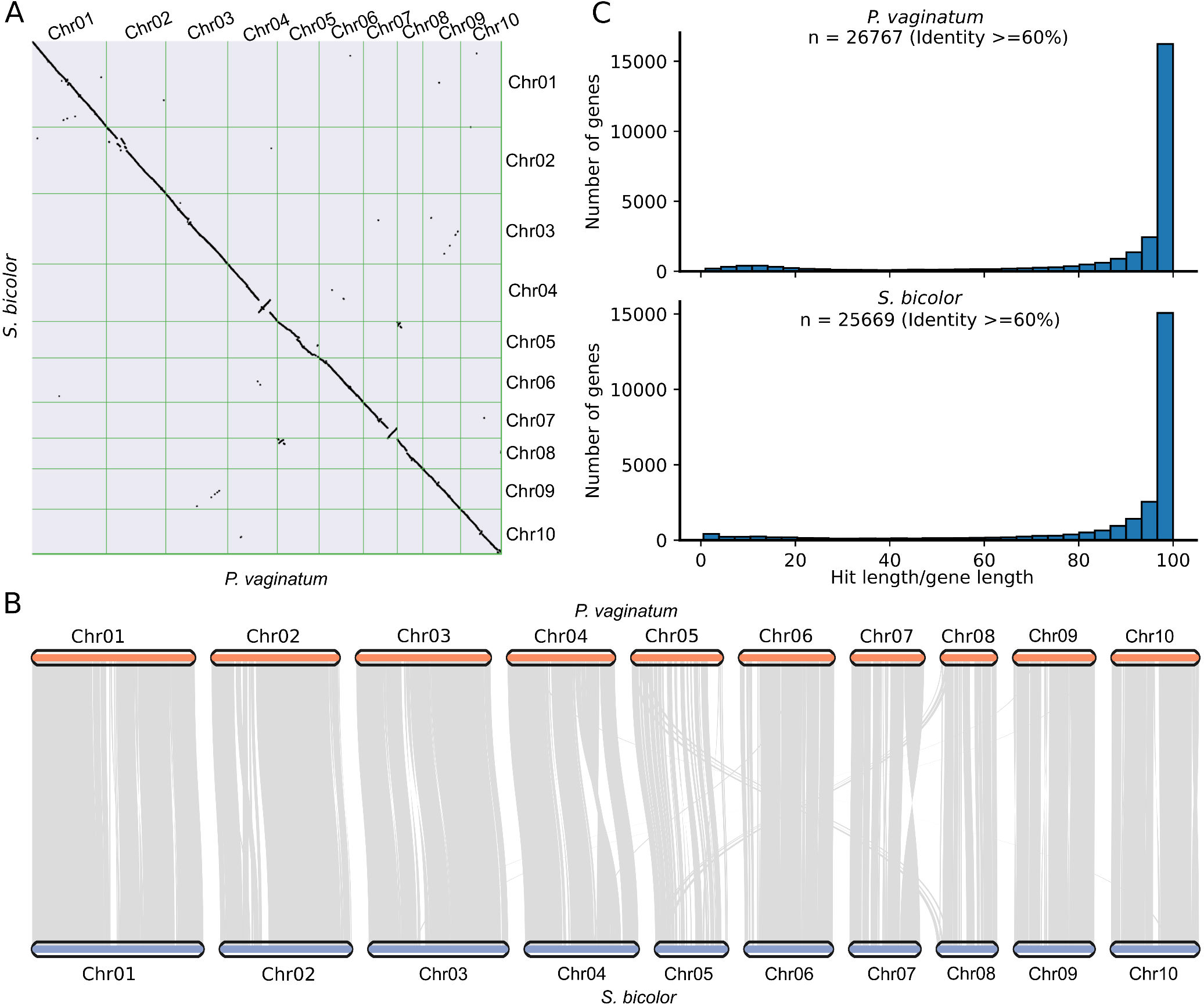
Gene families identified among the seven grass species. (A-B) Syntenic regions conserved between the paspalum genome and sorghum genome. (C) Length of homologs with identity >60% over the annotated length of proteins annotated in paspalum genome when BLAST paspalum proteome against sorghum proteome (upper) and reversely, the length of homologs with identity >60% over the annotated length of proteins annotated in sorghum genome when BLAST sorghum proteome against paspalum proteome (lower).

**Figure S3.**
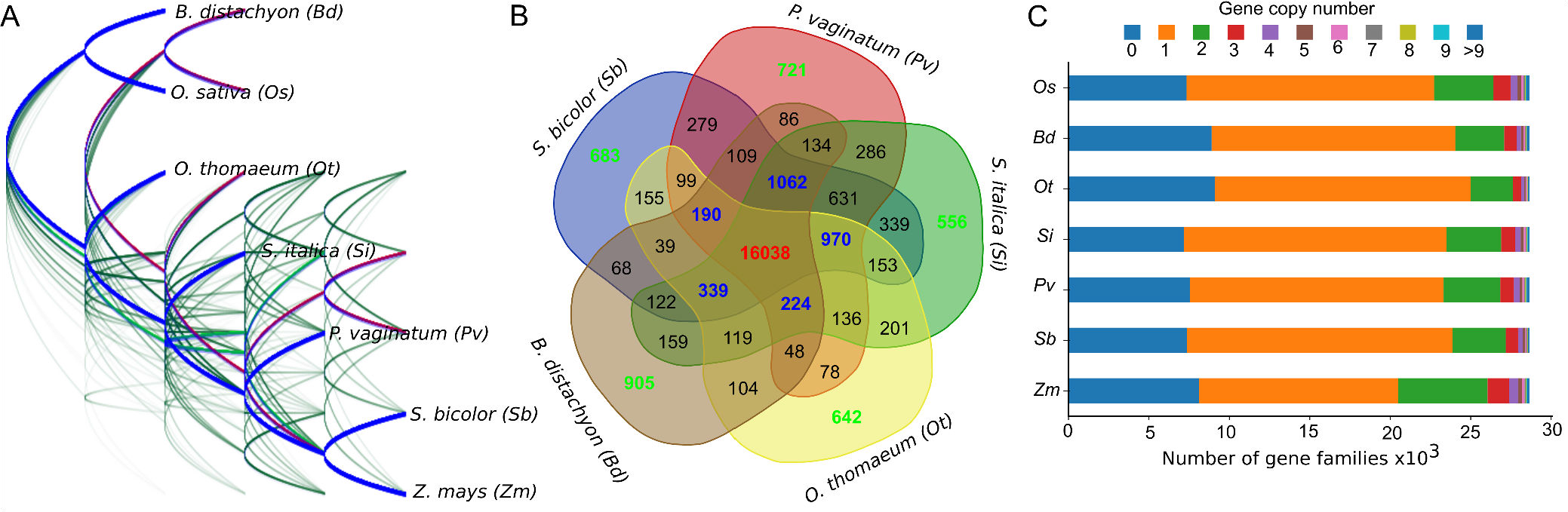
Gene families identified among the seven grass species. (A) DensiTree drawn from phylogenies constructed based on selected individual single-copy sytenic orthologous gene pairs across species: paspalum (*Paspalum vaginatum*), maize (*Zea mays*), sorghum (*Sorghum bicolor*), foxtail millet (*Setaria italica*), *Oropetium* (*Oropetium thomaeum*), *Brachypodium* (*Brachypodium distachyon*), and rice (*Oryza sativa*). The consensus tree drawn in blue was supported by 4,265 (73%) of the individual gene trees and the second most common topology drawn in purple was supported by 762 individual gene trees (13%). (B) Comparison of shared and species-specific gene families among the five grass species. Green numbers indicate species-specific gene families. Blue numbers indicate gene families shared by all but one of the five species compared, while numbers in red indicate the number of gene families shared across all five species. Gene families shared by either two or three of the five species are shown in black. Maize and rice which do not have a unique most recent common ancestor (MRCA) with paspalum (the MRCA of maize and paspalum is the MRCA of sorghum and pasaplum, and the MRCA of rice and pasplaum is the MRCA of *Brachypodium* and paspalum), were omitted to simplify visualization. (C) Distribution of copy numbers for gene families in each of the seven species shown in panel A.

**Figure S4.**
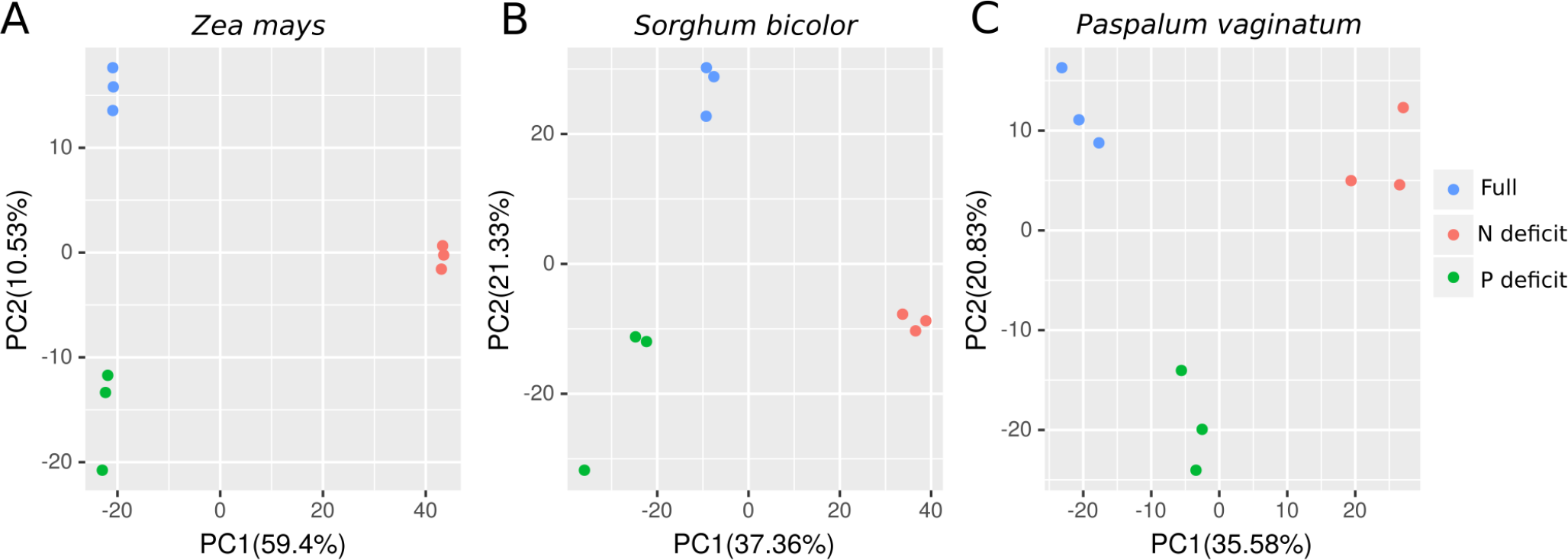
Principal component analysis of biological replicates of root transcriptomes under three experimental nutrient conditions. (A) Principal component analysis based on log transformed expression of syntenic genes in maize (*Zea mays*) (B) Principal component analysis based on log transformed expression of syntenic genes in sorghum (*Sorghum bicolor*) (C) Principal component analysis based on log transformed expression of syntenic genes in paspalum (*Paspalum vaginatum*). Panels (A-C) Nutrient conditions are color coded. PC, principal component.

**Figure S5.**
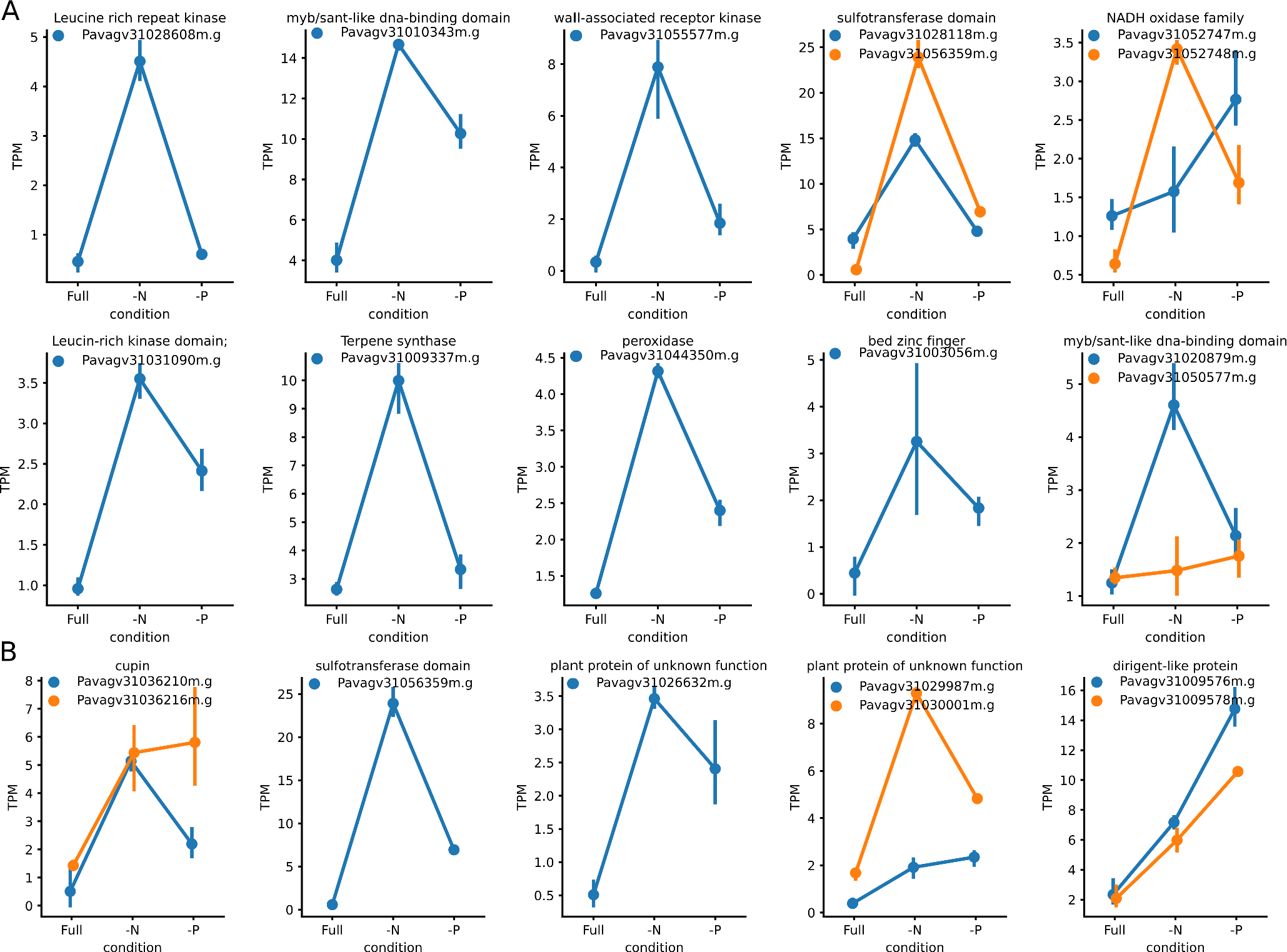
Members of gene families that are transcriptionally responsive to nutrient-deficit conditions. (A) Members of paspalum-specific expanded gene families that are transcriptionally responsive to nitrogen deficiency. (B) Members of paspalum-specific expanded gene families that are transcriptionally responsive to phosphorus deficiency.

**Figure S6.**
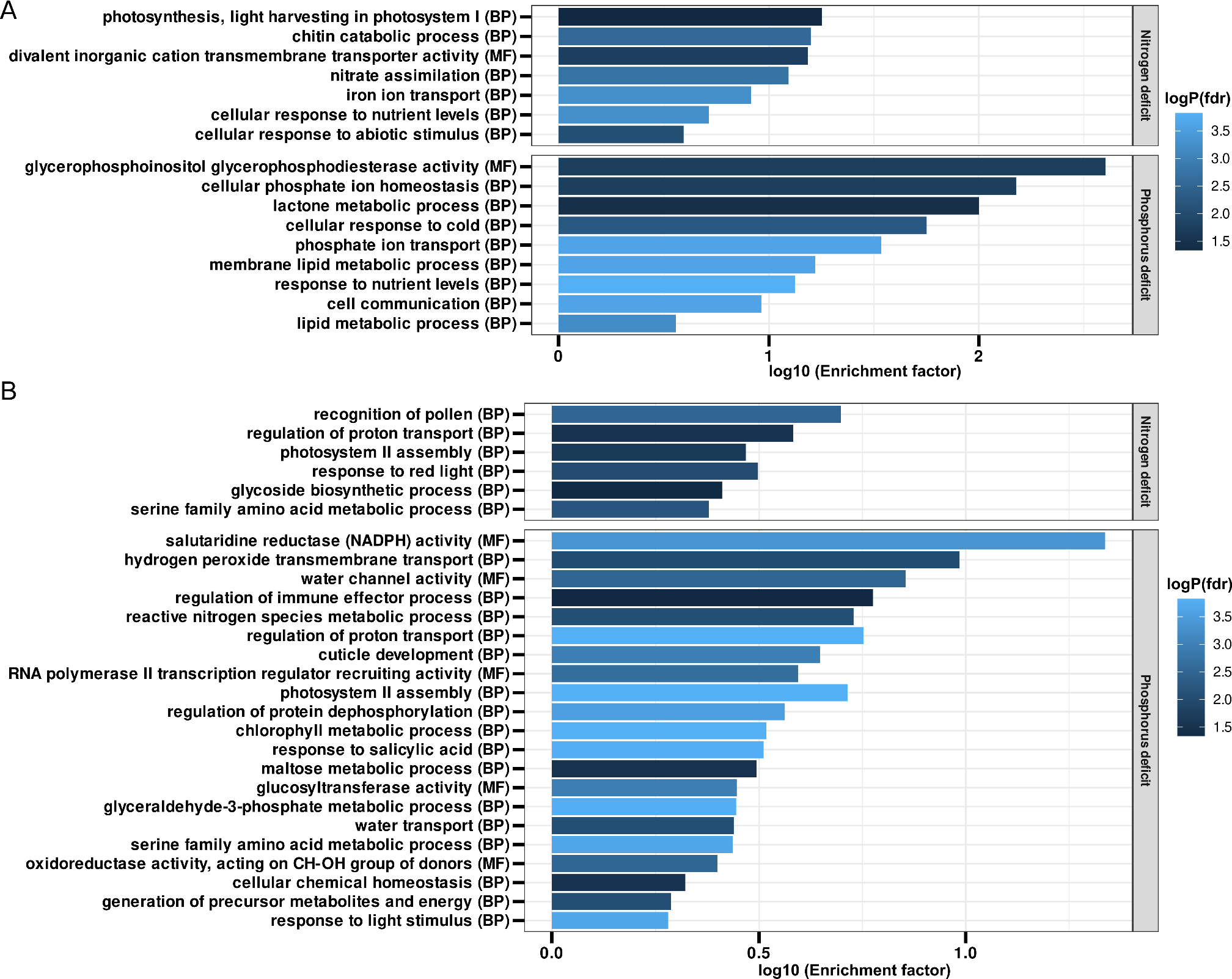
Gene ontology (GO) analysis of differentially expressed syntenic orthologous genes across the three species and in papspalum alone. (A) Significantly enriched GO terms (false discovery rate (FDR) ≤ 0.05) for 220 and 37 syntenic orthologous genes that were differentially expressed in all of the three species in response to N-deficit and P-deficit conditions, respectively. Bars indicate the log-transformed enrichment factor (number of genes associated with the overrepresented GO terms in the study gene set over the number of genes associated with the GO term in the background gene set) for enriched GO terms. Negative log-transformed multi-test corrected p values are color coded. (B) Significantly enriched GO terms (false discovery rate (FDR) ≤ 0.05) in 825 and 650 syntenic orthologous genes that were differentially expressed only in paspalum in response to N-deficit and P-deficit conditions, respectively. Bars indicate the log-transformed enrichment factor (number of genes associated with the overrepresented GO terms in the study gene set over the number of genes associated with the GO term in the background gene set) for enriched GO terms. Negative log-transformed multi-test corrected p values are color coded.

**Figure S7.**
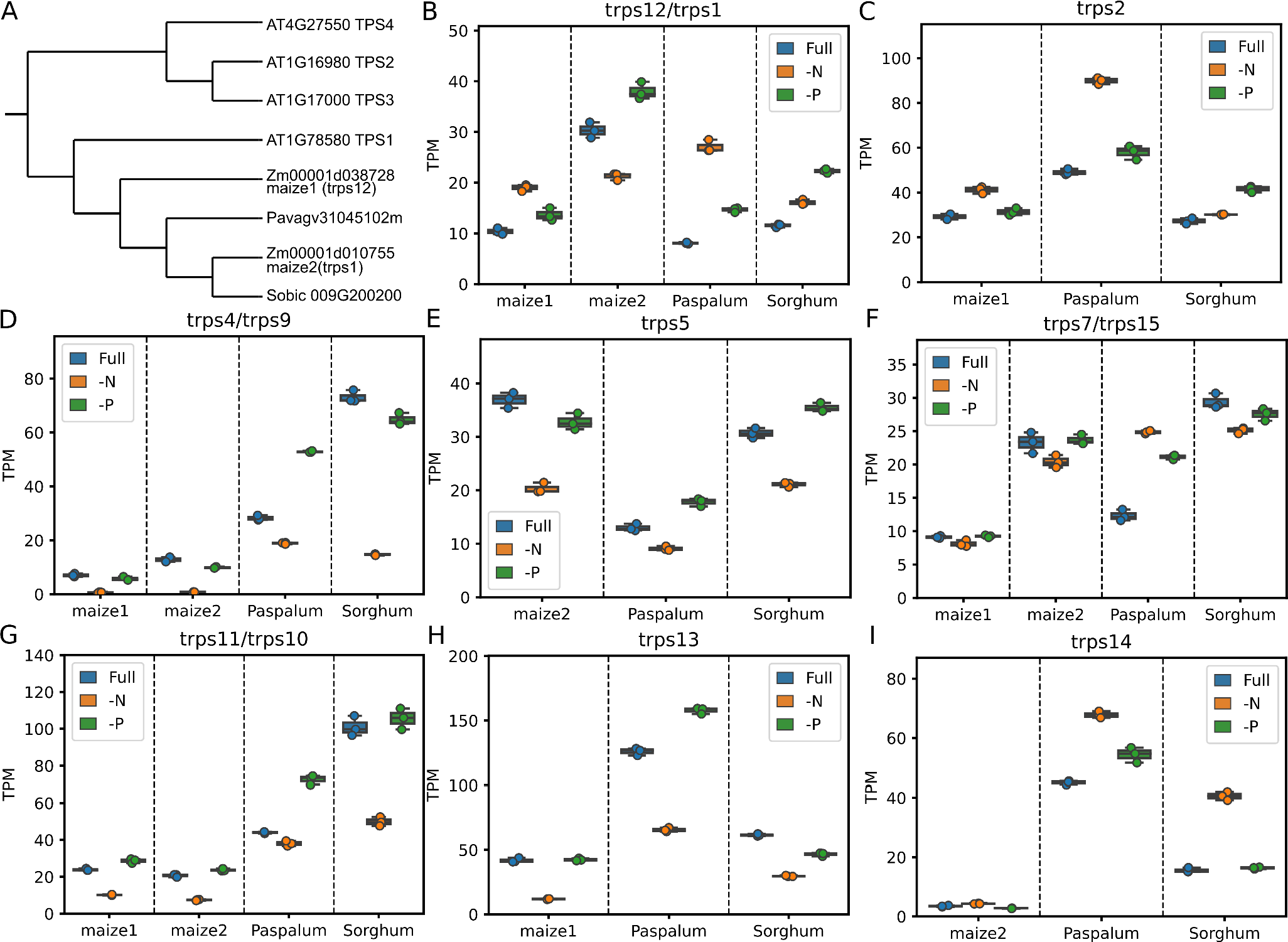
Expression patterns of genes encoding trehalose-6-phosphate synthase in response to nutrient stress across maize (*Zea mays*), sorghum (*Sorghum bicolor*), and paspalum (*Paspalum vaginatum*). (A) Phylogeny of orthologs of Arabidopsis trehalose-6-phosphate synthase 1 (*TPS1*) genes in the three species. (B) Expression pattern of the trehalose-6-phosphate synthase genes in the three species under nutrient-optimal (Full), nitrogen-deficit (–N), and phosphorus-deficit (–P) conditions. “maize1” and “maize2” indicate the two subgenomes that formed in maize after the recent whole-genome duplication event 12–16 million years ago. (C-I) Expression patterns of other syntenic genes annotated as encoding trehalose-6-phosphate synthase that did not cluster with Arabidopsis homologs in the three species under nutrient-optimal (Full), nitrogen-deficit (–N), and phosphorus-deficit (–P) conditions.

**Figure S8.**
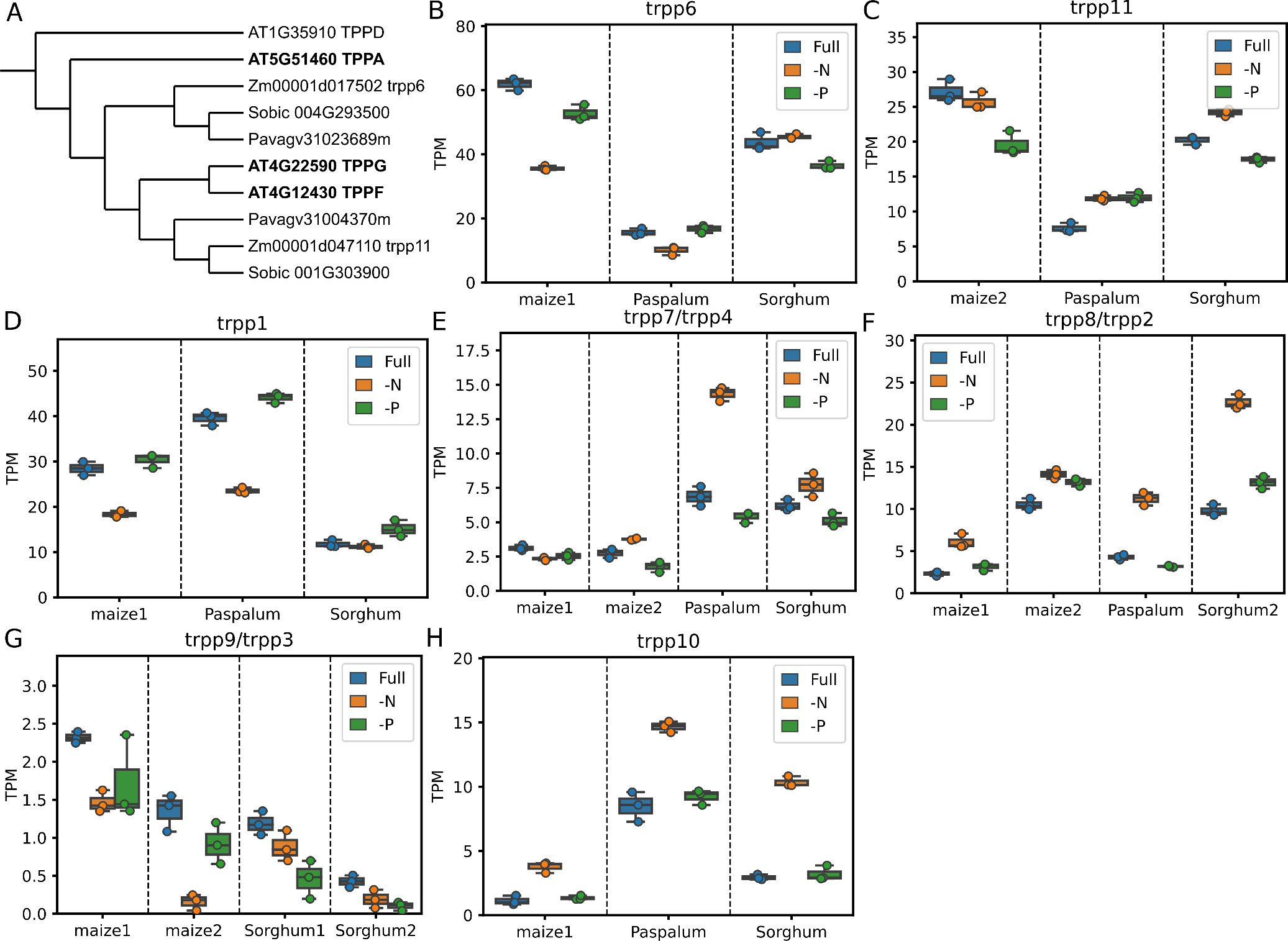
Expression patterns of genes encoding trehalose-6-phosphate phosphatase enzymes in response to nutrient stress across maize (*Zea mays*), sorghum (*Sorghum bicolor*), and paspalum (*Paspalum vaginatum*). (A) Phylogeny of orthologs of characterized Arabidopsis trehalose-6-phosphate phosphatase (TPP) genes in the three species. (B-C) Expression patterns of the trehalose-6-phosphate phosphatase genes (*trpp6* and *trpp11*) that clustered with their Arabidopsis homologous (*TPPA*, *TPPG*, *TPPF*) under nutrient-optimal (Full), nitrogen-deficit (–N), and phosphorus-deficit (–P) conditions. (D-H) Expression patterns of other syntenic genes annotated as trehalose-6-phosphate phosphatase that did not cluster with Arabidopsis homologs in the three species grown under nutrient-optimal (Full), nitrogen-deficit (–N), and phosphorus-deficit (–P) conditions. For panels B to I, “maize1” and “maize2” indicate the two subgenomes that formed in maize after the recent whole-genome duplication event 12–16 million years ago.

**Figure S9.**
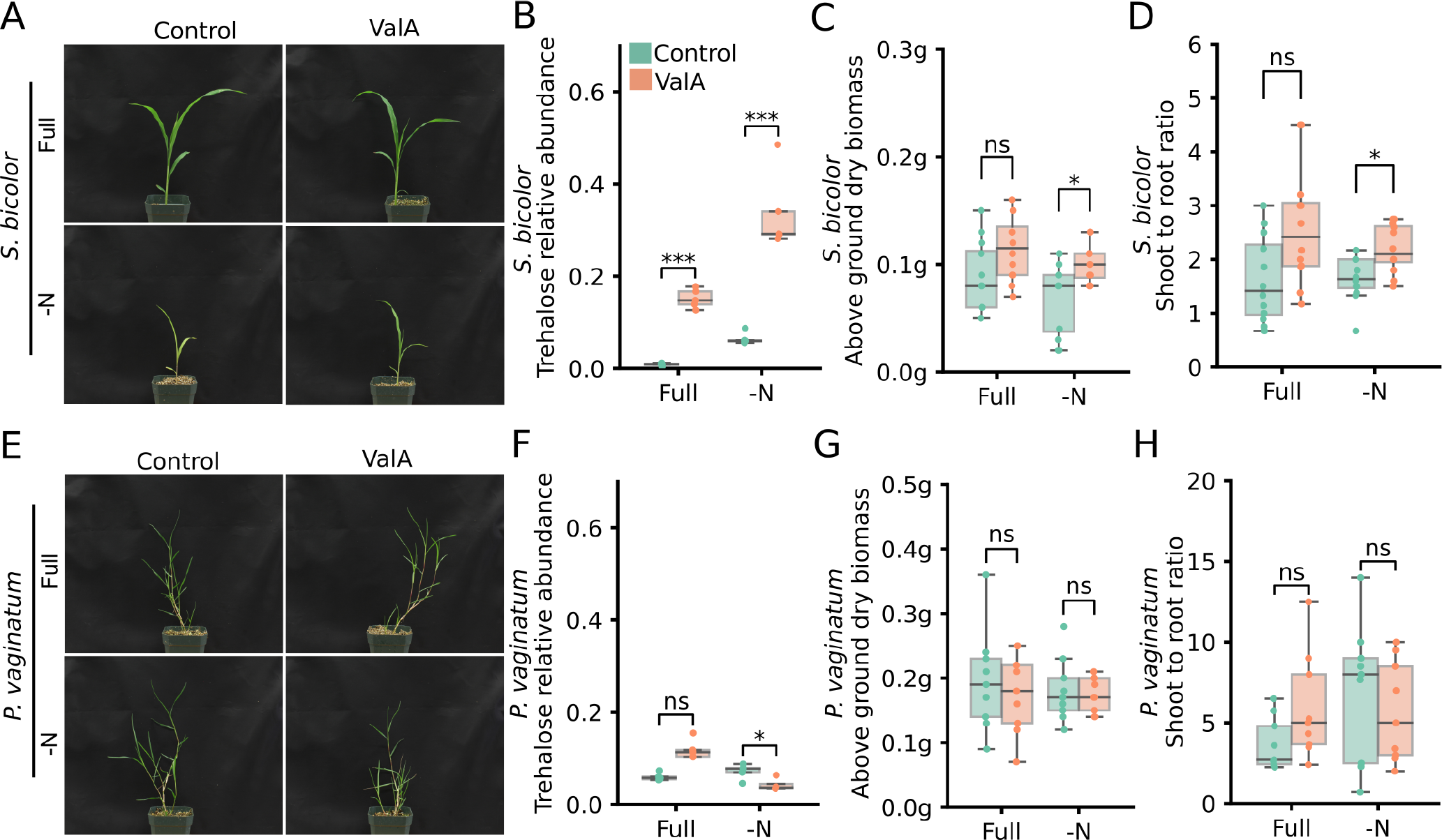
ValA treatment alters biomass accumulation and nutrient reallocation to sorghum (*Sorghum bicolor*) grown under nutrient-deficient conditions. (A) Representative images of sorghum seedlings grown under nutrient optimal and N-deficit conditions with or without validamycin A (ValA) treatment. Images were taken 21 days after planting. For the ValA treatment, a 30 µM solution was added at 6 PM on the day that the plants were watered with the indicated nutrient solutions. (B) Changes in observed trehalose abundance – normalized to an internal reference (ribitol) – in response to validamycin A and/or nutrient conditions in sorghum root tissues. Error bars are standard deviations. **Student’s t-test** (* = p <**0.05**; ** = p <**0.005**; *** = p <**0.0005**). (C) Dry weight of the above-ground tissue of sorghum seedlings grown under nutrient-optimal and nitrogen-deficit conditions harvested at 3 weeks after planting. Plant tissues were freeze-dried for 48 hours after harvesting. (D) Shoot-to-root ratio calculated from the dry weight of above-ground tissues and roots of the same sorghum seedlings. (E) Representative images of paspalum seedlings at 3 weeks after planting grown under nutrient optimal (Full) and nitrogen-deficient (-N) conditions with (ValA) or without (Control) validamycin A treatment. (F) Lack of significant increases in trehalose abundance (normalized to an internal reference [ribitol]) in response to validamycin A treatment (ValA) in 3-week-old paspalum seedlings under either full-nutrient or N-deficient conditions. (G) No significant change observed in above ground dry weight of 3-week-old paspalum seedlings in response to validamycin A treatment (ValA) under full-nutrient or N-deficient conditions. (H) Ratio of shoot-to-root dry weight in 3-week-old paspalum seedlings grown with or without validamycin A under full-nutrient or N-deficient conditions.

**Figure S10.**
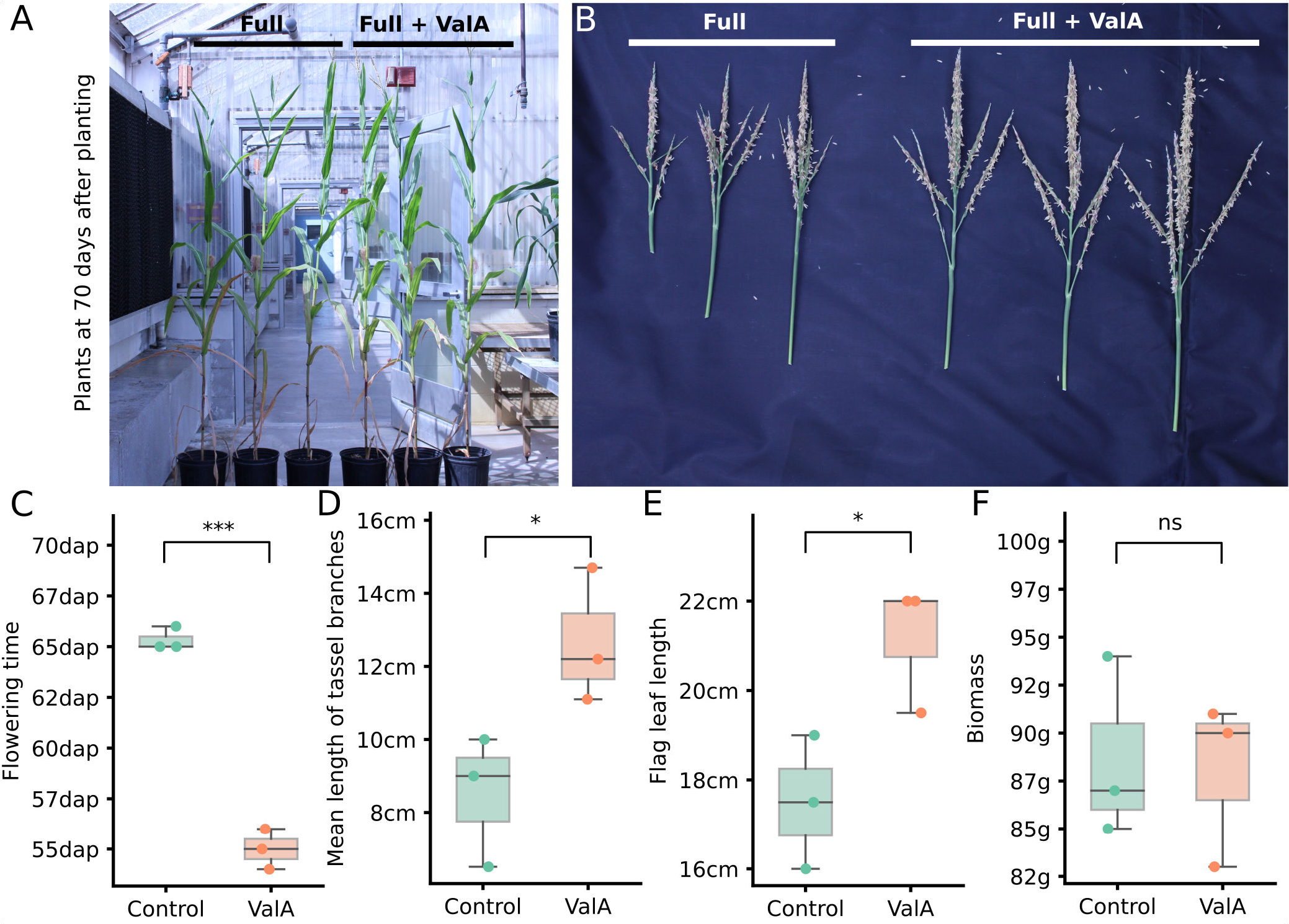
Validamycin A treatment improves important agronomic traits in adult maize plants. (A-B) Images of whole plants (A) and tassels (B) taken 70 days after planting. The left three plants were grown under full-nutrient conditions and the right three were grown under full-nutrient conditions with a weekly 30 µM ValA treatment. (C-F) Flowering time (C), mean length of tassel branches (D), flag leaf length (E) and above ground dry biomass (F) of plants with or without ValA treatment. **Student’s t-test** (* = p <**0.05**; ** = p <**0.005**; *** = p <**0.0005**).

**Figure S11.**
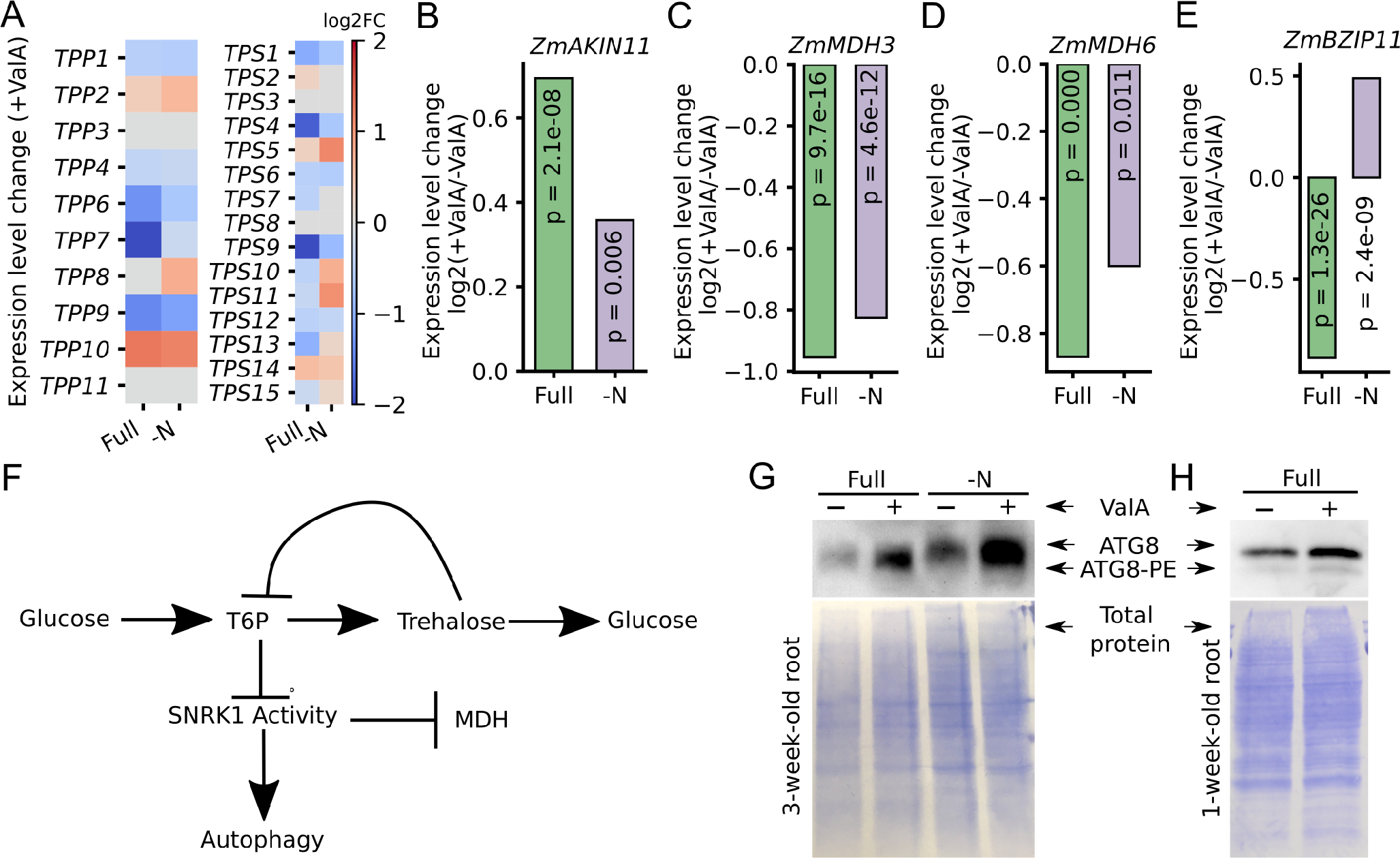
Transcriptional responses of SNRK1 target genes to Validamycin A treatment. (A) Expression level fold changes of the genes encoding trehalose-6-phosphate synthases (TPS) and trehalose-6-phosphate phosphatases (TPP) relative to the seedlings treated with validamycin A under Full and -N conditions. (B-E) Expression level fold changes of the *ZmAKIN2* (A), *ZmMDH3* (B), *ZmMDH6* (C) and *ZmBZIP11* with our without validamycin A treatment under full nutrient and N-deficient conditions. p values were calculated by DESeq2 after correction for false discovery rate lower than 0.05. (F) Trehalose accumulation might lead to a lower T6P level, resulting in the release of inhibition of SNRK1 activity. The active status of SNRK1 would promote autophagy and *ZmAKIN11* expression while repressing the expression of MDH and bZIP genes. (G-H) A biological replicate of the immunoblot measuring the abundance of both free ATG8 (upper band) and the ATG8-PE conjugate (lower band) in root samples collected from 3-week-old maize seedlings grown under optimal nutrient (Full) and nitrogen-deficit (-N) conditions with or without ValA treatment (G) and in root samples collected from 1-week-old maize seedlings grown under optimal nutrient conditions with or without ValA treatment (H). Total protein loading control is shown in the lower panel.

Supplementary Notes attached to this submission as separate files.

- Supplementary Note 1: Detailed paspalum genome assembly and annotation methods.
- Supplementary Note 2: Markers used for paspalum genetic map construction.
- Supplementary Note 3: Calculated Ka, Ks, and Ka/Ks ratios for each grass gene employed in this study.
- Supplementary Note 4: Genes from the paspalum specific expanded gene families and GO terms enriched among these genes.
- Supplementary Note 5: Raw fold change values for each metabolite plotted in Figure 3.
- Supplementary Note 6: Phylogeny of TRPP homologues across arabidopsis, maize, sorghum and paspalum.
- Supplementary Note 7: Phylogeny of TRPS homologues across arabidopsis, maize, sorghum and paspalum.
- Supplementary Note 8: Recipes for full and modified hoagland solutions employed in this study.

## Notes

### Competing Interest Statement

James C. Schnable has equity interests in Data2Bio, LLC; Dryland Genetics LLC; and EnGeniousAg LLC. He is a member of the scientific advisory board of GeneSeek and currently serves as a guest editor for The Plant Cell.

