## Supplementary note 1 for "The genome of stress tolerant crop wild relative *Paspalum vaginatum* leads to increased biomass productivity in the crop *Zea mays*"

Paspalum genome assembly and annotation supplemental methods:

### *Sequencing*

We sequenced *Paspalum vaginatum* using a whole genome shotgun sequencing strategy and standard sequencing protocols. Sequencing reads were collected using Illumina and PacBio platforms. Illumina and PacBio reads were sequenced at Department of Energy (DOE) Joint Genome Institute (JGI) in Walnut Creek, California. Illumina reads were sequenced using the Illumina HiSeq-2500 platform, and the PacBio reads were sequenced using the SEQUEL I platform. One 400bp insert 2x150 Illumina fragment library (102.4x) was used (See Table S1). Prior to assembly, Illumina fragment reads were screened for phix contamination. Reads composed of >95% simple sequence were removed. Illumina reads <50bp after trimming for adapter and quality (q<20) were removed. The final read set consists of 266,209,446 reads for a total of 102.4x of high quality Illumina bases. For the PacBio sequencing, a total of 14 PB chemistry 2.0 chips (10 hour movie time) were sequenced with a sequence yield of 52.01 Gb, with a total coverage of 74.30x (See Table S2).

### *Genome assembly and construction of pseudomolecule chromosomes*

The version 1.0 assembly was generated by assembling the 5,012,142 PacBio reads (74.30x sequence coverage) using the MECAT assembler (Xiao et al., 2017) and subsequently polished using QUIVER (Chin et al., 2013). This produced an initial assembly of 5,358 scaffolds (5,358 contigs), with a contig N50 of 771.9 Kb, 1,099 scaffolds larger than 100 Kb, and a total assembled size of 838.4 Mb (Table S3).

Two genetic maps totaling 7,680 markers (provided by the Katrien Devos group, University of Georgia), combined with primary transcripts from the version 2 release of *Panicum hallii* var. HAL2 were used to identify misjoins in the initial assembly. Misjoins were characterized as a discontinuity in the *P. vaginatum* linkage group. A total of 15 misjoins were identified and resolved. The resulting broken contigs were then oriented, ordered, and joined together into 10 chromosomes using both the map, as well as the *P. hallii* primary transcripts. A total of 347 joins were made during this process. Each chromosome join is padded with 10,000 Ns. Significant telomeric sequence was identified using the $(TTTAGGG)_n$ repeat, and care was taken to make sure that it was properly oriented in the production assembly. The remaining scaffolds were screened against bacterial proteins, organelle sequences, GenBank nr and removed if found to be a contaminant.

Heterozygous snp/indel phasing errors were corrected using the 74.3x raw PACBIO data. A total of 102,526 (4.2% of the 2,317,409 heterozygous SNPS/InDels) were corrected. Finally, homozygous SNPs and InDels were corrected in the release consensus sequence using 78x of Illumina reads (2x150, 400bp insert) by aligning the reads using bwa mem (Li, 2013) and identifying homozygous SNPs and InDels with the GATK's UnifiedGenotyper tool (McKenna et al., 2010). A total of 1,591 homozygous SNPs and 233,333 homozygous InDels were corrected in the release. The final version 1.0 release contains 652.7 Mb of sequence, consisting of 2,274 contigs with a contig N50 of 1.5 Mb and a total of 75.0% of assembled bases in the 10 chromosomes.

**Construction of the scaffold assembly**

A total of 5,012,142 PacBio reads (74.30x) were assembled using MECAT assembler (Xiao et al., 2017), and formed the starting point of the version 1.0 release. The 266,209,446 Illumina fragment reads (102.40x sequence coverage) was used for fixing homozygous snp/indel errors in the consensus.

| Library | Sequencing Platform | Average Read/Insert Size | Read Number | Assembled Sequence Coverage (x) |
|---|---|---|---|---|
| USGX | Illumina | 400 | 266,209,446 | 102.4 |
| | PacBio | 9,523* | 5,012,142 | 74.30 |
| **Total** | | N/A | 271,221,588 | 176.70 |

**Table S3.1.** Genomic libraries included in the *Paspalum vaginatum* genome assembly and their respective assembled sequence coverage levels in the final release.

*Average read length of PacBio reads.

| Cutoff | Number of Reads | Basepairs | Average Read Length | Coverage |
|---|---|---|---|---|
| 0 | 5,021,142 | 52,011,227,922 | 9,523 | 74.30x |
| 1,000 | 4,598,012 | 51,803,193,177 | 10,358 | 74.00x |
| 2,000 | 4,284,360 | 51,337,470,200 | 10,963 | 73.34x |
| 3,000 | 4,011,480 | 50,660,511,262 | 11,490 | 72.38x |
| 4,000 | 3,772,204 | 49,824,505,385 | 11,967 | 71.19x |
| 5,000 | 3,545,694 | 48,806,129,176 | 12,435 | 69.72x |
| 6,000 | 3,326,274 | 47,599,675,919 | 12,908 | 68.00x |
| 7,000 | 3,100,356 | 46,130,347,947 | 13,419 | 65.91x |
| 8,000 | 2,871,056 | 44,410,763,109 | 13,968 | 63.44x |
| 9,000 | 2,637,734 | 42,426,702,674 | 14,563 | 60.61x |
| 10,000 | 2,390,849 | 40,080,198,809 | 15,239 | 57.26x |
| 11,000 | 2,132,551 | 37,367,664,499 | 16,009 | 53.38x |
| 12,000 | 1,878,010 | 34,441,849,335 | 16,844 | 49.20x |
| 13,000 | 1,642,413 | 31,498,764,753 | 17,705 | 45.01x |
| 14,000 | 1,429,052 | 28,620,342,486 | 18,575 | 40.89x |
| 15,000 | 1,238,057 | 25,852,785,076 | 19,449 | 36.93x |
| 16,000 | 1,067,747 | 23,214,688,325 | 20,327 | 33.16x |
| 17,000 | 916,859 | 20,726,586,386 | 21,209 | 29.61x |
| 18,000 | 784,038 | 18,403,847,127 | 22,088 | 26.29x |
| 19,000 | 666,673 | 16,233,960,809 | 22,979 | 23.19x |

**Table S3.2.** PacBio library statistics for the libraries included in the *Paspalum vaginatum* genome assembly and their respective assembled sequence coverage levels.

| Minimum Scaffold Length | Number of Scaffolds | Number of Contigs | Scaffold Size | Basepairs | % Non-gap Basepairs |
|---|---|---|---|---|---|
| 5 Mb | 4 | 4 | 30,869,566 | 30,869,566 | 100.00% |
| 2.5 Mb | 54 | 54 | 202,593,315 | 202,593,315 | 100.00% |
| 1 Mb | 165 | 165 | 377,268,994 | 377,268,994 | 100.00% |
| 500 Kb | 296 | 296 | 472,159,561 | 472,159,561 | 100.00% |
| 250 Kb | 444 | 444 | 524,257,651 | 524,257,651 | 100.00% |
| 100 Kb | 1,009 | 1,009 | 604,765,225 | 604,765,225 | 100.00% |
| 50 Kb | 3,282 | 3,282 | 755,766,926 | 755,766,926 | 100.00% |
| 25 Kb | 5,279 | 5,279 | 836,810,963 | 836,810,963 | 100.00% |
| 10 Kb | 5,358 | 5,358 | 838,436,451 | 838,436,451 | 100.00% |
| 5 Kb | 5,358 | 5,358 | 838,436,451 | 838,436,451 | 100.00% |
| 2.5 Kb | 5,358 | 5,358 | 838,436,451 | 838,436,451 | 100.00% |
| 1 Kb | 5,358 | 5,358 | 838,436,451 | 838,436,451 | 100.00% |
| 0 bp | 5,358 | 5,358 | 838,436,451 | 838,436,451 | 100.00% |

**Table S3.3**. Summary statistics of the initial output of the QUIVER polished MECAT assembly. The table shows total contigs and total assembled basepairs for each set of scaffolds greater than the size listed in the left hand column.

**Plots of Marker Maps**

**Plots of the marker placements for the 10 chromosomes are shown in Figures S3.1-S3.10.**

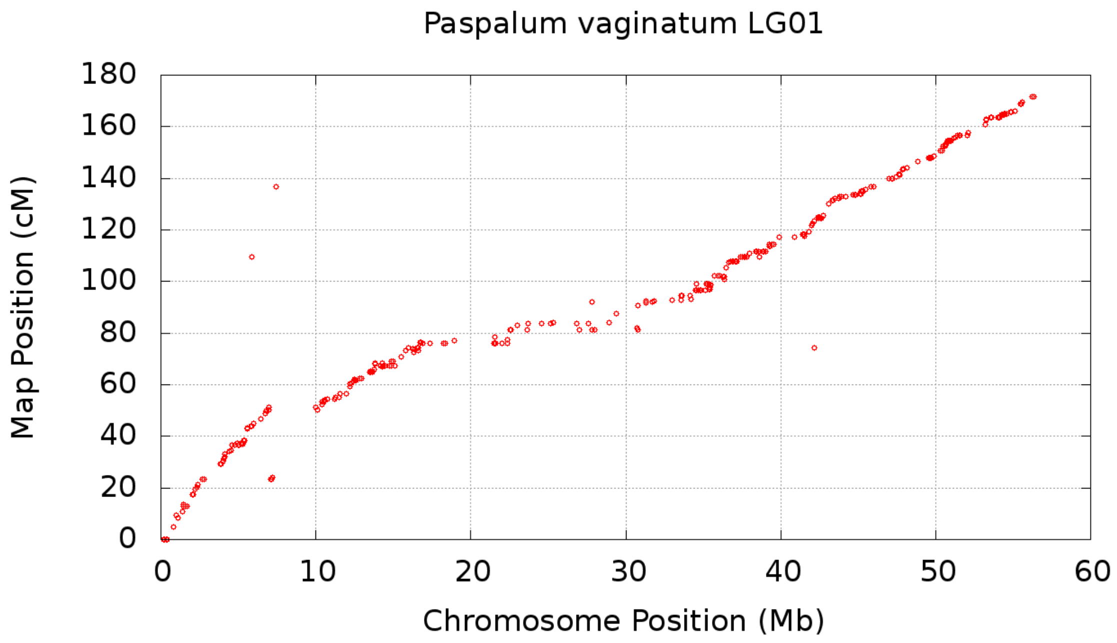


**Figure S3.1:** Marker map placements on the *Paspalum vaginatum* chromosome 1.

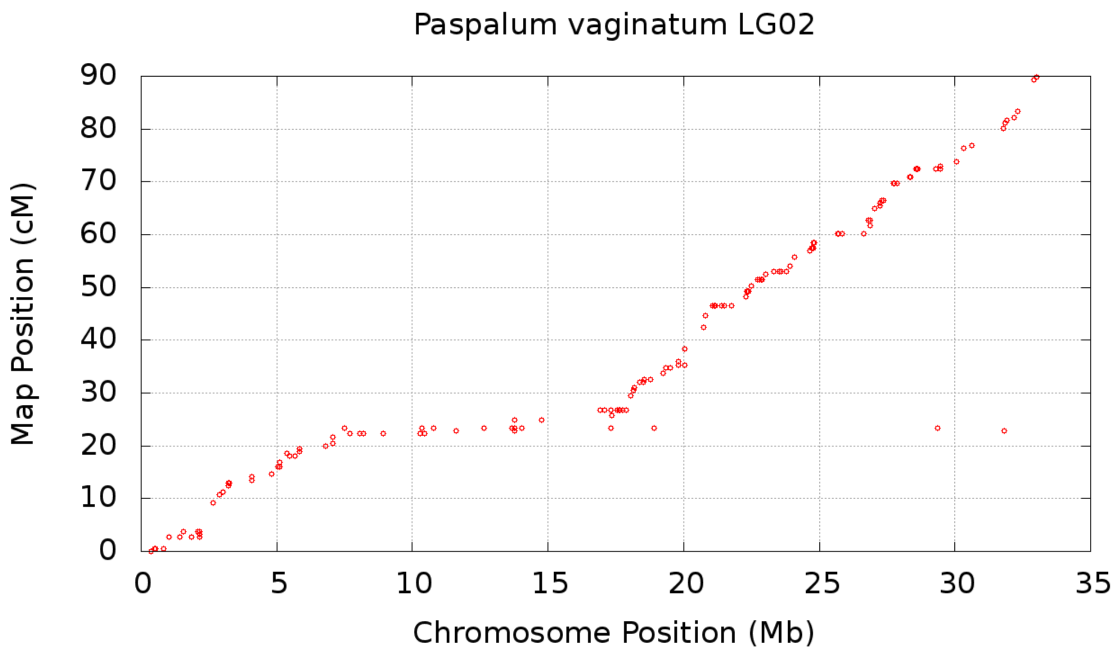


**Figure S3.2:** Marker map placements on the *Paspalum vaginatum* chromosome 2.

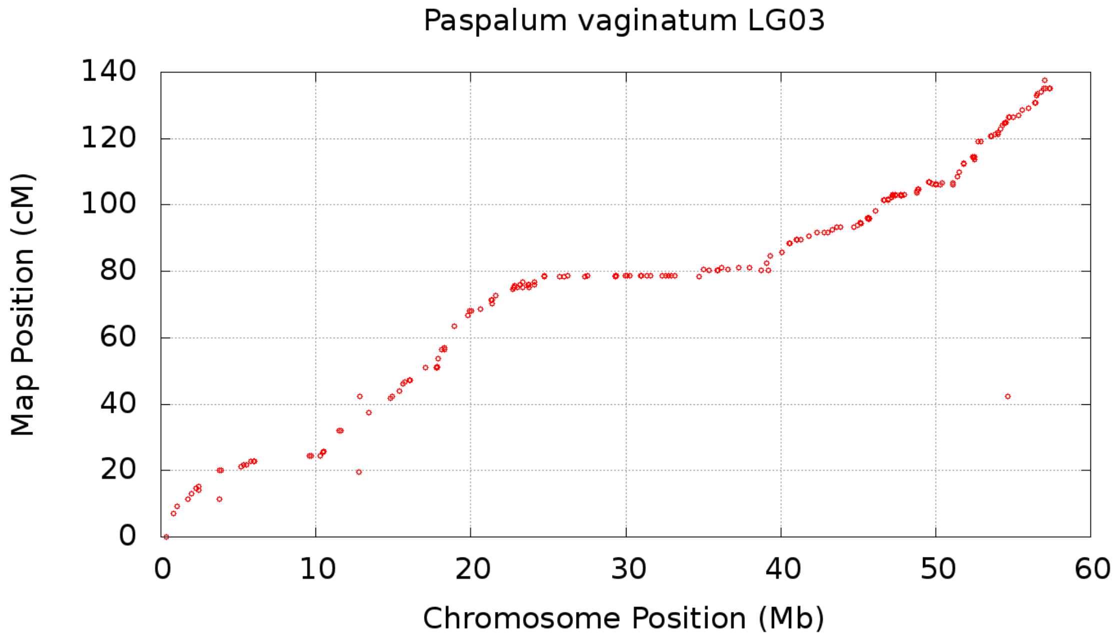


**Figure S3.3:** Marker map placements on the *Paspalum vaginatum* chromosome 3.

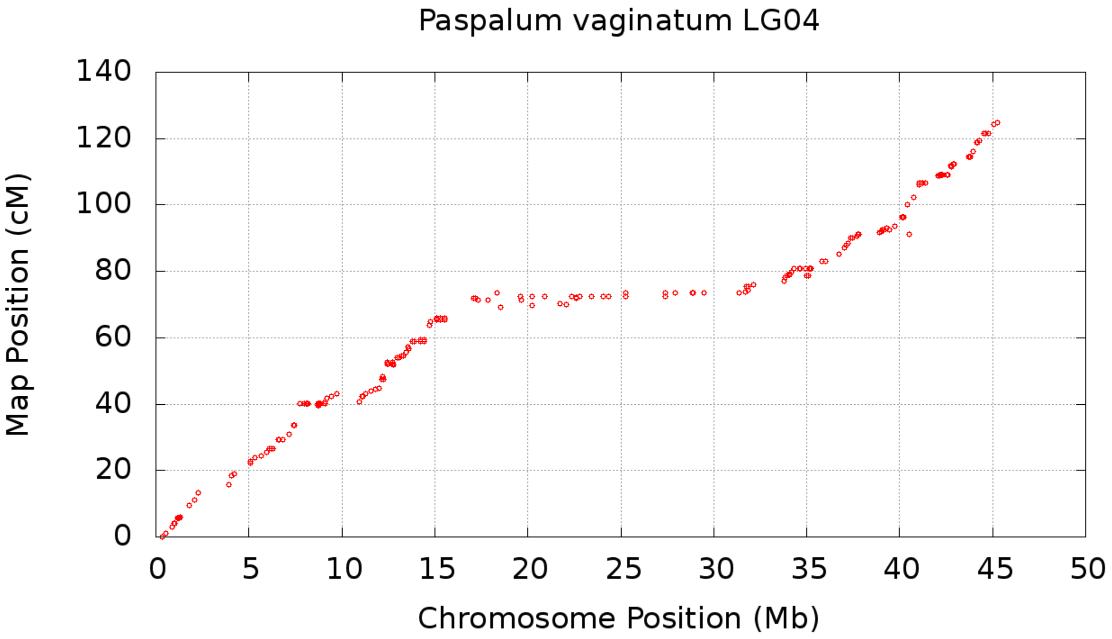


**Figure S3.4:** Marker map placements on the *Paspalum vaginatum* chromosome 4.

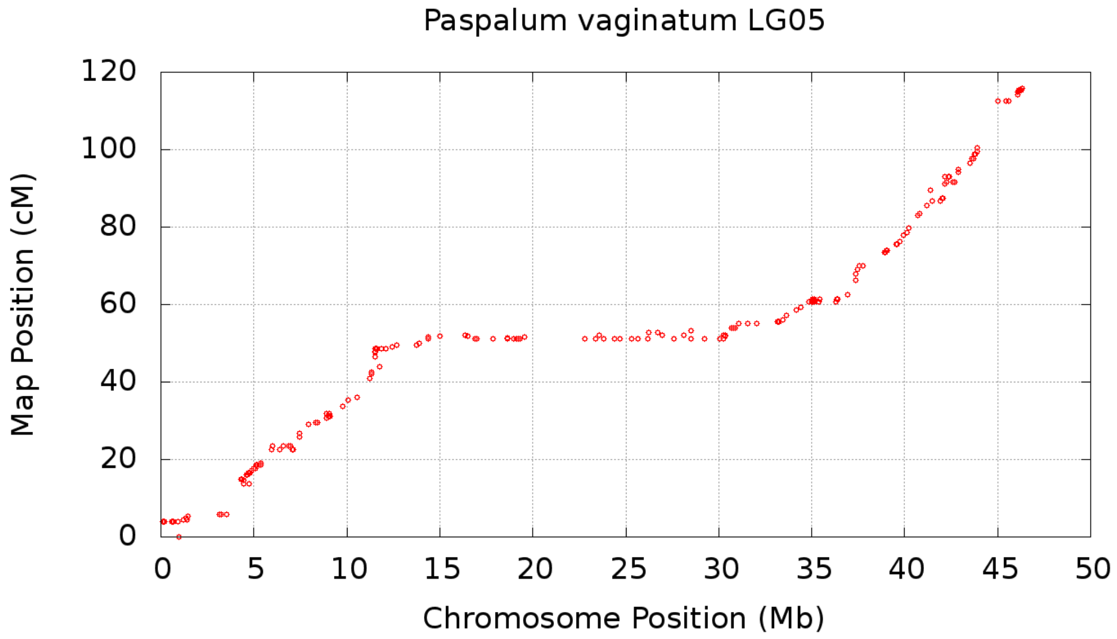


**Figure S3.5:** Marker map placements on the *Paspalum vaginatum* chromosome 5.

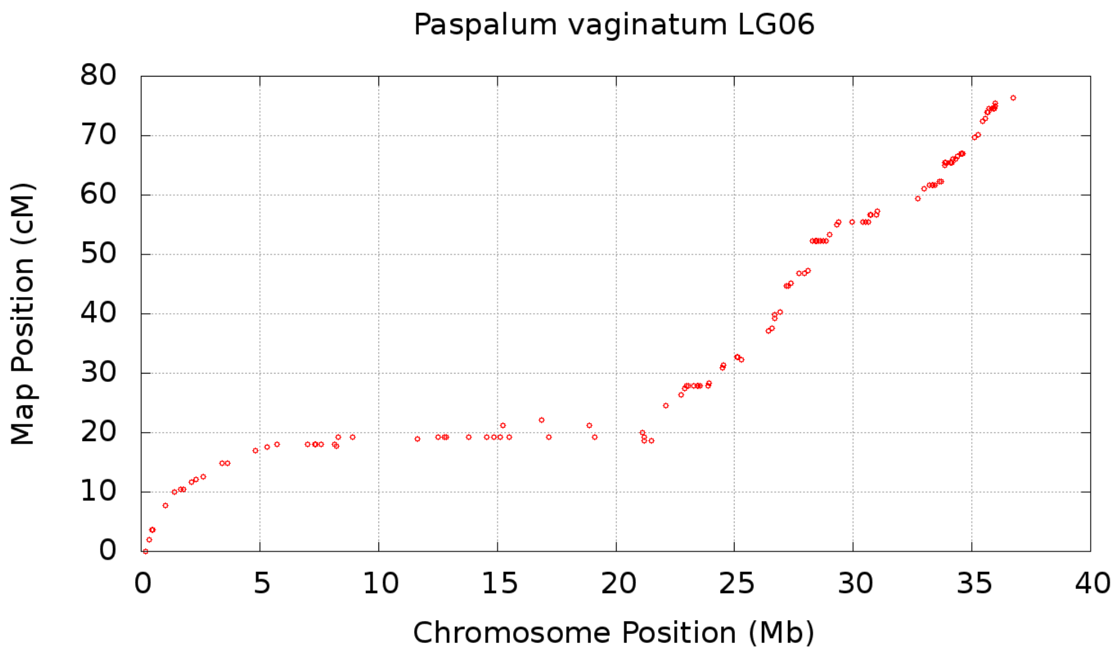


**Figure S3.6:** Marker map placements on the *Paspalum vaginatum* chromosome 6.

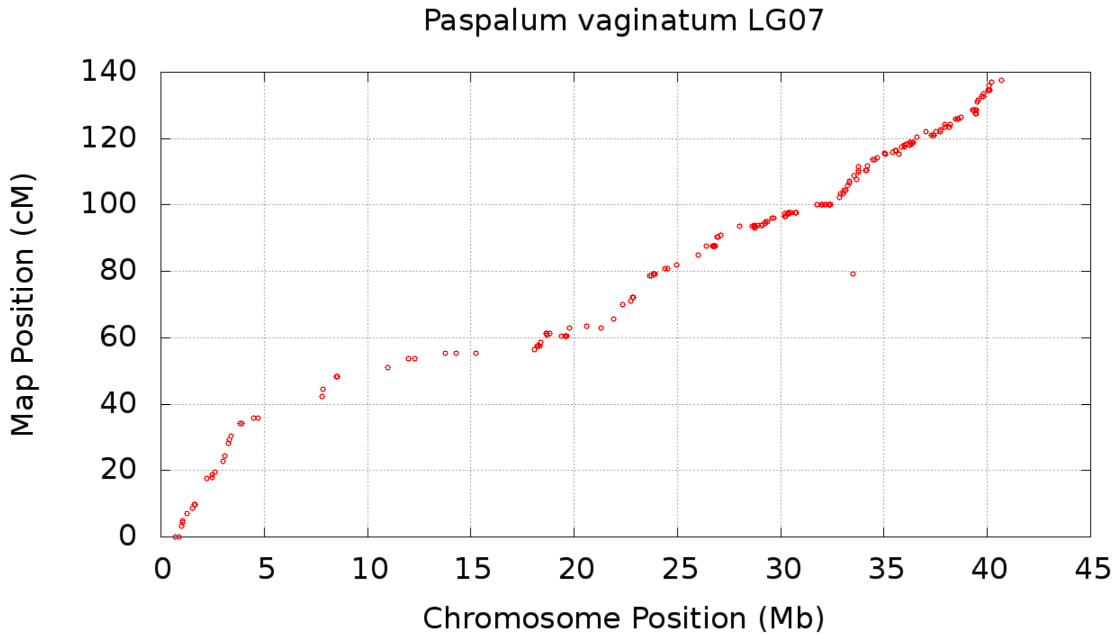


**Figure S3.7:** Marker map placements on the *Paspalum vaginatum* chromosome 7.

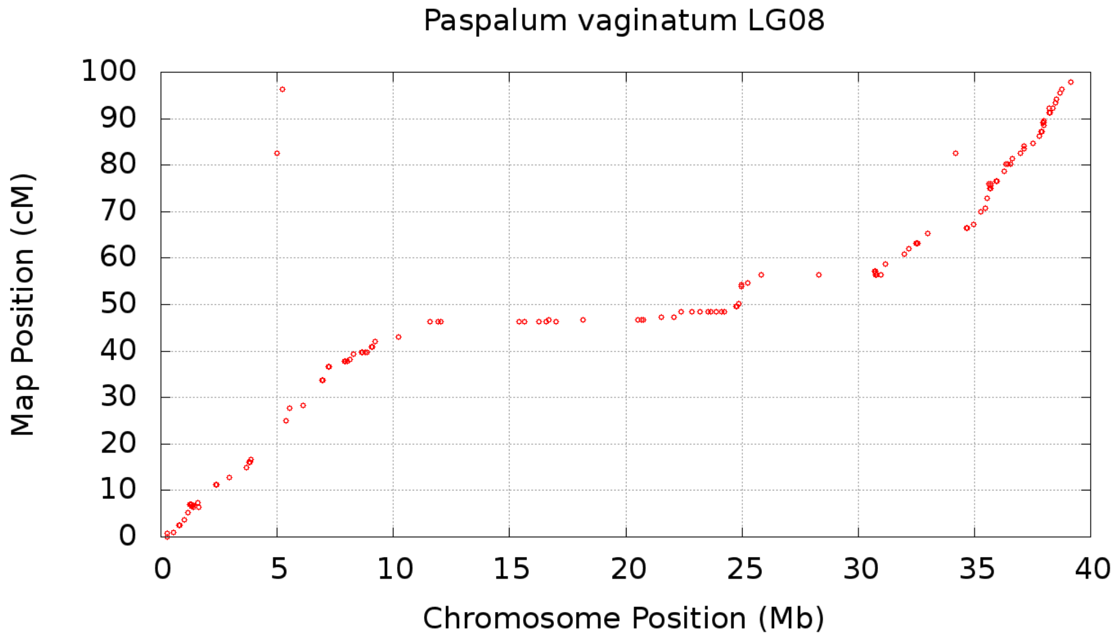


**Figure S3.8:** Marker map placements on the *Paspalum vaginatum* chromosome 8.

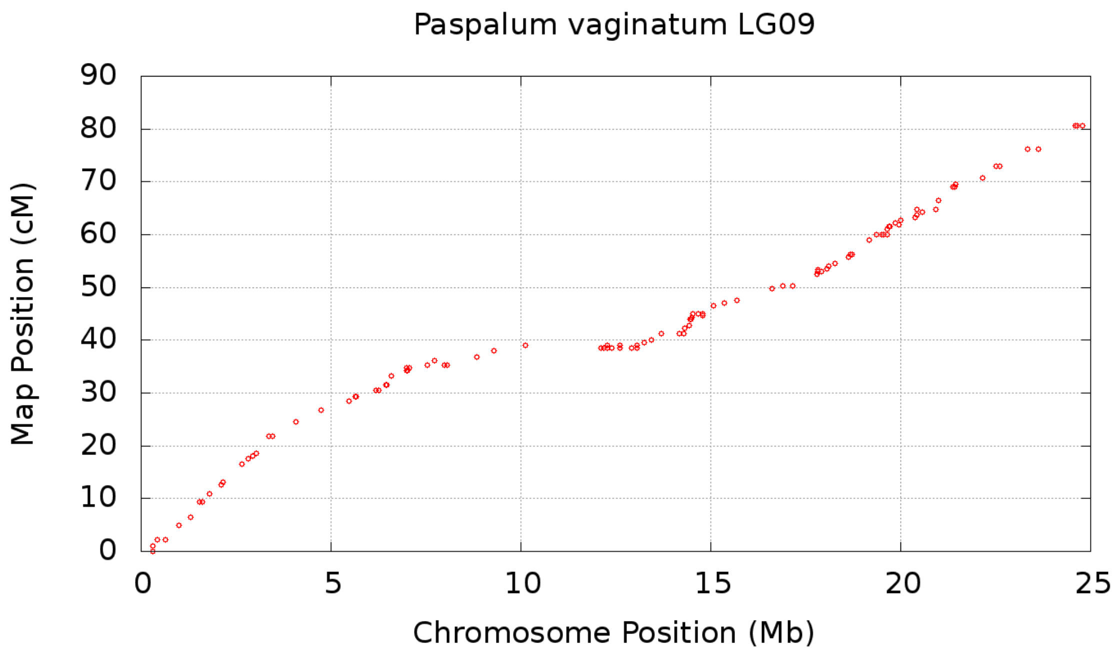


**Figure S3.9:** Marker map placements on the *Paspalum vaginatum* chromosome 9.

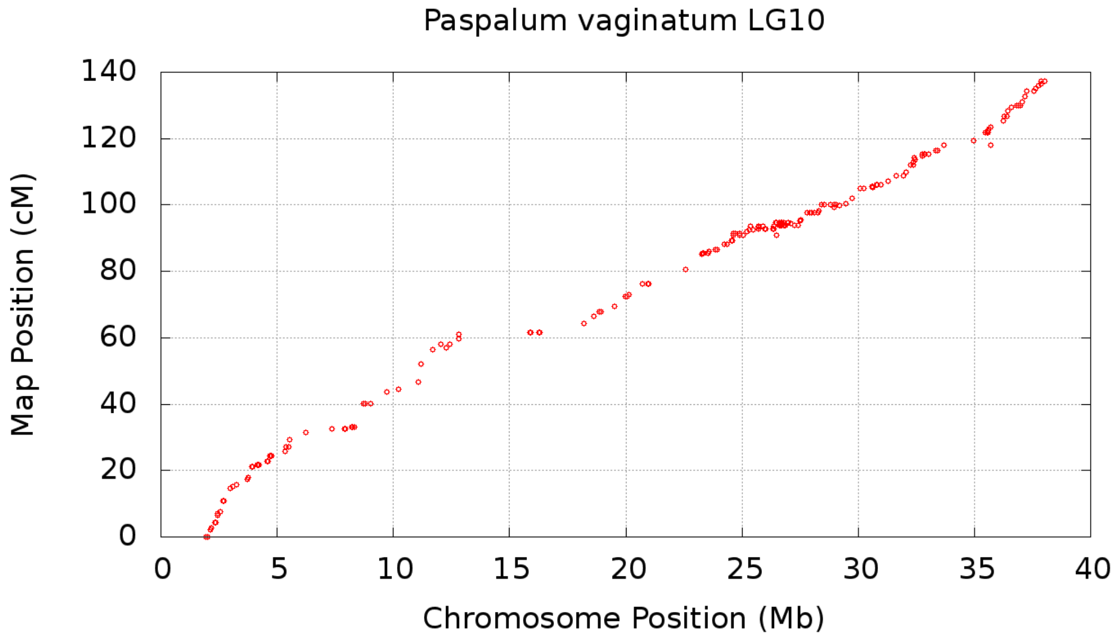


**Figure S3.10:** Marker map placements on the *Paspalum vaginatum* chromosome 10.


1. Chuan-Le Xiao, Ying Chen, Shang-Qian Xie, Kai-Ning Chen, Yan Wang, Yue Han, Feng Luo, Zhi Xie. MECAT: fast mapping, error correction, and de novo assembly for single-molecule sequencing reads. Nature Methods, 2017, 14: 1072-1074
2. Kent, W. J. BLAT--the BLAST-like alignment tool. Genome Res 12, 656-64 (2002).
3. Krumsiek J, Arnold R, Rattei T. Gepard: A rapid and sensitive tool for creating dotplots on genome scale. Bioinformatics 2007; 23(8): 1026-8. PMID: 17309896
4. Chin, C.-S. et al. Nonhybrid, finished microbial genome assemblies from long-read SMRT sequencing data. Nature Methods 10, 563–569 (2013)
5. Li H. (2013) Aligning sequence reads, clone sequences and assembly contigs with BWA-MEM. arXiv:1303.3997v1 [q-bio.GN]
6. McKenna A, Hanna M, Banks E, Sivachenko A, Cibulskis K, Kernytsky A, Garimella K, Altshuler D, Gabriel S, Daly M, DePristo MA: The Genome Analysis Toolkit: a MapReduce framework for analyzing next-generation DNA sequencing data. *Genome Res* 2010, 20:1297-1303.
7. Chin, C. S., Peluso, P., Sedlazeck, F. J., Nattestad, M., Concepcion, G. T., Clum, A., Dunn, C., O'Malley, R., Figueroa-Balderas, R., Morales-Cruz, A., Cramer, G. R., Delledonne, M., Luo, C., Ecker, J. R., Cantu, D., Rank, D. R., & Schatz, M. C. (2016). Phased diploid genome assembly with single-molecule real-time sequencing. Nature methods, 13(12), 1050–1054. https://doi.org/10.1038/nmeth.4035
