## Supplementary note 5 for "The genome of stress tolerant crop wild relative *Paspalum vaginatum* leads to increased biomass productivity in the crop *Zea mays*"

Fold_Change_Of_Primary_Metabolites

| Maize_Nitrogen_Deficit | | | | |
|---|---|---|---|---|
| | FoldChange | log2(FC)_Maize | p.value | -log10(p_value) |
| 1,3-diaminopropane | 0.362555808 | -1.4637250094 | 7.7E-06 | 5.11305715748 |
| 1,6-anhydro-glucose | 1.017820532 | 0.025483199944 | 0.70924 | 0.14920811902 |
| 2-hydroxypyridine | 1.133174789 | 0.180370410287 | 0.34823 | 0.4581337865 |
| 3-hydroxypyridine | 0.922807092 | -0.11589900306 | 0.30752 | 0.51212916514 |
| 3,4 dihydroxyphenylalanine (DOPA) | 1.477596857 | 0.563252702606 | 0.03782 | 1.42233053322 |
| 3,4-dihydroxybenzoic acid | 0.824137132 | -0.27904368152 | 0.20632 | 0.68545561688 |
| 3,5-dimethoxy-4-hydroxycinnamic acid | 2.345913661 | 1.230149917496 | 4.3E-07 | 6.37122145113 |
| 4-hydroxy-3-methoxybenzoic acid | 0.94797134 | -0.07708465242 | 0.59511 | 0.22540240665 |
| 4-hydroxybenzoic acid | 1.270765191 | 0.345697477034 | 0.04374 | 1.35907930525 |
| 4-hydroxycinnamic acid | 0.920010627 | -0.12027756877 | 0.38159 | 0.41840837096 |
| 5,6-dihydrouracil | 0.513446594 | -0.96171387342 | 0.0046 | 2.33695696051 |
| acetohydroxamic acid | 1.56076489 | 0.642253229474 | 0.02136 | 1.67044407149 |
| adenine | 0.602594916 | -0.73073959339 | 0.0152 | 1.81803023663 |
| adenosine | 0.450588544 | -1.15011746189 | 0.00152 | 2.81805205908 |
| allantoin | 0.03315122 | -4.91479422661 | 2.1E-09 | 8.68299487475 |
| alpha ketoglutaric acid | 1.091126252 | 0.125818042552 | 0.5186 | 0.28516450495 |
| Antiarol | 0.425472529 | -1.23286210913 | 0.00176 | 2.75375156576 |
| arachidic acid | 1.093471997 | 0.128916274133 | 0.4372 | 0.35931516494 |
| arbutin | 1.069880711 | 0.0974499494 | 0.86846 | 0.06125104321 |
| aspartic acid | 0.234530629 | -2.0921517494 | 2.2E-06 | 5.6567910245 |
| benzene-1,2,4-triol | 1.01347462 | 0.019309960512 | 0.88892 | 0.05113695273 |
| benzoic acid | 1.195436527 | 0.257537530746 | 0.18903 | 0.72346394695 |
| benzoquinone | 1.755488059 | 0.8118721829 | 0.00235 | 2.62954420071 |
| Beta- alanine | 0.063186257 | -3.98424538779 | 0.00013 | 3.90261340296 |
| beta-cyano-L-alanine | 0.238963978 | -2.06513493556 | 0.00171 | 2.76796826519 |
| beta-gentiobiose | 1.390670409 | 0.475780539002 | 0.02095 | 1.67891485324 |
| caffeic acid | 4.845156245 | 2.27654319006 | 0.00427 | 2.36955100052 |
| capric acid | 0.223433627 | -2.16208176747 | 0.00457 | 2.33994961182 |
| citric acid | 0.496910844 | -1.00894106885 | 2.9E-06 | 5.53531069941 |
| D-glucose | 1.012697007 | 0.018202593069 | 0.942 | 0.02595098321 |
| D-glucose-6-phosphate | 0.768272818 | -0.38030938415 | 0.14374 | 0.84242659069 |
| D-malic acid | 0.546113867 | -0.87272630546 | 2.8E-05 | 4.54758552313 |
| D-mannitol | 1.104531849 | 0.143435019304 | 0.56678 | 0.24658457504 |
| D-mannose | 1.01275246 | 0.018281588931 | 0.94172 | 0.0260800023 |
| D-saccharic acid | 1.549159689 | 0.631485866766 | 0.06266 | 1.20303517655 |
| dehydroascorbic acid | 0.558716235 | -0.8398123523 | 0.00122 | 2.91216603794 |
| eicosapentaenoic acid | 0.994972022 | -0.0072721358 | 0.98849 | 0.00502906903 |
| elaidic acid | 2.09480215 | 1.066813990311 | 1.2E-05 | 4.93444882762 |
| ethanolamine | 0.832170989 | -0.26504810125 | 0.52484 | 0.2799763355 |
| ferulic acid | 1.427592264 | 0.513583988416 | 0.00236 | 2.62631022464 |
| fructose | 0.786788556 | -0.34595212055 | 0.01124 | 1.94906005493 |
| fumaric acid | 0.538612177 | -0.89268124712 | 2.7E-05 | 4.57033535873 |
| galactinol | 0.539244026 | -0.89098980557 | 0.00014 | 3.85049308974 |
| gamma-aminobutyric acid (GABA) | 0.200400636 | -2.31904100461 | 4.8E-06 | 5.32077688608 |
| glyceric acid | 0.634254433 | -0.65686639692 | 0.00748 | 2.12638400616 |
| glycerol | 1.705708754 | 0.77037133033 | 0.24647 | 0.60824113987 |
| glycerol 1-phosphate | 2.697213766 | 1.431469865835 | 3.1E-05 | 4.50801463406 |

| | | | | |
|---|---|---|---|---|
| glycine | 0.832800694 | -0.26395682339 | 0.52674 | 0.278407606 |
| glyoxylic acid | 0.863775025 | -0.21127249089 | 0.10662 | 0.97217826764 |
| heptanoic acid | 1.743508388 | 0.801993305271 | 0.01068 | 1.97134926859 |
| itaconic acid | 1.028615619 | 0.04070396606 | 0.82019 | 0.08608788287 |
| L-alanine | 0.978679101 | -0.03109220327 | 0.82139 | 0.08545155119 |
| L-asparagine | 0.191050351 | -2.38797518765 | 0.00037 | 3.42959704733 |
| L-glutamic acid | 0.443393052 | -1.17334193055 | 4.7E-06 | 5.32569842336 |
| L-glutamine | 0.040450077 | -4.62771375472 | 0.00204 | 2.69043612969 |
| L-homoserine | 0.377737717 | -1.40454325164 | 4.9E-07 | 6.31135241427 |
| L-Isoleucine | 0.425370574 | -1.23320786011 | 0.03611 | 1.44237192011 |
| L-leucine | 0.425464611 | -1.23288895819 | 0.03622 | 1.44109630894 |
| L-lysine | 0.914010365 | -0.12971756926 | 0.61764 | 0.20926149896 |
| L-mimosine | 1.089499732 | 0.123665841522 | 0.72919 | 0.13715883185 |
| L-proline | 0.241955075 | -2.04718889378 | 0.14313 | 0.84427669424 |
| L-serine | 0.296545531 | -1.75367446233 | 0.00192 | 2.71635323704 |
| L-threonine | 0.217572147 | -2.20043421867 | 0.00046 | 3.33523271246 |
| L-tryptophan | 0.926059183 | -0.11082369792 | 0.62668 | 0.20295517052 |
| L-valine | 0.263301454 | -1.92521260637 | 7.5E-06 | 5.12307204776 |
| lactose | 0.750180827 | -0.41468970394 | 0.31008 | 0.5085280283 |
| lauric acid | 1.653726957 | 0.725721053828 | 0.23541 | 0.6281742703 |
| maleic acid | 2.428874419 | 1.280287899616 | 0.00046 | 3.33310783121 |
| maltose | 1.040432607 | 0.057183518974 | 0.66801 | 0.17521396204 |
| myo-inositol | 0.641307437 | -0.64091195783 | 0.00351 | 2.45513126672 |
| myristic acid | 0.347049329 | -1.52678735546 | 1.9E-07 | 6.72672412711 |
| nicotinic acid | 0.998669948 | -0.00192013649 | 0.99586 | 0.00180140173 |
| oxalic acid | 0.978769664 | -0.03095870739 | 0.82281 | 0.08469942267 |
| palmitic acid | 1.024713948 | 0.035221232871 | 0.61528 | 0.21092845196 |
| pantothenic acid | 0.51898557 | -0.94623366875 | 0.00459 | 2.33796871548 |
| pelargonic acid (nonanoic acid) | 0.91464399 | -0.12871778731 | 0.53907 | 0.26835358967 |
| phosphoric acid | 1.068163068 | 0.095131908762 | 0.66261 | 0.17874193918 |
| Pyruvic acid | 1.293039287 | 0.370766109322 | 0.01991 | 1.7008217047 |
| quinic acid | 1.416156721 | 0.501980932608 | 0.00017 | 3.77858913332 |
| ribose | 1.03406306 | 0.048324168391 | 0.39985 | 0.39810724888 |
| sedoheptulose anhydride monohydrate | 0.739922417 | -0.43455408744 | 0.00722 | 2.14144537488 |
| serotonin | 0.513629516 | -0.96119998437 | 0.25507 | 0.59334283263 |
| shikimic acid | 1.647202416 | 0.720017851108 | 5.2E-05 | 4.28381753225 |
| spermidine | 0.968681162 | -0.04590620887 | 0.60938 | 0.2151114346 |
| stearic acid | 1.336677061 | 0.418650954931 | 0.28263 | 0.54878108559 |
| succinic acid | 2.418504626 | 1.274115297158 | 0.00045 | 3.34468528802 |
| Sucrose | 1.055125401 | 0.077414473177 | 0.43257 | 0.36394118536 |
| threonic acid | 0.225166429 | -2.15093634851 | 1.3E-06 | 5.89922510434 |
| trans-aconitic acid | 0.975659965 | -0.03554966527 | 0.72528 | 0.13949517209 |
| Trehalose | 1.396150304 | 0.481454264952 | 0.01946 | 1.71078637494 |
| tyrosine | 0.723436401 | -0.46706190282 | 0.01613 | 1.79232637544 |
| xylose | 0.548901533 | -0.86538072567 | 0.04336 | 1.36291723819 |
| **Maize_Phosporus_Deficit** | | | | |
| | FoldChange | log2(FC) | p.value | -log10 |
| serotonin | 0.107905591 | -3.21215848256 | 0.04895 | 1.31022796269 |
| L-glutamine | 0.115678559 | -3.11180661444 | 0.00339 | 2.46992631324 |

Fold_Change_Of_Primary_Metabolites

| | | | | |
|---|---|---|---|---|
| capric acid | 0.217308676 | -2.20218231791 | 0.00431 | 2.36550029617 |
| D-glucose-6-phosphate | 0.23120021 | -2.1127853884 | 0.0002 | 3.70452402775 |
| Beta- alanine | 0.234217415 | -2.09407974321 | 0.0005 | 3.30132582984 |
| L-asparagine | 0.327815906 | -1.60904223869 | 0.00136 | 2.86702530338 |
| beta-cyano-L-alanine | 0.344945446 | -1.53555987958 | 0.00358 | 2.44563088874 |
| gamma-aminobutyric acid (GABA) | 0.35684399 | -1.48663461836 | 3.7E-07 | 6.4362395566 |
| adenosine | 0.361968137 | -1.46606538954 | 0.00043 | 3.36980534453 |
| myristic acid | 0.435466285 | -1.19936706815 | 5.6E-08 | 7.24917644893 |
| Antiarol | 0.442591119 | -1.17595359158 | 0.00015 | 3.82263269991 |
| glycerol 1-phosphate | 2.097624204 | 1.068756237814 | 5.1E-07 | 6.28910073055 |
| 1,3-diaminopropane | 0.508766255 | -0.97492511043 | 5.1E-05 | 4.29425573808 |
| 5,6-dihydrouracil | 0.543702196 | -0.87911144001 | 0.00468 | 2.32995020199 |
| L-threonine | 0.556223777 | -0.84626267883 | 0.00242 | 2.61691510604 |
| glyceric acid | 0.584918164 | -0.77369330472 | 0.00067 | 3.17212978953 |
| maleic acid | 0.588023116 | -0.7660552239 | 0.00036 | 3.44373531521 |
| succinic acid | 0.588459853 | -0.7649841026 | 0.00034 | 3.46902007578 |
| L-proline | 0.605332697 | -0.72419981497 | 0.45172 | 0.34513208243 |
| xylose | 0.653657573 | -0.61339303631 | 0.05117 | 1.29100935714 |
| phosphoric acid | 0.654603587 | -0.61130658686 | 1.7E-05 | 4.77871406362 |
| aspartic acid | 0.65724665 | -0.60549321211 | 1.1E-05 | 4.950611526 |
| alpha ketoglutaric acid | 0.65921818 | -0.60117206507 | 0.00645 | 2.19044197827 |
| L-serine | 0.667892535 | -0.58231210463 | 0.01687 | 1.77297128448 |
| allantoin | 0.669889432 | -0.57800510312 | 0.00101 | 2.99759025784 |
| shikimic acid | 1.487809472 | 0.57318978749 | 2.9E-06 | 5.54376261289 |
| L-glutamic acid | 0.672564861 | -0.57225469008 | 1.1E-06 | 5.96550968178 |
| galactinol | 0.672975643 | -0.57137380375 | 6.7E-05 | 4.17362132898 |
| citric acid | 0.681646504 | -0.55290433167 | 1.5E-06 | 5.81509368277 |
| threonic acid | 0.693909444 | -0.52718069253 | 0.0011 | 2.96023254447 |
| L-lysine | 0.704801293 | -0.50471152424 | 0.0757 | 1.12092137212 |
| L-valine | 0.706872679 | -0.50047771265 | 0.00517 | 2.28661743224 |
| stearic acid | 0.731019283 | -0.45201863274 | 0.03033 | 1.51815619223 |
| L-homoserine | 0.746673248 | -0.42145105292 | 0.00186 | 2.72982214235 |
| nicotinic acid | 0.74730043 | -0.42023974222 | 0.12708 | 0.89590947805 |
| quinic acid | 1.325845488 | 0.406912655344 | 1.3E-06 | 5.87952872954 |
| 3,4 dihydroxyphenylalanine (DOPA) | 1.309627684 | 0.389156723875 | 0.12624 | 0.89881325428 |
| pantothenic acid | 0.772824986 | -0.37178635668 | 0.07982 | 1.09786327489 |
| fumaric acid | 0.776965764 | -0.36407706472 | 0.00324 | 2.48985761807 |
| lactose | 0.780142012 | -0.35819132888 | 0.35112 | 0.4545435846 |
| pelargonic acid (nonanoic acid) | 1.277989941 | 0.353876480731 | 0.00453 | 2.34412616382 |
| maltose | 0.784631527 | -0.34991279017 | 0.01371 | 1.86307224996 |
| benzoquinone | 1.228554816 | 0.296962230063 | 0.05931 | 1.22685043145 |
| acetohydroxamic acid | 0.814510135 | -0.29599544515 | 0.49697 | 0.30366892358 |
| caffeic acid | 1.212695738 | 0.278217627292 | 0.56679 | 0.24657886154 |
| adenine | 1.211488311 | 0.276780485144 | 0.64136 | 0.19290143677 |
| dehydroascorbic acid | 0.829169401 | -0.27026121686 | 0.02134 | 1.67072470225 |
| beta-gentiobiose | 1.204346087 | 0.268250032182 | 0.1216 | 0.91505114841 |
| Trehalose | 1.204343769 | 0.268247255605 | 0.1216 | 0.91505122915 |
| D-malic acid | 0.838764053 | -0.25366306255 | 0.00248 | 2.60556308478 |
| glycerol | 0.848989105 | -0.23618205529 | 0.01178 | 1.92877182155 |

| | | | | |
|---|---|---|---|---|
| heptanoic acid | 0.857416108 | -0.22193257428 | 0.67145 | 0.1729847461 |
| ethanolamine | 0.861860993 | -0.21447289534 | 0.00741 | 2.13044251536 |
| glycine | 0.862382858 | -0.21359959439 | 0.00759 | 2.11987719067 |
| D-mannitol | 1.15732717 | 0.210796763787 | 0.3963 | 0.401973514 |
| 4-hydroxy-3-methoxybenzoic acid | 1.134854474 | 0.182507307424 | 0.11757 | 0.92969344645 |
| myo-inositol | 0.882913666 | -0.17965572096 | 0.06262 | 1.20332010192 |
| L-mimosine | 1.131274387 | 0.177948893146 | 0.62372 | 0.20501222729 |
| trans-aconitic acid | 1.12575856 | 0.17089744749 | 0.01304 | 1.88467060273 |
| tyrosine | 0.894356106 | -0.16107871007 | 0.14127 | 0.84993694585 |
| 3,5-dimethoxy-4-hydroxycinnamic acid | 0.896919949 | -0.15694886562 | 0.1417 | 0.84863731342 |
| fructose | 0.910592459 | -0.13512258247 | 0.10102 | 0.9955896074 |
| 4-hydroxycinnamic acid | 1.097310364 | 0.133971635716 | 0.33427 | 0.47590193698 |
| glyoxylic acid | 0.921334658 | -0.11820281062 | 0.30301 | 0.51854884731 |
| oxalic acid | 1.082866203 | 0.114854997131 | 0.29767 | 0.52626815826 |
| L-alanine | 1.080399751 | 0.111565212906 | 0.31285 | 0.50466012192 |
| L-tryptophan | 1.079870247 | 0.110857974764 | 0.48729 | 0.31221483402 |
| Sucrose | 0.927177292 | -0.10908286118 | 0.00717 | 2.1446708702 |
| Pyruvic acid | 1.072030864 | 0.100346441774 | 0.21968 | 0.65821333621 |
| 2-hydroxypyridine | 1.066111356 | 0.09235813621 | 0.17663 | 0.75294534471 |
| D-glucose | 1.064916137 | 0.090739821299 | 0.71349 | 0.14660984297 |
| ferulic acid | 0.939104245 | -0.09064278202 | 0.45119 | 0.34564073919 |
| arachidic acid | 0.940373095 | -0.0886948323 | 0.03689 | 1.4331080593 |
| D-mannose | 1.0616954 | 0.086369917017 | 0.72644 | 0.13879905443 |
| sedoheptulose anhydride monohydrate | 0.944143189 | -0.08292241935 | 0.22572 | 0.64642363594 |
| arbutin | 0.946134853 | -0.0798822691 | 0.89775 | 0.04684563399 |
| 3,4-dihydroxybenzoic acid | 0.956748564 | -0.06378826385 | 0.61379 | 0.2119770747 |
| spermidine | 0.956887277 | -0.06357911215 | 0.04762 | 1.32224493071 |
| benzene-1,2,4-triol | 1.037320271 | 0.052861392958 | 0.22771 | 0.6426124006 |
| itaconic acid | 0.964670738 | -0.05189149017 | 0.74644 | 0.12700667491 |
| 3-hydroxypyridine | 0.966519588 | -0.04912912426 | 0.52378 | 0.28085257374 |
| lauric acid | 1.02764904 | 0.039347642897 | 0.60984 | 0.21478457815 |
| 4-hydroxybenzoic acid | 0.974821223 | -0.03679043496 | 0.38298 | 0.41682724135 |
| benzoic acid | 0.976546671 | -0.03423909996 | 0.62694 | 0.20277475671 |
| elaidic acid | 1.021438653 | 0.030602558934 | 0.8005 | 0.09663793644 |
| palmitic acid | 0.984830547 | -0.02205258355 | 0.23038 | 0.63756074228 |
| 1,6-anhydro-glucose | 1.012283139 | 0.017612872851 | 0.55842 | 0.25303854805 |
| ribose | 1.010739047 | 0.015410569872 | 0.74703 | 0.12666198667 |
| eicosapentaenoic acid | 1.004672508 | 0.006725304524 | 0.99192 | 0.00352169388 |
| L-Isoleucine | 1.003074068 | 0.004428139794 | 0.9894 | 0.00462615157 |
| L-leucine | 1.00298118 | 0.004294534727 | 0.98973 | 0.00448541878 |
| D-saccharic acid | 0.999429939 | -0.00082265857 | 0.99597 | 0.00175213407 |
| **Sorghum_Nitrogen_Deficit** | | | | |
| | FoldChange | log2(FC) | p.value | -log10 |
| L-asparagine | 0.126201107 | -2.9862035304 | 1.3E-05 | 4.88912514424 |
| beta-cyano-L-alanine | 0.126920327 | -2.97800494853 | 5.2E-05 | 4.284581685 |
| Beta- alanine | 0.133614777 | -2.90384852289 | 3E-05 | 4.51979497087 |
| L-glutamine | 0.1526831 | -2.71138770767 | 6.2E-05 | 4.21064546182 |
| 5,6-dihydrouracil | 0.166922435 | -2.58275022583 | 6.5E-06 | 5.18952591457 |
| allantoin | 0.213289914 | -2.22911235174 | 2.3E-06 | 5.64536654125 |

| | | | | |
|---|---|---|---|---|
| stearic acid | 0.238993164 | -2.06495874459 | 0.18152 | 0.74108003638 |
| L-threonine | 0.270068976 | -1.88860017481 | 3.8E-06 | 5.41866195921 |
| caffeic acid | 3.170510361 | 1.664715091589 | 0.00866 | 2.06237173888 |
| shikimic acid | 2.991944947 | 1.581083629153 | 0.00454 | 2.34276123832 |
| L-valine | 0.345554265 | -1.53301580855 | 1.8E-06 | 5.73946895706 |
| gamma-aminobutyric acid (GABA) | 0.349517966 | -1.51656147896 | 1.4E-07 | 6.85021582614 |
| 3,5-dimethoxy-4-hydroxycinnamic acid | 2.811605678 | 1.491394273528 | 0.00091 | 3.04292434231 |
| capric acid | 0.364595189 | -1.45563257225 | 1.5E-05 | 4.82390668391 |
| L-serine | 0.366929126 | -1.44642666608 | 2.7E-05 | 4.56706788091 |
| 1,3-diaminopropane | 0.382483889 | -1.386529115 | 0.00014 | 3.84710365578 |
| L-proline | 0.399763724 | -1.32278053317 | 0.00723 | 2.1410408068 |
| itaconic acid | 0.41746068 | -1.26028777639 | 0.00433 | 2.36378523842 |
| aspartic acid | 0.435339345 | -1.19978768008 | 5E-06 | 5.29915364393 |
| maleic acid | 2.214571593 | 1.147027637511 | 9.1E-05 | 4.0391648517 |
| succinic acid | 2.200078289 | 1.137554862225 | 9.2E-05 | 4.03792912469 |
| Antiarol | 0.464545694 | -1.10610758509 | 0.00192 | 2.7161322289 |
| spermidine | 0.488621805 | -1.03320984805 | 0.00174 | 2.76058705825 |
| threonic acid | 0.494842464 | -1.014958788 | 0.00023 | 3.63389372807 |
| galactinol | 1.973533736 | 0.980781181066 | 0.1204 | 0.91936424256 |
| elaidic acid | 1.971475842 | 0.979276032465 | 0.00343 | 2.46449719595 |
| L-glutamic acid | 0.514442583 | -0.95891802925 | 2.9E-05 | 4.53092811556 |
| myristic acid | 0.517659624 | -0.94992429857 | 0.00319 | 2.49630056487 |
| myo-inositol | 1.917291545 | 0.939069731291 | 0.02331 | 1.63251293149 |
| citric acid | 0.555921381 | -0.84704722465 | 0.00075 | 3.12281223475 |
| adenosine | 0.563171781 | -0.82835304848 | 0.00115 | 2.93987519524 |
| quinic acid | 1.724097937 | 0.785841728566 | 0.00476 | 2.3220161061 |
| 3,4 dihydroxyphenylalanine (DOPA) | 1.713364019 | 0.776831697129 | 0.07758 | 1.11022449369 |
| dehydroascorbic acid | 0.602272773 | -0.73151105541 | 0.00241 | 2.61734639129 |
| L-homoserine | 0.653847632 | -0.61297361436 | 0.01462 | 1.83515454466 |
| serotonin | 0.656669957 | -0.6067596422 | 0.0354 | 1.4510445118 |
| 3,4-dihydroxybenzoic acid | 1.495861982 | 0.580977068432 | 0.17038 | 0.76857543426 |
| alpha ketoglutaric acid | 1.47324735 | 0.558999670818 | 0.02898 | 1.53792090229 |
| L-alanine | 1.408194869 | 0.49384699066 | 0.00059 | 3.23136734083 |
| oxalic acid | 1.404146862 | 0.489693837047 | 0.00064 | 3.19245662575 |
| arachidic acid | 0.722904094 | -0.46812383371 | 0.00308 | 2.5110085084 |
| L-leucine | 0.731708137 | -0.45065979145 | 0.10557 | 0.97644932456 |
| L-Isoleucine | 0.732025946 | -0.45003331103 | 0.10574 | 0.97577814746 |
| L-tryptophan | 1.348043772 | 0.430867342622 | 0.01468 | 1.83326242428 |
| D-malic acid | 0.743730569 | -0.42714802428 | 0.01445 | 1.84024733191 |
| glyceric acid | 0.747656668 | -0.41955217459 | 0.00942 | 2.02602305136 |
| benzoquinone | 1.33708607 | 0.419092336497 | 0.01968 | 1.70603156379 |
| eicosapentaenoic acid | 0.752600362 | -0.41004411074 | 0.12346 | 0.90846639067 |
| 4-hydroxybenzoic acid | 1.291773848 | 0.369353517807 | 0.25119 | 0.59999051835 |
| D-glucose-6-phosphate | 0.778462616 | -0.36130033632 | 0.10537 | 0.9772666087 |
| adenine | 0.786844009 | -0.34585044289 | 0.42525 | 0.37135274605 |
| fumaric acid | 0.788459597 | -0.34289126636 | 0.07793 | 1.10829758528 |
| glycine | 0.825639889 | -0.27641542203 | 0.03189 | 1.4963176565 |
| ethanolamine | 0.827200568 | -0.27369091822 | 0.03291 | 1.48264247677 |
| L-lysine | 1.206721294 | 0.271092508156 | 0.37628 | 0.42448614329 |

| | | | | |
|---|---|---|---|---|
| glycerol | 1.179488347 | 0.238161165271 | 0.28858 | 0.53974062965 |
| 4-hydroxy-3-methoxybenzoic acid | 1.166177887 | 0.221787871592 | 0.3109 | 0.50737724218 |
| L-mimosine | 0.867059828 | -0.20579655012 | 0.4395 | 0.35703642561 |
| D-saccharic acid | 0.872139916 | -0.19736849223 | 0.51646 | 0.28696058421 |
| Sucrose | 1.142903424 | 0.192703499863 | 0.05385 | 1.26879671636 |
| heptanoic acid | 1.135237469 | 0.182994112271 | 0.39097 | 0.40785714715 |
| lactose | 1.134086049 | 0.18153010949 | 0.09357 | 1.02887798266 |
| 4-hydroxycinnamic acid | 0.887045366 | -0.17292020471 | 0.33093 | 0.48026937638 |
| lauric acid | 0.889529613 | -0.16888546061 | 0.15152 | 0.81952739435 |
| benzoic acid | 1.116565688 | 0.159068127481 | 0.06821 | 1.16612891527 |
| xylose | 1.112992876 | 0.154444357754 | 0.43169 | 0.36483171834 |
| pantothenic acid | 0.898666439 | -0.15414236967 | 0.53847 | 0.26883535051 |
| Trehalose | 1.11255469 | 0.153876256826 | 0.49058 | 0.3092905524 |
| pelargonic acid (nonanoic acid) | 0.901533323 | -0.14954727647 | 0.18675 | 0.72874138574 |
| glycerol 1-phosphate | 0.9021365 | -0.14858235482 | 0.40969 | 0.38754176452 |
| beta-gentiobiose | 1.103945617 | 0.142669103501 | 0.52871 | 0.27678192308 |
| ferulic acid | 1.093526273 | 0.128987883154 | 0.17152 | 0.76567514415 |
| trans-aconitic acid | 1.091167361 | 0.125872395748 | 0.41127 | 0.3858694738 |
| phosphoric acid | 1.084176371 | 0.116599469382 | 0.38953 | 0.40946156652 |
| Pyruvic acid | 1.08008614 | 0.111146376064 | 0.34935 | 0.45673871015 |
| 2-hydroxypyridine | 0.933244825 | -0.09967249177 | 0.57054 | 0.24371390243 |
| fructose | 0.933902039 | -0.09865686816 | 0.3718 | 0.42969260256 |
| acetohydroxamic acid | 1.06522559 | 0.091158991501 | 0.5005 | 0.30059336936 |
| arbutin | 0.96417429 | -0.05263413424 | 0.9183 | 0.03701712495 |
| palmitic acid | 1.032674833 | 0.046386052177 | 0.38853 | 0.41057792552 |
| D-mannose | 0.971489842 | -0.04172918394 | 0.65697 | 0.18245463389 |
| D-glucose | 0.971681913 | -0.04144397991 | 0.65893 | 0.1811600719 |
| D-mannitol | 0.972394885 | -0.04038579066 | 0.66812 | 0.17514426167 |
| glyoxylic acid | 1.026250461 | 0.037382869776 | 0.85898 | 0.06601458167 |
| benzene-1,2,4-triol | 0.978056187 | -0.03201074855 | 0.55575 | 0.25512269463 |
| 1,6-anhydro-glucose | 1.022058191 | 0.031477338536 | 0.53049 | 0.27532110278 |
| tyrosine | 0.981421149 | -0.02705573414 | 0.83488 | 0.0783738842 |
| sedoheptulose anhydride monohydrate | 0.983322519 | -0.02426341293 | 0.83101 | 0.0803916263 |
| ribose | 1.012828328 | 0.018389661324 | 0.80423 | 0.09462112983 |
| nicotinic acid | 1.009407948 | 0.013509351539 | 0.93953 | 0.02708794394 |
| 3-hydroxypyridine | 0.996891591 | -0.00449147052 | 0.96562 | 0.01519512183 |
| maltose | 1.00055257 | 0.000796969255 | 0.99595 | 0.00176263491 |
| **Sorghum_Phosphorus_Deficit** | | | | |
| | FoldChange | log2(FC) | p.value | -log10 |
| D-glucose-6-phosphate | 0.208797622 | -2.25982281561 | 2.4E-05 | 4.62757970922 |
| 5,6-dihydrouracil | 0.226048306 | -2.1452969881 | 1.4E-05 | 4.86922452954 |
| beta-cyano-L-alanine | 0.227285532 | -2.13742224644 | 0.00014 | 3.86828795082 |
| L-asparagine | 0.240799979 | -2.05409282751 | 4E-05 | 4.40029326415 |
| serotonin | 4.103188135 | 2.036745304605 | 0.10951 | 0.96054720976 |
| stearic acid | 0.356893831 | -1.48643313259 | 0.27217 | 0.56516140511 |
| L-glutamine | 0.35724488 | -1.48501476196 | 0.00045 | 3.34231895394 |
| capric acid | 0.40173401 | -1.31568749113 | 3.9E-05 | 4.4136577417 |
| itaconic acid | 0.480391238 | -1.05771825594 | 0.01137 | 1.94405318288 |
| glycerol 1-phosphate | 0.481941325 | -1.05307058164 | 0.00022 | 3.65451691495 |

| | | | | |
|---|---|---|---|---|
| 1,3-diaminopropane | 0.486397017 | -1.0397937143 | 0.00091 | 3.04309232826 |
| L-threonine | 0.495273054 | -1.01370396235 | 0.00029 | 3.53346174093 |
| L-mimosine | 0.567373771 | -0.81762863553 | 0.02722 | 1.56513766793 |
| myristic acid | 0.568864936 | -0.8138419355 | 0.00546 | 2.26283623105 |
| succinic acid | 0.572905203 | -0.80363165607 | 0.02365 | 1.62621236473 |
| maleic acid | 0.573913871 | -0.80109385213 | 0.0249 | 1.60383463468 |
| Beta- alanine | 0.579883035 | -0.78616616455 | 0.0057 | 2.24445355492 |
| L-serine | 0.585324565 | -0.77269126886 | 0.00069 | 3.16318772413 |
| adenosine | 0.603455532 | -0.7286806318 | 0.00118 | 2.92751798281 |
| L-valine | 0.629548633 | -0.66761026405 | 0.00042 | 3.37724328316 |
| spermidine | 0.632621682 | -0.66058509162 | 0.02768 | 1.55788741953 |
| 3,5-dimethoxy-4-hydroxycinnamic acid | 1.570382791 | 0.651116268111 | 3.8E-06 | 5.4197794256 |
| citric acid | 0.640603918 | -0.64249547267 | 0.00222 | 2.65284423393 |
| adenine | 0.641862079 | -0.63966476574 | 0.19192 | 0.71688272126 |
| phosphoric acid | 0.649974331 | -0.62154535011 | 0.00432 | 2.36451957741 |
| caffeic acid | 1.472883549 | 0.558643370818 | 0.00549 | 2.26077013382 |
| glycine | 0.701359297 | -0.51177438686 | 0.00357 | 2.44757231257 |
| ethanolamine | 0.701959102 | -0.51054111742 | 0.00357 | 2.4470688043 |
| alpha ketoglutaric acid | 1.386659346 | 0.471613411105 | 0.01105 | 1.95650898747 |
| L-proline | 0.721727727 | -0.47047341461 | 0.18706 | 0.72801077922 |
| myo-inositol | 1.373639966 | 0.458003920134 | 0.00287 | 2.54203839697 |
| aspartic acid | 0.729842037 | -0.45434384659 | 0.0003 | 3.52820026044 |
| glyceric acid | 0.742015038 | -0.43047966992 | 0.00807 | 2.09309526259 |
| threonic acid | 0.74853931 | -0.41785001269 | 0.0122 | 1.91365082663 |
| L-leucine | 0.753074065 | -0.40913633341 | 0.13365 | 0.87402717787 |
| L-Isoleucine | 0.753146029 | -0.40899847661 | 0.13382 | 0.87347088335 |
| Antiarol | 0.758452734 | -0.39886881831 | 0.11137 | 0.9532409809 |
| Trehalose | 0.765521021 | -0.38548610078 | 3.1E-05 | 4.51174974229 |
| beta-gentiobiose | 0.77043593 | -0.37625310809 | 4.1E-05 | 4.39026863924 |
| D-saccharic acid | 0.792368972 | -0.33575570753 | 0.01407 | 1.85174085562 |
| 4-hydroxycinnamic acid | 1.255013514 | 0.327702899547 | 0.01443 | 1.84073925097 |
| glycerol | 0.797679523 | -0.32611885109 | 0.19384 | 0.7125457001 |
| L-glutamic acid | 0.808725107 | -0.30627869491 | 0.00494 | 2.30642522725 |
| arachidic acid | 0.8141868 | -0.29656826365 | 0.06626 | 1.17873280478 |
| benzoquinone | 1.223601983 | 0.291134350238 | 0.04852 | 1.31409201961 |
| maltose | 0.832519116 | -0.26444469618 | 0.00245 | 2.6111641015 |
| pantothenic acid | 0.834929541 | -0.26027364035 | 0.27031 | 0.56813269522 |
| pelargonic acid (nonanoic acid) | 0.840505124 | -0.25067147987 | 0.09 | 1.04576024488 |
| L-homoserine | 0.859342673 | -0.21869455708 | 0.30698 | 0.51289551248 |
| quinic acid | 0.880122361 | -0.18422398385 | 0.23533 | 0.62831439463 |
| trans-aconitic acid | 0.882095704 | -0.18099290334 | 0.12556 | 0.9011556534 |
| nicotinic acid | 0.883192032 | -0.17920093893 | 0.30901 | 0.51003208766 |
| elaidic acid | 1.129004927 | 0.175051782707 | 0.16049 | 0.79454980489 |
| arbutin | 0.891129487 | -0.16629301422 | 0.73701 | 0.13252446897 |
| galactinol | 1.119796904 | 0.163237096737 | 0.40175 | 0.39604396918 |
| lactose | 0.896956855 | -0.15688950324 | 0.5056 | 0.29618935193 |
| heptanoic acid | 0.898785348 | -0.15395148866 | 0.57126 | 0.24316809936 |
| 2-hydroxypyridine | 1.103429995 | 0.141995104078 | 0.32547 | 0.48749059728 |
| acetohydroxamic acid | 0.910226391 | -0.13570267936 | 0.49114 | 0.30879034748 |

| | | | | |
|---|---|---|---|---|
| shikimic acid | 0.91980144 | -0.12060563891 | 0.46216 | 0.33520674122 |
| allantoin | 0.925506647 | -0.11168474382 | 0.44584 | 0.35082160771 |
| 4-hydroxybenzoic acid | 0.945760364 | -0.08045341443 | 0.48266 | 0.31635431369 |
| eicosapentaenoic acid | 0.951117224 | -0.07230493237 | 0.76312 | 0.11740837762 |
| L-tryptophan | 0.956861477 | -0.06361801051 | 0.64684 | 0.18920560053 |
| sedoheptulose anhydride monohydrate | 0.957102992 | -0.0632539157 | 0.55191 | 0.25813538334 |
| D-mannitol | 0.958533558 | -0.06109915402 | 0.47253 | 0.32557468306 |
| D-mannose | 0.958685214 | -0.06087091286 | 0.47457 | 0.32370121601 |
| Sucrose | 1.042999872 | 0.06073898022 | 0.12144 | 0.91563289813 |
| D-glucose | 0.960876506 | -0.05757707066 | 0.50572 | 0.29608778351 |
| 3,4 dihydroxyphenylalanine (DOPA) | 1.039170144 | 0.05543188743 | 0.8376 | 0.07696473369 |
| fructose | 0.963452506 | -0.0537145456 | 0.54293 | 0.26525777991 |
| benzene-1,2,4-triol | 0.963470046 | -0.05368828054 | 0.40982 | 0.38740177391 |
| 3-hydroxypyridine | 0.963636854 | -0.05343852423 | 0.48184 | 0.31709270564 |
| L-alanine | 0.966735027 | -0.04880758051 | 0.56699 | 0.24642739248 |
| oxalic acid | 0.967590917 | -0.0475308686 | 0.58155 | 0.23541268298 |
| tyrosine | 0.97080884 | -0.04274084945 | 0.70828 | 0.1497953313 |
| 3,4-dihydroxybenzoic acid | 1.029112077 | 0.041400109724 | 0.39652 | 0.40173626827 |
| xylose | 0.972266233 | -0.04057667821 | 0.82891 | 0.08149226606 |
| lauric acid | 0.972297243 | -0.04053066393 | 0.81812 | 0.08718075897 |
| palmitic acid | 1.022189629 | 0.031662859438 | 0.3915 | 0.4072694331 |
| Pyruvic acid | 1.020728731 | 0.02959950579 | 0.65361 | 0.18468380124 |
| fumaric acid | 0.98088854 | -0.02783888445 | 0.85403 | 0.06852676466 |
| ferulic acid | 1.019216985 | 0.02746122567 | 0.6883 | 0.16222356298 |
| D-malic acid | 0.983335566 | -0.0242442706 | 0.84189 | 0.07474329145 |
| dehydroascorbic acid | 1.016627276 | 0.023790843622 | 0.83831 | 0.07659367052 |
| gamma-aminobutyric acid (GABA) | 0.983894156 | -0.02342497114 | 0.86093 | 0.06503158629 |
| 1,6-anhydro-glucose | 1.010624371 | 0.015246876227 | 0.74657 | 0.12692981588 |
| ribose | 0.99292681 | -0.01024071556 | 0.84628 | 0.07248362073 |
| 4-hydroxy-3-methoxybenzoic acid | 0.994912047 | -0.00735910237 | 0.9584 | 0.018454206 |
| L-lysine | 1.004916924 | 0.007076238776 | 0.97298 | 0.01189703844 |
| glyoxylic acid | 1.003119011 | 0.004492778472 | 0.9762 | 0.01046306885 |
| benzoic acid | 1.002155326 | 0.003106131891 | 0.96108 | 0.01723964775 |
| **Paspalum_Nitrogen_Deficit** | | | | |
| | FoldChange | log2(FC) | p.value | -log10 |
| xylose | 0.13344705 | -2.90566068081 | 0.04086 | 1.38865790558 |
| allantoin | 0.146596725 | -2.77007522363 | 0.00065 | 3.18610528686 |
| glyceric acid | 0.231880981 | -2.10854360107 | 0.02145 | 1.66854647036 |
| L-proline | 0.273859505 | -1.86849213884 | 0.11064 | 0.95610103353 |
| Trehalose | 3.215359593 | 1.684980092056 | 0.02045 | 1.68925647091 |
| 4-hydroxy-3-methoxybenzoic acid | 2.837981449 | 1.504865159026 | 0.20259 | 0.69337411598 |
| maleic acid | 2.694072759 | 1.429788814515 | 2E-05 | 4.69914369455 |
| succinic acid | 2.688252356 | 1.426668575282 | 1.9E-05 | 4.73011958169 |
| L-lysine | 2.535717677 | 1.342394126887 | 1.6E-08 | 7.7909943652 |
| 1,3-diaminopropane | 0.431139956 | -1.2137718252 | 0.00936 | 2.02869140329 |
| beta-gentiobiose | 2.006194519 | 1.004461495321 | 0.03059 | 1.51448144991 |
| glycerol 1-phosphate | 1.979111095 | 0.984852598573 | 0.00065 | 3.18580221401 |
| shikimic acid | 1.960724704 | 0.971386988182 | 2E-05 | 4.70182460074 |
| quinic acid | 1.940460436 | 0.956399018514 | 0.00012 | 3.90420634096 |

| | | | | |
|---|---|---|---|---|
| threonic acid | 0.518206112 | -0.94840206228 | 0.04496 | 1.3472012096 |
| D-glucose-6-phosphate | 1.839132157 | 0.879025153042 | 0.00636 | 2.19666132238 |
| gamma-aminobutyric acid (GABA) | 0.54426116 | -0.87762901067 | 0.06206 | 1.20716192957 |
| sedoheptulose anhydride monohydrate | 1.831407381 | 0.872952742313 | 0.00036 | 3.43850441779 |
| pantothenic acid | 1.719265202 | 0.781792102122 | 0.00152 | 2.81697002827 |
| alpha ketoglutaric acid | 1.699429465 | 0.765050483794 | 0.0013 | 2.88497449797 |
| aspartic acid | 0.599644991 | -0.73781946235 | 0.00236 | 2.62661482192 |
| galactinol | 1.626620219 | 0.701877451619 | 0.00515 | 2.28779617869 |
| Beta- alanine | 0.618662256 | -0.69277607493 | 0.0746 | 1.12727895637 |
| 4-hydroxybenzoic acid | 1.569470886 | 0.650278267284 | 0.028 | 1.55286110161 |
| heptanoic acid | 1.549933675 | 0.632206480678 | 0.02076 | 1.68267050621 |
| acetohydroxamic acid | 1.48151699 | 0.567075171641 | 0.02012 | 1.6964487062 |
| dehydroascorbic acid | 0.680238135 | -0.55588820828 | 0.04592 | 1.33798781021 |
| tyrosine | 1.419831148 | 0.505719369166 | 3.5E-05 | 4.45920054277 |
| L-valine | 0.71828154 | -0.47737865617 | 0.1678 | 0.77521969212 |
| L-homoserine | 0.732013704 | -0.4500574371 | 0.12411 | 0.9061795759 |
| D-mannose | 1.361460627 | 0.445155260517 | 0.00546 | 2.262516733 |
| D-glucose | 1.360767915 | 0.444421029451 | 0.00539 | 2.26817940423 |
| D-mannitol | 1.36006858 | 0.443679400017 | 0.00561 | 2.25067679503 |
| L-tryptophan | 0.743979587 | -0.4266650576 | 0.2158 | 0.66593892601 |
| nicotinic acid | 1.337450452 | 0.419485445673 | 0.04531 | 1.34383493838 |
| stearic acid | 0.762880299 | -0.3904713877 | 0.38849 | 0.4106147418 |
| D-malic acid | 0.776230252 | -0.36544343493 | 0.12112 | 0.91679952427 |
| eicosapentaenoic acid | 0.778417648 | -0.36138367631 | 0.44485 | 0.35178221851 |
| phosphoric acid | 1.282927949 | 0.359440148629 | 0.02854 | 1.54452240543 |
| L-mimosine | 1.273285473 | 0.348555909971 | 0.35147 | 0.45411088722 |
| 3,4-dihydroxybenzoic acid | 0.791993732 | -0.3364390828 | 0.14174 | 0.84852156291 |
| fumaric acid | 0.794657856 | -0.33159426081 | 0.17844 | 0.74850269654 |
| L-alanine | 1.251181813 | 0.32329144674 | 0.00094 | 3.02880505312 |
| oxalic acid | 1.249587555 | 0.321451990874 | 0.00082 | 3.08440698076 |
| benzoic acid | 1.245206331 | 0.31638481655 | 0.00188 | 2.72482330159 |
| arachidic acid | 1.241687375 | 0.31230198557 | 0.03135 | 1.50377935425 |
| L-glutamic acid | 0.809879026 | -0.30422167011 | 0.1587 | 0.79941298087 |
| 3-hydroxypyridine | 0.81412178 | -0.29668347896 | 0.00672 | 2.17267679268 |
| benzoquinone | 1.226802683 | 0.294903227377 | 0.20545 | 0.68730295587 |
| beta-cyano-L-alanine | 1.224805263 | 0.292552387313 | 0.35035 | 0.4554968313 |
| adenine | 1.20516558 | 0.269231375218 | 0.17113 | 0.76667211544 |
| 2-hydroxypyridine | 0.835135861 | -0.25991717878 | 0.07724 | 1.11215869744 |
| L-Isoleucine | 0.837902545 | -0.25514563935 | 0.56482 | 0.24809028204 |
| lauric acid | 1.191325259 | 0.252567355757 | 0.02264 | 1.64517465428 |
| L-leucine | 0.839481697 | -0.25242922372 | 0.56823 | 0.2454794453 |
| D-saccharic acid | 0.859042849 | -0.21919800019 | 0.26861 | 0.57087670851 |
| L-serine | 0.863343726 | -0.21199303619 | 0.65375 | 0.18458964235 |
| ethanolamine | 1.150627003 | 0.202420233005 | 0.48285 | 0.31618415474 |
| glycine | 1.149635263 | 0.201176220005 | 0.48606 | 0.31331212778 |
| benzene-1,2,4-triol | 1.148913211 | 0.200269820387 | 0.14055 | 0.85216215039 |
| L-threonine | 0.87342663 | -0.1952415761 | 0.67806 | 0.16873143372 |
| L-glutamine | 0.878122051 | -0.18750661906 | 0.48667 | 0.31276870507 |
| glycerol | 1.136688219 | 0.184836592799 | 0.39318 | 0.40540824824 |

| | | | | |
|---|---|---|---|---|
| 3,5-dimethoxy-4-hydroxycinnamic acid | 1.124825141 | 0.169700745751 | 0.06509 | 1.18646734457 |
| fructose | 1.105793022 | 0.145081372414 | 0.21763 | 0.66229038578 |
| ribose | 0.909318677 | -0.13714210884 | 0.07047 | 1.15202246479 |
| maltose | 1.093276549 | 0.128658383569 | 0.51522 | 0.28800394819 |
| 4-hydroxycinnamic acid | 0.917594673 | -0.12407107926 | 0.40615 | 0.39131369785 |
| ferulic acid | 1.089557611 | 0.123742481731 | 0.34073 | 0.46758654284 |
| 3,4 dihydroxyphenylalanine (DOPA) | 0.920063099 | -0.12019528852 | 0.23639 | 0.62637830138 |
| trans-aconitic acid | 0.92764593 | -0.10835384188 | 0.44904 | 0.34771388759 |
| itaconic acid | 0.927987879 | -0.10782213283 | 0.63986 | 0.19391298764 |
| capric acid | 1.074660731 | 0.103881274604 | 0.36054 | 0.44304984905 |
| L-asparagine | 1.073456146 | 0.10226325391 | 0.66654 | 0.17617557746 |
| adenosine | 0.932580437 | -0.10069992818 | 0.63409 | 0.19785014301 |
| lactose | 1.065518812 | 0.091556064946 | 0.82105 | 0.08563216214 |
| myo-inositol | 0.940728151 | -0.08815021739 | 0.69579 | 0.15752080982 |
| spermidine | 0.941520554 | -0.0869355048 | 0.12996 | 0.88618077338 |
| serotonin | 0.950541342 | -0.07317871871 | 0.72641 | 0.13881622397 |
| 5,6-dihydrouracil | 0.955402825 | -0.06581895191 | 0.31678 | 0.49923633409 |
| glyoxylic acid | 1.044836991 | 0.063277880148 | 0.65457 | 0.18404341066 |
| Sucrose | 1.042870945 | 0.060560636304 | 0.62046 | 0.20728590343 |
| caffeic acid | 0.965673386 | -0.05039277782 | 0.57982 | 0.23670489582 |
| 1,6-anhydro-glucose | 1.034676524 | 0.049179801751 | 0.05249 | 1.27990225754 |
| Pyruvic acid | 0.970121679 | -0.04376238344 | 0.52664 | 0.27848564148 |
| palmitic acid | 0.972692451 | -0.03994437349 | 0.43411 | 0.36240256907 |
| Antiarol | 0.979079972 | -0.03050139041 | 0.83078 | 0.08051564844 |
| citric acid | 0.982688089 | -0.02519452596 | 0.83212 | 0.07981423107 |
| elaidic acid | 1.006385057 | 0.009182406433 | 0.95639 | 0.01936624606 |
| arbutin | 1.006142127 | 0.008834113875 | 0.98301 | 0.00744323893 |
| pelargonic acid (nonanoic acid) | 0.998054289 | -0.0028098024 | 0.9845 | 0.00678564636 |
| myristic acid | 0.998422846 | -0.00227714907 | 0.98982 | 0.00444394043 |
| **Paspalum_Phosphorus_Deficit** | | | | |
| | FoldChange | log2(FC) | p.value | -log10 |
| L-proline | 0.168938443 | -2.56543043618 | 0.07768 | 1.10967472506 |
| xylose | 0.170220826 | -2.55452053697 | 0.04812 | 1.31769995682 |
| Trehalose | 5.584368583 | 2.481394166374 | 0.00864 | 2.06335851657 |
| glyceric acid | 0.215402267 | -2.21489466066 | 0.01951 | 1.70980736183 |
| L-tryptophan | 0.391370261 | -1.35339396207 | 0.01452 | 1.83808155138 |
| galactinol | 1.773102767 | 0.826276155772 | 0.01358 | 1.86708944734 |
| 4-hydroxybenzoic acid | 1.707022655 | 0.771482205845 | 0.2716 | 0.56606761194 |
| L-lysine | 1.620653049 | 0.696575270053 | 0.02291 | 1.64001117184 |
| myristic acid | 1.55981747 | 0.641377214939 | 0.02097 | 1.67830254021 |
| L-valine | 0.658224217 | -0.60334898922 | 0.15306 | 0.8151428792 |
| glycerol 1-phosphate | 0.674547336 | -0.56800840843 | 0.0778 | 1.10902862597 |
| D-glucose-6-phosphate | 0.700061908 | -0.51444558751 | 0.14485 | 0.83908847323 |
| phosphoric acid | 0.700822646 | -0.51287869997 | 0.00029 | 3.53716349081 |
| 1,3-diaminopropane | 1.408028383 | 0.493676416136 | 0.1535 | 0.81388639501 |
| L-Isoleucine | 0.712095116 | -0.48985813646 | 0.43253 | 0.36398421045 |
| L-leucine | 0.71413946 | -0.48572225695 | 0.43585 | 0.36066383615 |
| alpha ketoglutaric acid | 1.398259694 | 0.483632332586 | 0.06935 | 1.15898183462 |
| 3,4-dihydroxybenzoic acid | 0.718337457 | -0.47726634916 | 0.07932 | 1.10063056719 |

| | | | | |
|---|---|---|---|---|
| eicosapentaenoic acid | 1.385784319 | 0.470702736593 | 0.09465 | 1.02386063491 |
| shikimic acid | 1.354435545 | 0.437691740268 | 0.02774 | 1.55694419424 |
| arbutin | 1.35263207 | 0.435769464676 | 0.36543 | 0.43719004264 |
| itaconic acid | 1.329064373 | 0.410410982449 | 0.06709 | 1.1733679178 |
| L-serine | 0.760197692 | -0.39555345009 | 0.54362 | 0.26470573825 |
| threonic acid | 0.762382341 | -0.39141339194 | 0.27565 | 0.55963753948 |
| glycine | 0.764881306 | -0.38669220736 | 0.39221 | 0.4064764337 |
| acetohydroxamic acid | 0.765323945 | -0.38585755594 | 0.19627 | 0.70715360482 |
| ethanolamine | 0.767641763 | -0.38149489294 | 0.39673 | 0.40150150473 |
| maleic acid | 0.774260836 | -0.36910842517 | 0.20224 | 0.69412776272 |
| heptanoic acid | 0.776897552 | -0.36420372855 | 0.18955 | 0.72226808208 |
| succinic acid | 0.779125933 | -0.36007155972 | 0.21024 | 0.67728569647 |
| fumaric acid | 1.258905941 | 0.332170495906 | 0.13316 | 0.87561951244 |
| Beta- alanine | 0.794427968 | -0.33201168071 | 0.32581 | 0.48703423341 |
| quinic acid | 1.251208227 | 0.323321903836 | 0.17863 | 0.74803432238 |
| sedoheptulose anhydride monohydrate | 1.249509798 | 0.321362214724 | 0.33072 | 0.48054266937 |
| L-threonine | 0.816332453 | -0.29277128246 | 0.65861 | 0.18137251519 |
| citric acid | 1.201425342 | 0.264747000071 | 0.04741 | 1.3241131307 |
| stearic acid | 0.844775722 | -0.24335972135 | 0.55074 | 0.25904973906 |
| maltose | 1.18182929 | 0.24102165991 | 0.3451 | 0.46205458061 |
| nicotinic acid | 1.181362371 | 0.240451564743 | 0.23078 | 0.63680190732 |
| gamma-aminobutyric acid (GABA) | 0.851915338 | -0.23121803002 | 0.51528 | 0.28795970441 |
| L-asparagine | 1.170552383 | 0.227189497631 | 0.45095 | 0.34587177644 |
| tyrosine | 1.169059811 | 0.225348741895 | 0.06933 | 1.159058722 |
| pantothenic acid | 1.159542161 | 0.213555277464 | 0.46042 | 0.33684671177 |
| 4-hydroxycinnamic acid | 1.159064225 | 0.212960509636 | 0.13901 | 0.85694511293 |
| lactose | 1.145186613 | 0.195582711083 | 0.59766 | 0.2235484997 |
| L-homoserine | 1.138147305 | 0.186687290796 | 0.48323 | 0.31584425005 |
| beta-gentiobiose | 1.133192993 | 0.180393585887 | 0.73523 | 0.13357937949 |
| elaidic acid | 1.133043704 | 0.180203510063 | 0.2226 | 0.65246927903 |
| 4-hydroxy-3-methoxybenzoic acid | 0.886978875 | -0.17302834963 | 0.65021 | 0.18694825851 |
| serotonin | 0.890344634 | -0.16756421396 | 0.63369 | 0.19812129843 |
| dehydroascorbic acid | 1.116405217 | 0.158860771145 | 0.41253 | 0.38454011497 |
| D-saccharic acid | 1.107771069 | 0.147659766594 | 0.387 | 0.41228880877 |
| Antiarol | 1.106314976 | 0.145762189511 | 0.29403 | 0.53161160004 |
| 2-hydroxypyridine | 1.105745764 | 0.145019715787 | 0.2383 | 0.62288124923 |
| Sucrose | 1.098960885 | 0.136140037387 | 0.38257 | 0.41729206675 |
| lauric acid | 1.094152015 | 0.129813191233 | 0.15102 | 0.82097784496 |
| myo-inositol | 1.086506334 | 0.119696584626 | 0.59037 | 0.22887736171 |
| glyoxylic acid | 1.086027119 | 0.119060129381 | 0.40397 | 0.39365158833 |
| 3-hydroxypyridine | 0.921368634 | -0.11814960902 | 0.14694 | 0.83286998454 |
| 5,6-dihydrouracil | 0.927342301 | -0.10882613016 | 0.34106 | 0.46717198725 |
| ribose | 0.933129061 | -0.09985146157 | 0.25791 | 0.58853725725 |
| L-mimosine | 1.069303621 | 0.096671553995 | 0.75496 | 0.12207432555 |
| aspartic acid | 0.937470124 | -0.09315538116 | 0.55762 | 0.25366217924 |
| capric acid | 1.066455679 | 0.092824010067 | 0.35652 | 0.44791680227 |
| benzene-1,2,4-triol | 1.065287975 | 0.091243481599 | 0.27307 | 0.5637336352 |
| D-mannose | 1.061810785 | 0.086526700453 | 0.68564 | 0.16390184998 |
| D-mannitol | 1.061342766 | 0.085890657552 | 0.68796 | 0.16243646793 |

| | | | | |
|---|---|---|---|---|
| D-glucose | 1.060952601 | 0.085360203542 | 0.68909 | 0.16172097674 |
| arachidic acid | 1.060232618 | 0.084380830507 | 0.35146 | 0.45412995264 |
| fructose | 1.049670458 | 0.06993646747 | 0.44356 | 0.3530448415 |
| beta-cyano-L-alanine | 1.048053073 | 0.067711775596 | 0.89747 | 0.04697892219 |
| 3,5-dimethoxy-4-hydroxycinnamic acid | 1.04743134 | 0.066855677413 | 0.4694 | 0.32846069822 |
| Pyruvic acid | 0.956934003 | -0.06350866475 | 0.44124 | 0.35532077255 |
| ferulic acid | 1.042527906 | 0.060086002375 | 0.61477 | 0.21128737037 |
| trans-aconitic acid | 0.963588707 | -0.05351060879 | 0.73873 | 0.13151252506 |
| oxalic acid | 0.963652839 | -0.05341459392 | 0.55547 | 0.25534017907 |
| adenosine | 1.035888476 | 0.050868691167 | 0.82402 | 0.08406288746 |
| L-alanine | 0.965431568 | -0.05075409305 | 0.58599 | 0.23211227024 |
| adenine | 0.968936973 | -0.04552527042 | 0.82465 | 0.08372865808 |
| caffeic acid | 1.029550643 | 0.042014796821 | 0.54632 | 0.26255222395 |
| spermidine | 0.97336149 | -0.03895239758 | 0.47709 | 0.3213987615 |
| benzoquinone | 0.976661823 | -0.03406899115 | 0.92969 | 0.03165993685 |
| L-glutamic acid | 0.977470467 | -0.03287498075 | 0.85996 | 0.06552124554 |
| benzoic acid | 1.022439005 | 0.032014779675 | 0.80275 | 0.09541937949 |
| allantoin | 0.978401953 | -0.03150081053 | 0.90415 | 0.04375885354 |
| 1,6-anhydro-glucose | 1.02184986 | 0.031183237044 | 0.09381 | 1.02774964121 |
| 3,4 dihydroxyphenylalanine (DOPA) | 1.021214722 | 0.030286240781 | 0.6885 | 0.1620940853 |
| pelargonic acid (nonanoic acid) | 1.020040449 | 0.028626362714 | 0.81037 | 0.09131405121 |
| palmitic acid | 0.980762799 | -0.02802383791 | 0.64788 | 0.18850800227 |
| D-malic acid | 0.987620309 | -0.01797159163 | 0.92928 | 0.03185379356 |
| glycerol | 1.004893017 | 0.00704191782 | 0.97856 | 0.00941073069 |
| L-glutamine | 0.997141533 | -0.00412980129 | 0.99098 | 0.0039347185 |
