## Supplementary figures and images for "The genome of stress tolerant crop wild relative *Paspalum vaginatum* leads to increased biomass productivity in the crop *Zea mays*"

### Supplementary note 6

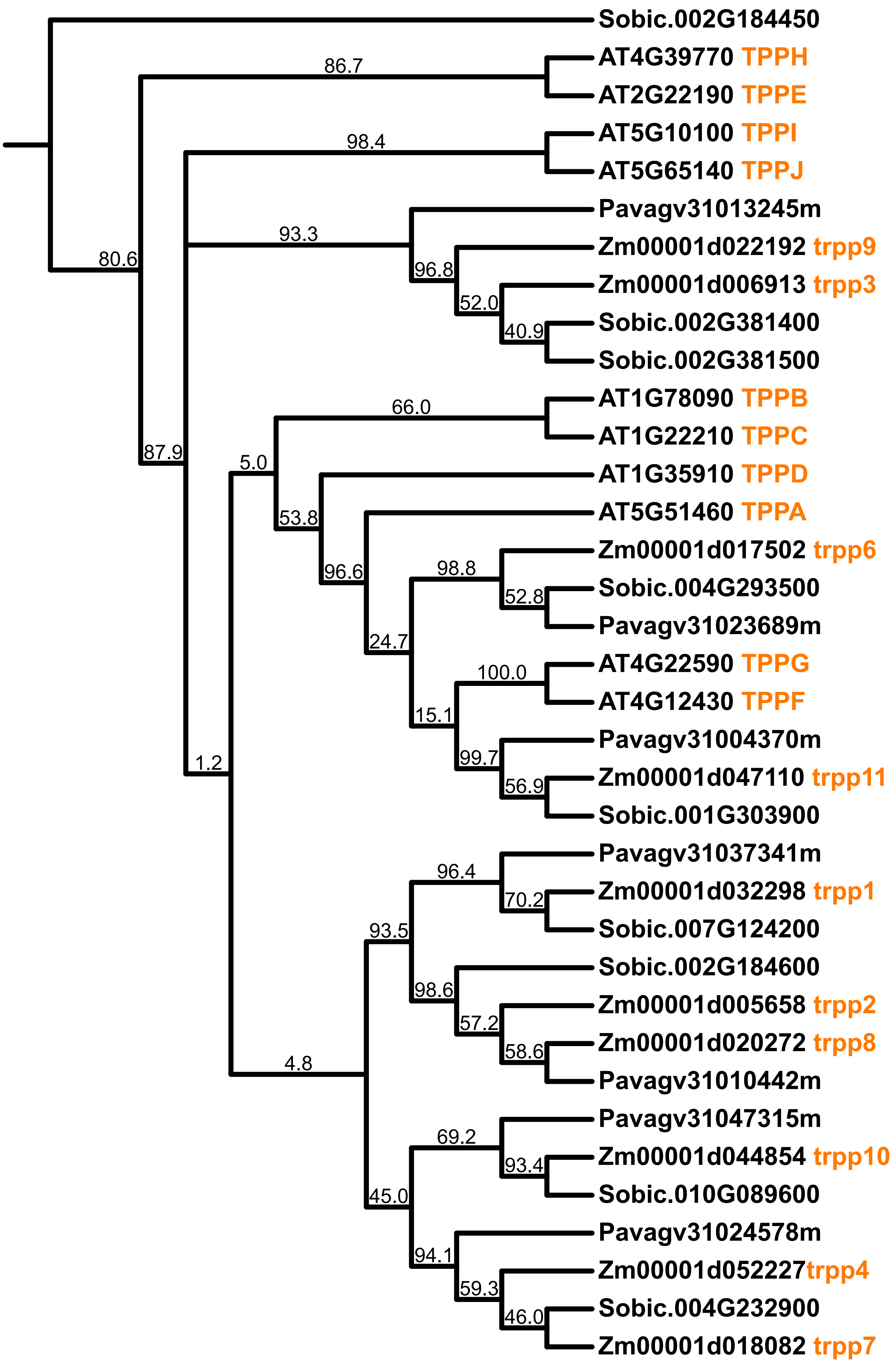

### Supplementary note 7

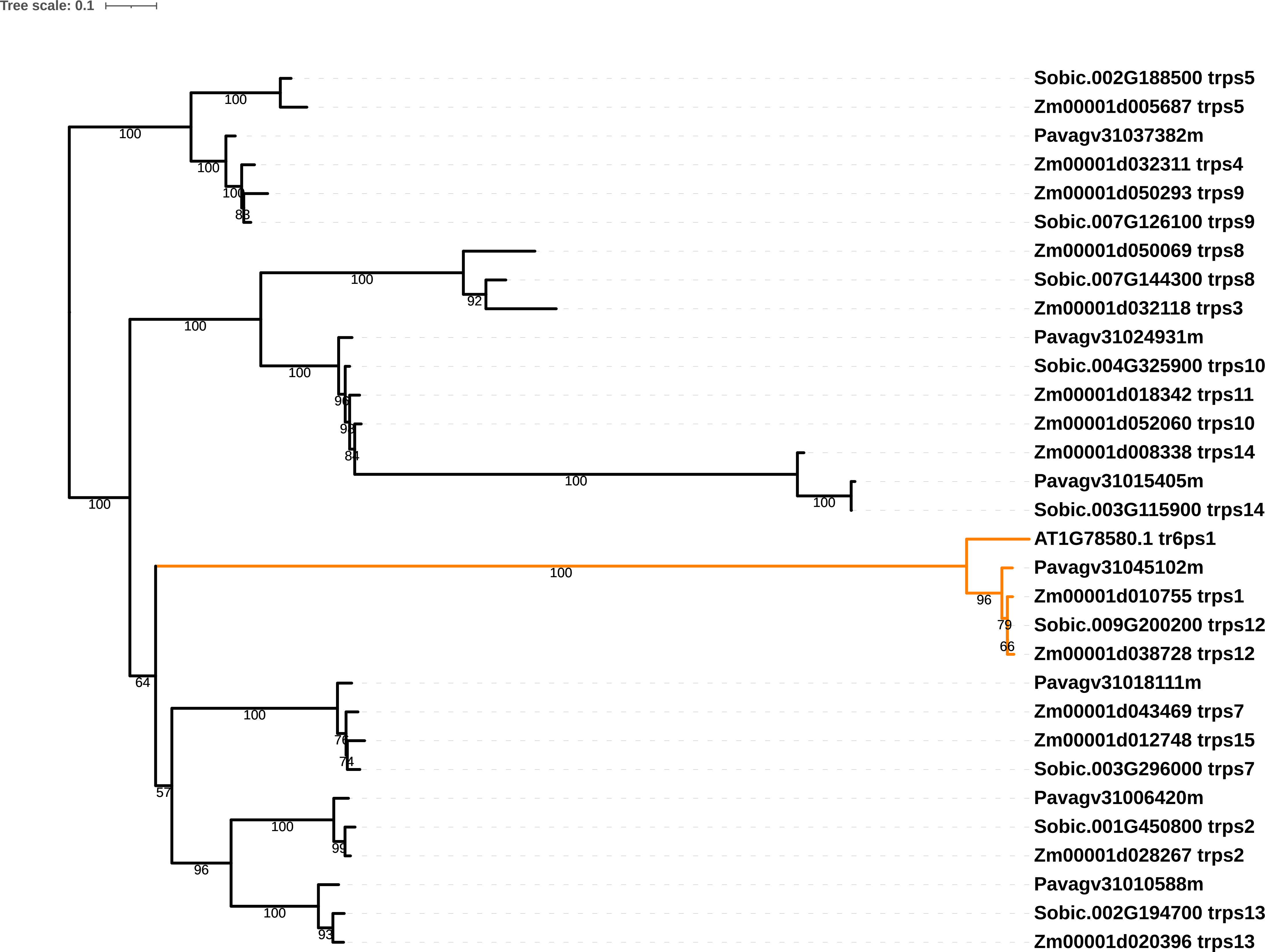
