## Supplementary note 8 for "The genome of stress tolerant crop wild relative *Paspalum vaginatum* leads to increased biomass productivity in the crop *Zea mays*"

| **Hoagland Full nutrient solution (1L)** | | | | | |
|---|---|---|---|---|---|
| Concentration | Chemical | wt(g)/L | For 250 mL Stock | mL/L for working stock | Final Concentration |
| 2M | KNO3 | 202.2 | 50.55 | 2.5 | 5 mM |
| 2M | $Ca(NO3)_2.4H_2O$ | 472.3 | 118.075 | 2.5 | 5 mM |
| 1M | KH2PO4(PH=6) | 136.09 | 34.0225 | 1 | 1 mM |
| 2M | MgSO4.7H2O | 492.94 | 123.3235 | 1 | 2 mM |
| 0.05M | Fe-EDTA* | 2.8 | 0.7 | 1 | 50 uM |
| | Micro Nutrients | | | 1 | |
| | | | Total | 9 | |
| ***0.05M Fe-EDTA (200ml):** 3.72g DETA in 70 mL ddiH2O mixed with 2.78 g FeSO4.7H2O in 70 mL ddiH2O and then volume up to 200 mL | | | | | |

| **Hoagland -Nitrogen nutrient solution (1L)** | | | | | |
|---|---|---|---|---|---|
| Concentration | Chemical | wt(g)/L | For 250 mL Stock | mL/L for working stock | Final Concentration |
| 0.5M | KSO4 | 87 | 21.75 | 10 | 5 mM |
| 2M | $CaCl2.2H_2O$ | 294.02 | 73.505 | 2.5 | 5 mM |
| 1M | KH2PO4(PH=6) | 136.09 | 34.0225 | 1 | 1 mM |
| 2M | MgSO4.7H2O | 492.94 | 123.3235 | 1 | 2 mM |
| 0.05M | Fe-EDTA* | 2.8 | 0.7 | 1 | 0.1 mM |
| | Micro Nutrients | | | 1 | |
| | | | Total | 16.5 | |

| **Hoagland -Phosphorus nutrient solution (1L)** | | | | | |
|---|---|---|---|---|---|
| Concentration | Chemical | wt(g)/L | For 250 mL Stock | mL/L for working stock | Final Concentration |
| 2M | KNO3 | 202.2 | 50.55 | 2.5 | 5 mM |
| 2M | $Ca(NO3)_2.4H_2O$ | 472.3 | 118.075 | 2.5 | 5 mM |
| 1M | KSO4 | 87 | 21.75 | 10 | 10 mM |
| 2M | MgSO4.7H2O | 492.94 | 123.3235 | 1 | 2 mM |
| 0.05M | Fe-EDTA* | 2.8 | 0.7 | 1 | 0.1 mM |
| | Micro Nutrients | | | 1 | |
| | | | Total | 18 | |

| **Micro Nutrients (1L)** | | |
|---|---|---|
| Final Concentration | Chemical | wt(g)/L |
| 0.046M | h3bo3 (Boric Acid) | 2.86 |
| 0.009M | MnCl2.4H2O (Manganese Chloride) | 1.81 |
| 7.5X10-4M | ZnSO4.7H2O (Zinc Sulphate) | 0.22 |
| 3.2X10-4M | CuSO4 (Copper Sulphate) | 0.08 |
| 1.11X10-4M | H2MoO4.4H2O (Molybdic Acid 85%) | 0.02 |

| Concentration | Chemical | wt(g)/L | For 250 mL Stock |
|---|---|---|---|
| 2M | KNO3 | 202.2 | 50.55 |
| 2M | $Ca(NO3)_2.4H_2O$ | 472.3 | 118.075 |
| 1M | KH2PO4(PH=6) | 136.09 | 34.0225 |
| 2M | MgSO4.7H2O | 492.94 | 123.3235 |
| 0.05M | Fe-EDTA* | 2.8 | 0.7 |
| 0.5M | KNO3 | 202.2 | 50.55 |
| 2M | $CaCl2.2H_2O$ | 294.02 | 73.505 |
